# Low-Dimensional and Optimised Representations of High-Level Information in the Expert Brain

**DOI:** 10.1101/2025.11.12.688012

**Authors:** Andrea I. Costantino, Artem Platonov, Felipe Fontana Vieira, Emily Van Hove, Merim Bilalić, Hans Op de Beeck

## Abstract

What transforms a novice into an expert? Decades of research show that expertise relies on domain-specific knowledge, but a neural account of this transformation has remained fragmentary: we lack an understanding of what information expert representations encode, how they are structured for efficient use, and where in the brain they reside. Using chess as a model system, we combine neuroimaging with multivariate pattern analysis to reveal three principles of the expert brain. Expertise drives a shift in representational content, from surface visual features to high-level, relational information. It is accompanied by a structural change to low-dimensional, optimised representation: codes become more compact and better organised, yet retain the detail needed for precise evaluation. Finally, the representational load shifts from sensory-specific cortices to domain-general frontoparietal networks. The expert brain packs more into less, concentrating richer knowledge into fewer, better-organised representations that support the rapid, flexible decisions of mastery.

## Introduction

What makes a novice into an expert? This longstanding question sits at the intersection of neuroscience, cognitive science, and artificial intelligence ^1–4^. Across diverse domains such as mathematics ^5,6^, language ^7^, motor skills ^8–10^, and board games ^11–13^, extensive experience equips individuals with domain-specific knowledge, which serves as the foundation for fast, accurate, and flexible decision-making ^14^. Decades of cognitive research have established how such knowledge supports expert performance at the behavioural and computational levels. Yet, we still lack a precise account of its neural basis: specifically, ***what*** information expert representations encode (the *content*), ***how*** that information is organised for efficient readout (the *structure*), and ***where*** in the brain such representations are expressed (the neural *substrate*).

Research on expertise has revealed that expert performance rests much more on knowledge than on raw general ability ^15,16^. Years of focused exposure tune perception and memory to a domain’s recurring structure ^3,17^. Co-occurring features are grouped into chunks ^11^ and richer templates ^18^ that bind surface cues to deeper relations and typical courses of action. When familiar constellations appear, they trigger these stored structures, which rapidly unpack the relevant details and predictions ^11,17,18^, functioning like an extension of working memory ^19^. This enables experts to quickly grasp a situation’s underlying structure while simultaneously preparing appropriate responses. Novices, in contrast, organise by appearances and get stuck on particulars. Expert cognition is therefore qualitatively different: recognition-driven and relational, powered by knowledge that supports both a quick recognition and fine-grained analysis when needed ^20,21^.

Neuroimaging studies confirm that expertise involves a functional reorganisation of neural resources toward domain-specific and memory-based processing, which enable access to knowledge for fast, accurate decisions ^20,22^. Across domains, these systems are engaged more strongly (and often bilaterally) in experts than in novices ^21^. The precise loci vary with stimulus type, mapping where domain knowledge is stored and read out. In chess, experts preferentially recruit lateral temporal and parietal regions for object identities and functions but invoke scene/navigation systems for pattern recognition ^23–27^. Radiological expertise, with its holistic processing at its core, is characterised by ventral occipito-temporal selectivity ^28,29^. In higher-level calculation, professional mathematicians engage a bilateral frontoparietal-temporal network ^5,30^, while mental calculators draw on episodic memory components in the temporal lobe ^31^. When computation is externalised, as with the abacus, experts rely on visuospatial and premotor simulation circuits related to instrument manipulation ^32^. Finally, in motor expertise, a premotor-parietal action-observation network (AON) allows for efficient anticipation from kinematic cues ^8,22^.

These studies mostly rely on univariate contrasts. They highlight regions with increased activation but do not reveal the structure or content of the encoded information. We know which regions are more active in experts, but not how they represent domain-specific knowledge. To close that gap, we move from localisation to representational geometry, investigating how domain knowledge is represented. Addressing these questions requires analytical methods that go beyond region-wise activation.

Multivariate pattern analysis (MVPA)–including decoding and representational similarity analysis (RSA)–offers a framework for probing the informational content and structure of neural activity ^33^. These methods allow us to quantify not just whether two conditions elicit different responses, but also how stimuli are internally represented and organised in representational space ^34–36^. While MVPA methods have been used with success in vision ^35,37,38^, language ^39,40^, and memory ^41,42^, they remain underused in research on expertise.

Here, we apply these methods to the domain of chess, a model system in cognitive science and artificial intelligence ^43,44^, with objective skill rating (Elo) and well-controlled stimuli that jointly engage low-level perceptual processing (board layout, piece identity) and high-level cognitive operations (strategy, planning, goal evaluation). We combine behavioural measures with univariate fMRI, MVPA (decoding/RSA), and manifold-based dimensionality analyses (Fig. 1). We then ask not only where experts and novices show different average activations, but also **what** information expert representations encode (content), **how** that information is organised for efficient readout (structure), and **where** in the brain these codes are expressed (neural substrate). This representational approach moves beyond localisation or activation-based accounts by characterising the internal geometry of neural codes rather than their magnitude alone.

**Fig. 1.**
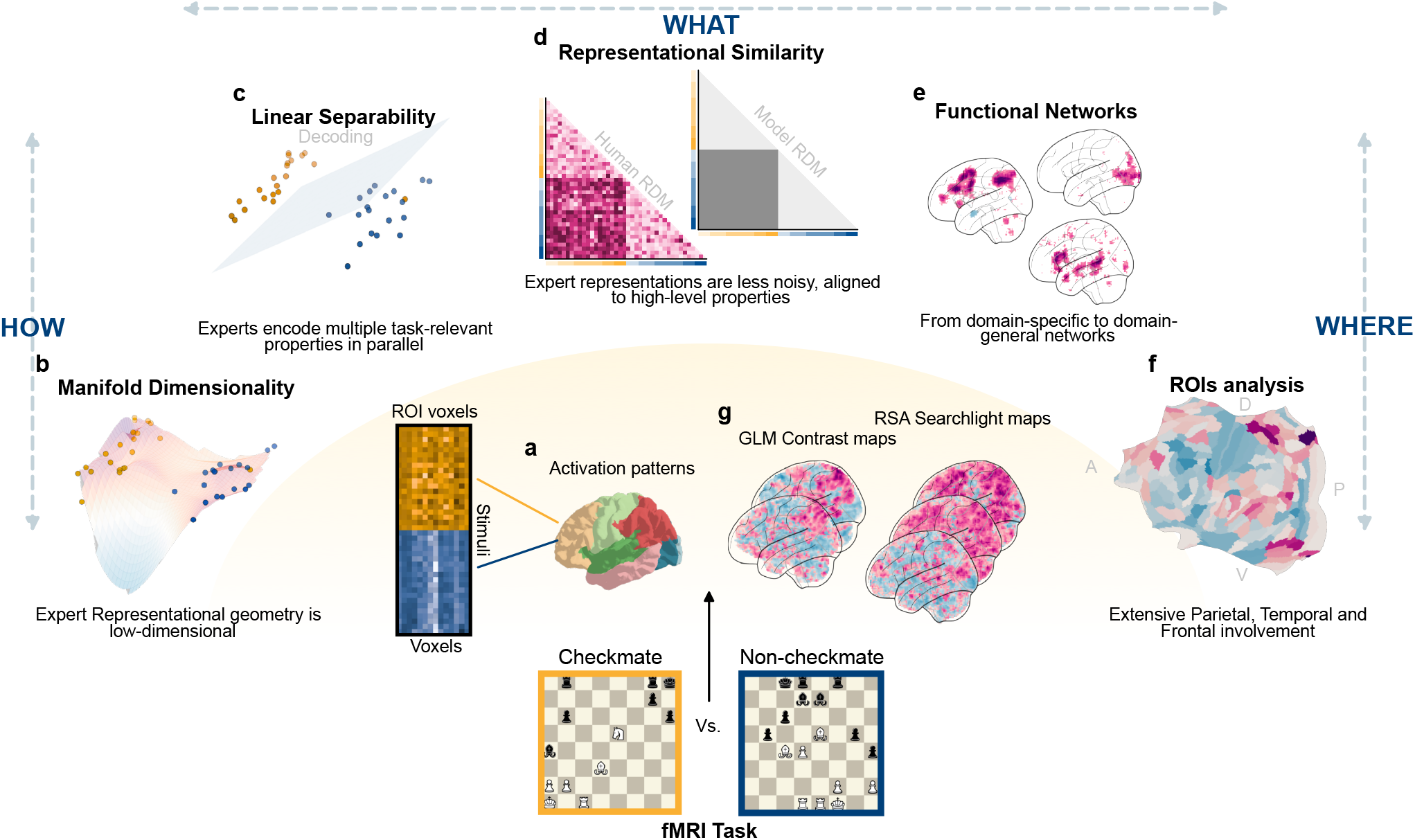
Overview of questions, analyses, and results. Chessboard stimuli in the fMRI task elicit multivoxel patterns, analysed to investigate *what* is represented, *how* it is structured, and *where* it is expressed. Activation patterns are parcellated into 22 bilateral cortical ROI groups from the Glasser atlas (**a**; colours indicate cortical families, e.g., primary visual, frontal). Within each ROI, we perform representational similarity analysis (RSA) to test the correspondence between theoretical model representational dissimilarity matrices (RDMs) and neural RDMs (**d**; *what*), estimate manifold dimensionality via participation ratio (**b**), and test linear separability of stimulus categories in the multivoxel space (**c**; *how*). At the whole-brain level, group-level general linear model (GLM) contrasts and RSA searchlight maps (**g**) are correlated with meta-analytic Neurosynth association maps for seven cognitive terms (**e**; three shown) to characterise the functional identity of expertise effects (*where*). The surface map (**f**) depicts the unconstrained colourmap from the Visual Similarity RSA searchlight; direction descriptors at the corners (A: anterior, P: posterior, D: dorsal, V: ventral) indicate brain orientation (see Supplementary Fig. 15 for the full statistical maps with labelled ROI borders).

Crucially, the answers to these questions are not predetermined by existing theory or data. Expert knowledge could either expand or compress neural codes: richer distinctions among board states could yield higher-dimensional representations, while abstraction over shared relational structure could collapse them onto fewer dimensions. Chunk and template theories describe content enrichment without prescribing a specific geometry ^11,17,18^.. The representational locus could remain in domain-specific cortex, as prior neuroimaging has consistently localised expertise effects to category-selective regions (fusiform, temporal, parahippocampal) ^20,21,45^. And if expert representations do compress, that compression could sacrifice fine-grained detail for speed, producing fast but coarse evaluation. Whereas classical expertise theories specify the shift from perceptual to relational content, the present study asks how that shift is neurally realised: whether expertise compresses or expands neural codes, whether its locus shifts from domain-specific to domain-general systems, and whether compression preserves or discards the detail that expert performance requires.

Using this multilevel framework, we reveal three principles. First, expert representations prioritise relational, goal-relevant information over surface features (what/content). Second, they are more compact but still easier to distinguish because they preserve fine detail (how/structure). Third, expert representations are expressed most strongly across domain-general frontoparietal control systems (where/substrate). These properties provide a representational account of expertise: compressed, separable codes that retain decision-critical details and enable rapid, flexible performance. By clarifying how rich knowledge is deployed rapidly while retaining access to task-relevant detail, this pattern bridges classical cognitive theory with neural and computational theories of learning.

## Results

We compared twenty long-time rated tournament chess players (experts) with twenty chess-literate hobby players (novices). Both experts and novices performed a domain-specific 1-back task during fMRI: on each trial they judged which of the two most recent boards was more strategically advantageous for White, pressing one of two buttons (two-alternative forced choice; Fig. 2D for procedure, for more details see Methods).

**Fig. 2.**
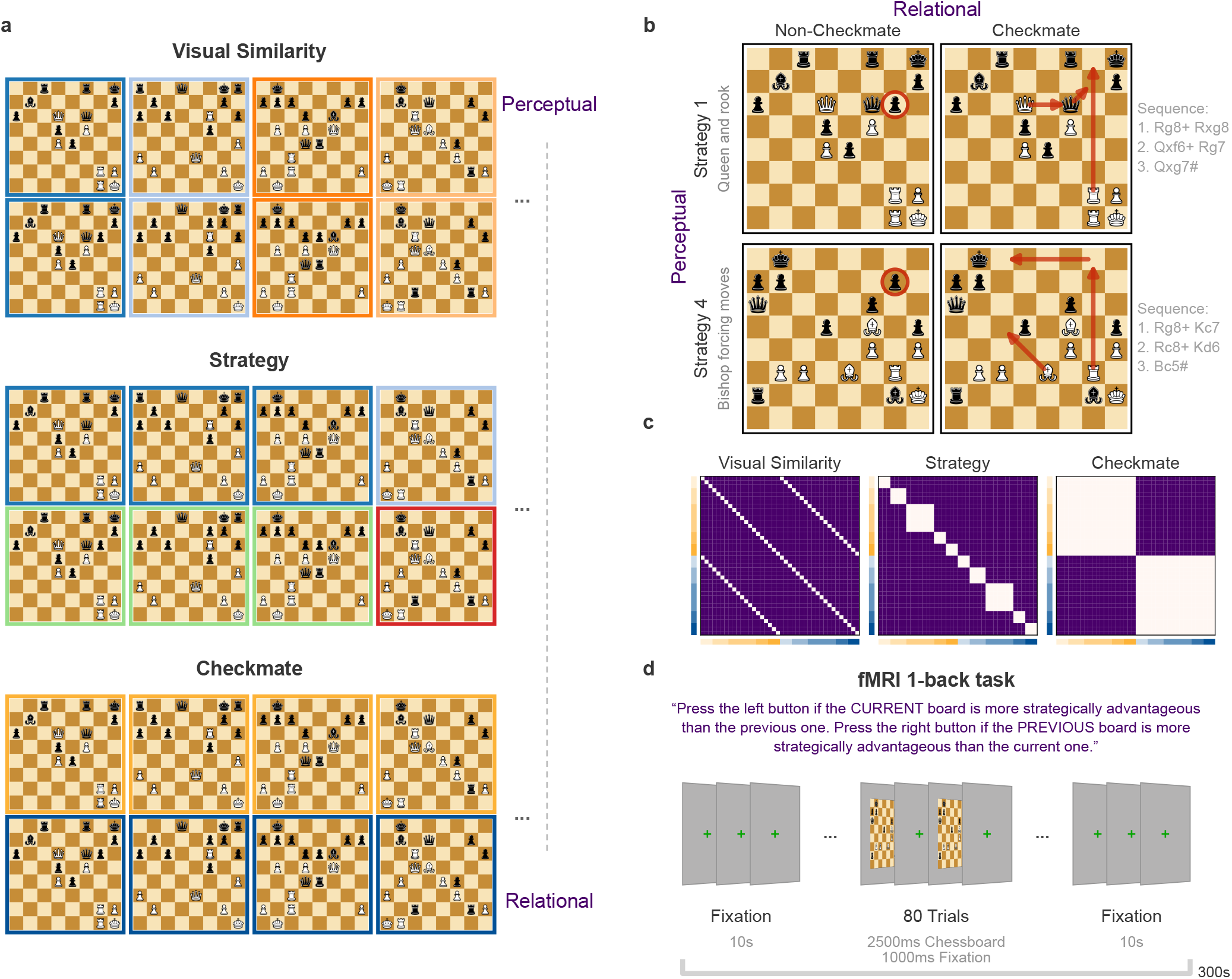
Stimulus design and predicted representational structures. **a**. Visual depiction of the stimuli. Each row reflects a different clustering logic, corresponding to a distinct predicted representational geometry. Coloured borders indicate the stimulus label under each categorisation: Checkmate vs. non-checkmate boards (resp. orange vs. dark-blue); Strategy 1 vs. 2 vs. 6 vs. 7 (dark blue, light blue, green, red) etc.; Visual pair 1 vs. 2 vs. 3 (dark blue, light blue, dark orange) etc. **b**. Example stimuli from the dataset. Columns show visually similar pairs from the Checkmate and Non-Checkmate conditions; rows reflect different strategies. Red circles highlight the pieces repositioned in Non-Checkmate boards to disable the mating line, and arrows indicate the checkmate sequence in Checkmate boards. **c**. Theoretical Representational Dissimilarity Matrices (RDMs) derived from categorical labels. Lighter values indicate greater predicted similarity (i.e., same-label pairs). **d**. Main experimental task performed during scanning. Each run included 80 trials (each of the 40 boards presented twice in randomised order), each presented for 2.5 seconds followed by a 1-second fixation. Participants performed a two-alternative forced-choice 1-back evaluation: on every trial, they judged which of the two most recent boards was more strategically advantageous for White and pressed the corresponding button (mapping counterbalanced across runs).

The stimulus set was built so that the same boards could be grouped under three complementary labels (Fig. 2A): (i) perceptual (visually matched pairs that share surface layout and piece identities), (ii) strategy (recurring relational motifs such as queen-rook ladder mates or bishop-forcing patterns), and (iii) checkmate (checkmate vs. non-checkmate). By “relational” we mean properties defined over relations among pieces and legal move constraints that realise a goal (e.g., attack/defence structure and forced move sequences that constitute a mate); “strategy” refers to the specific relational motif implementing that goal. In contrast, “perceptual” similarity refers to surface layout/identity irrespective of whether the legal relations yield a solution. One could think of perceptual as “the board looks the same,” while strategy/relational as “the same plan and constraints apply,” even if pieces are shifted to nearby squares (e.g., a queen–rook ladder plan still works one file over).

To dissociate these sources of similarity, each checkmate had a perceptually matched non-checkmate counterpart created by a minimal intervention, typically repositioning one or two pieces, that disables the mating line (Fig. 2B, red circles). Checkmate/non-checkmate pairs are perceptually alike but relationally distinct, whereas boards within the same strategy share a relational motif regardless of surface details. In other words, strategy pairs preserve the same attack–defence relations by design, while checkmate pairs deliberately break exactly one critical relation (e.g., an extra flight square/defender) to flip checkmate to non-checkmate while holding appearance constant.

These categorical labels yield theoretical representational dissimilarity matrices (RDMs) that specify our model predictions for behavioural and neural geometry (Fig. 2C): low dissimilarity for same-label pairs within each model (perceptual, strategy, checkmate) and higher dissimilarity otherwise. Variance partitioning confirms that the three RDMs are largely orthogonal, each capturing a distinct dimension of board similarity (Supplementary Sec. S1). We use these models to test what information is represented via behavioural RSA and fMRI RSA/decoding.

Our analyses follow three questions (Fig. 1). We first ask **what** information is represented by aligning behavioural RDMs (from 1-back value preferences) and neural RDMs (multivoxel pattern distances) with model RDMs for perceptual, strategy, and checkmate structure (Fig. 2C). We then ask **how** these representations are organised– quantifying effective dimensionality with the participation ratio and testing readout (linear separability) with multivariate decoding. Finally, we ask **where** these signatures reside by combining ROI and searchlight analyses with meta-analytic network and term maps to situate effects within broader functional systems.

### What Is Represented: Perceptual to Relational Content

#### Behavioural representational geometry

We first characterise the representational geometry of participants’ choices during the fMRI task by conducting a behavioural representational similarity analysis (RSA). For that purpose, we used participants’ responses in the 1-back chess evaluation task, in which they judged which of the two most recent boards was more advantageous for White. These choices implicitly reflect the dimensions guiding participants’ internal representations of chessboard value. We computed behavioural RDMs for each group, derived from pairwise stimulus choices (see Section 4.7.1). We then tested whether these behavioural RDMs were significantly correlated with model RDMs reflecting three candidate dimensions of board similarity: *visual similarity, checkmate status*, and *strategy* (Fig. 2).

Expert directional preference matrices and the derived behavioural RDMs exhibit a clear block structure by Checkmate, with sub-structure by Strategy (Fig. 3A; orange vs. blue axis labels; saturation indicates strategy). The selection frequencies further echo this structure: experts show systematic preferences for checkmates and particular strategic motifs rather than flat, uniform counts (Fig. 3C). The MDS embedding mirrors these patterns: expert configurations show partial separation of Checkmate (orange) from Non-checkmate (blue) boards, indicating an organised preference geometry (Fig. 3B).

**Fig. 3.**
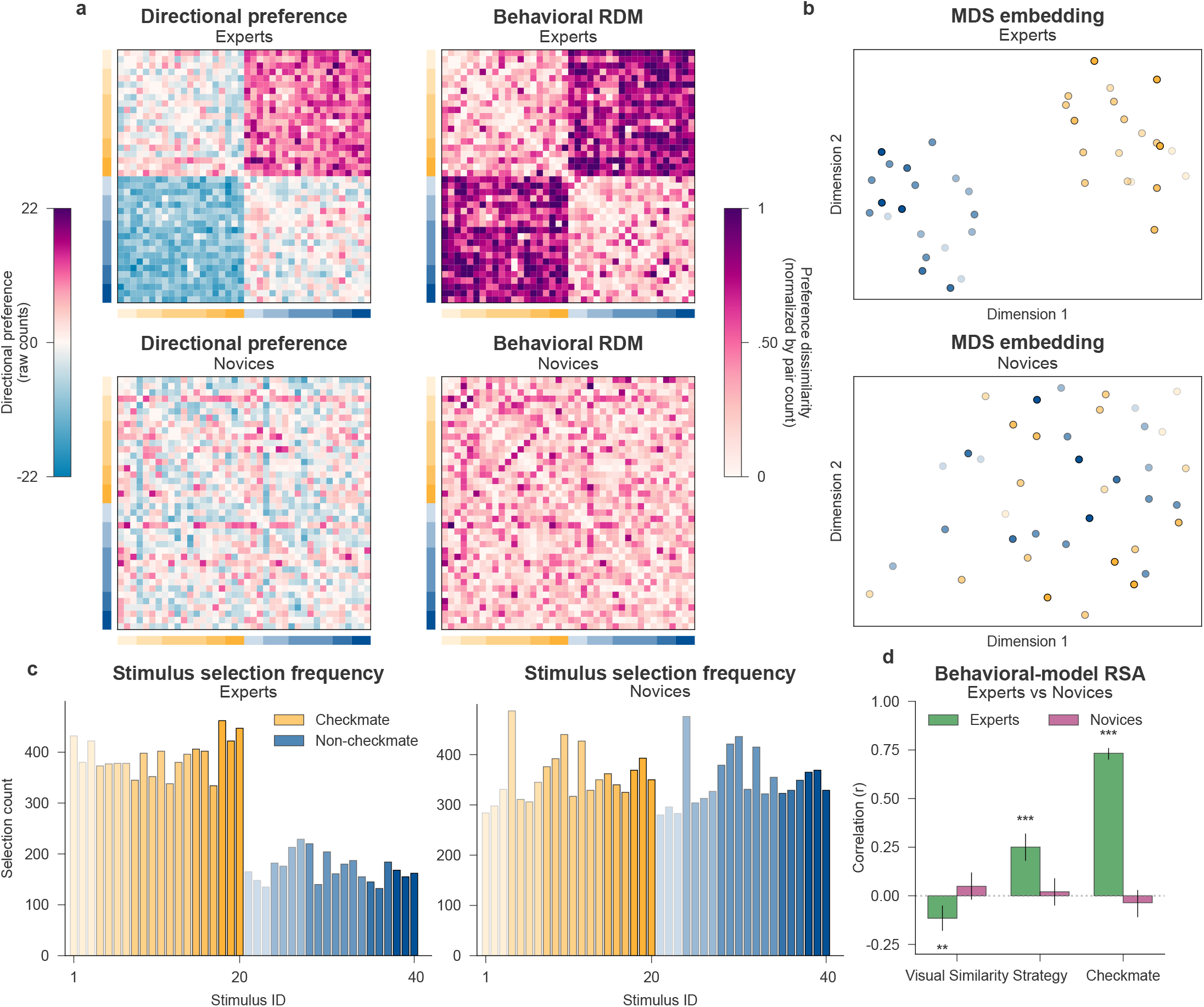
Behavioural representational analysis of chessboard preferences. **a**. Directional preference matrices (raw signed counts) and behavioural Representational Dissimilarity Matrices (RDMs; count-normalised) for Experts and Novices. The directional matrices show the net number of times one stimulus was preferred over another, pooled across all participants in each group: positive values (row *>* column) and negative values (column *>* row) summarise the group-level preference structure in raw counts. Because the 1-back design samples some pairs more often than others, those raw counts overestimate dissimilarity for more-frequently-sampled pairs; the behavioural RDMs therefore take the absolute signed difference and normalise each cell by the per-pair total (see Methods), and these normalised RDMs are what we use in all downstream behavioural RSA analyses. Orange and blue axis labels mark Checkmate and Non-checkmate stimuli, respectively; colour saturation reflects strategic labels. **b**. Two-dimensional multi-dimensional scaling (MDS) projection of the behavioural RDMs (`sklearn.manifold.MDS,` default Euclidean dissimilarity, `n_components=2`), illustrating the representational geometry of stimulus preferences. **c**. Choice frequency per stimulus, aggregated across participants. Bars indicate how often each chessboard was preferred in pairwise trials. Stimuli are colour-coded by checkmate status (orange = Checkmate; blue = Non-checkmate), with saturation indicating strategy label. Stimuli 1-20 are Checkmate boards, and 21-40 are their visually matched Non-checkmate counterparts. **d**. Correlation between count-normalised group-level behavioural RDMs and three categorical model RDMs (Checkmate, Strategy, Visual Similarity). Bars show Pearson correlations between the group-level behavioural RDM and each model RDM, computed on the lower triangle of the 40 × 40 RDM (*n* = 780 unique stimulus pairs per group; group RDMs aggregated across *n* = 19 Experts and *n* = 20 Novices, independent participants). Error bars are 95% confidence intervals from a 10,000-sample percentile bootstrap of stimulus pairs (`pingouin.corr`). *p*-values are two-sided and FDR-corrected (Benjamini–Hochberg) across the three models within each group at *α* = 0.05: ^∗^ *p*_FDR_ *<* .05, ^∗∗^ *p*_FDR_ *<* .01, ^∗∗∗^ *p*_FDR_ *<* .001. Exact *p*-values are listed in Table 2. Source data are provided as a Source Data file.

In contrast, novices’ preferences are not significantly predicted by any of the tested model dimensions. Novices’ directional matrices and behavioural RDMs lack the coherent block structure observed in experts (Fig. 3A), and selection counts are flatter and less differentiated by checkmate status across stimuli (Fig. 3C). The MDS for novices is diffuse, with no clear separation between checkmates and their matched non-checkmates (Fig. 3B).

Quantitative model fits (Fig. 3D, Table 2) converge with this picture (all *p*s are FDR-corrected; *n* = 780 lower-triangle RDM pairs): experts’ behavioural RDMs correlate strongly with Checkmate (Pearson’s *r* = 0.733, 95% CI [0.700, 0.760], *p*_FDR_ *<* .001) and reliably with Strategy (*r* = 0.251, 95% CI [0.180, 0.320], *p*_FDR_ *<* .001). This is consistent with the hypothesis that experts abstract over relational motifs and tactics. In contrast, visual similarity shows a small negative correlation with behaviour, yielding a weak but statistically significant effect (*r* = −0.116, 95% CI [−0.180, −0.050], *p*_FDR_ = .001). The negative association suggests that experts are less likely to rely on superficial visual features when making their choices–potentially suppressing irrelevant information. For novices, none of the model correlations are significant (Fig. 3D, Table 2), aligning with their unstructured preference matrices and diffuse MDS geometry.

These results indicate that, unlike experts, novices did not exhibit structured representations aligned with the relational or goal-relevant properties of the boards. However, their lack of significant model correlations at the full pairwise level should not be taken to imply an absence of structure altogether. Rather, it suggests that whatever structure does guide novice choices is not aligned with chess-relevant relational dimensions.

This interpretation is supported by a split-half reliability analysis (Supplementary Sec. S2), which reveals significantly higher within-group consistency among experts compared to novices. Further diagnostic analyses (Supplementary Sec. S2) confirm that novices were actively engaged with the task and applied internally consistent evaluation strategies, but these strategies were organised along perceptual rather than relational dimensions: novices preferred boards based on visual features shared within matched pairs, regardless of checkmate status. A systematic feature analysis confirmed that novice preferences were driven by visual complexity (officer count, image entropy, edge density), while expert preferences were predicted exclusively by checkmate status (Supplementary Sec. S2). Novices thus shared a common perceptual evaluation strategy, but because this strategy relies on multiple continuous visual features rather than a single categorical distinction, it did not yield stable fine-grained pairwise orderings across the full stimulus set. In contrast, expert preferences reflect a more abstract and relational understanding of the stimuli. Expert preferences are governed by relational features that determines each pairwise comparison consistently, producing reliable structure at both the marginal and pairwise levels. This behavioural asymmetry foreshadows the neural dimensionality finding: expert codes are compressed onto fewer dimensions, while novice codes are spread across a higher-dimensional perceptual feature space. A perceptual-to-relational feature gradient analysis across the 40 boards further quantified this dissociation: expert board preferences are uniquely predicted by checkmate status (partial *ρ* = 0.94, 95% CI [0.91, 0.99], *p*_FDR_ *<* .001) while novice preferences are largely predicted by officer count (partial *ρ* = 0.55, 95% CI [0.27, 0.77], *p*_FDR_ = .006), with 92.7% vs. 3.4% of explained variance attributable to the strategic-relational feature block, respectively (Supplementary Fig. 3).

#### Neural representational geometry

We further performed RSA on multivoxel activation patterns to assess how the brain represents chess-relevant information across regions and groups. RSA quantifies the degree to which the dissimilarity structure of neural activation patterns aligns with predicted categorical models, emphasizing the dominant representational dimensions that shape the geometry of neural space. All reported *p*-values are FDR-corrected.

Fig. 4 shows the RSA results comparing Experts and Novices across regions of interest (ROIs) that span the cerebral cortex (see Supplementary Table 4 for full values). For the *Visual Similarity* model—capturing low-level perceptual resemblance—group differences were small across ROIs (all |Δ*r* |*<* .03; e.g., Primary Visual Δ*r* = −.020, *t*(35.7) = −0.96, *p*_FDR_ = .922, Cohen’s *d* = −0.30, 95% CI [−.062,.022]; Ventral Visual Δ*r* = −.007, *t*(37.9) = −0.43, *p*_FDR_ = .922, Cohen’s *d* = −0.14, 95% CI [−.042,.028]; Dorsal Stream Visual Δ*r* = .009, *t*(37.7) = 0.46, *p*_FDR_ = .922, Cohen’s *d* = 0.15, 95% CI [−.029,.046]). This pattern indicates broadly similar representational geometries for low-level visual features in both groups.

**Fig. 4.**
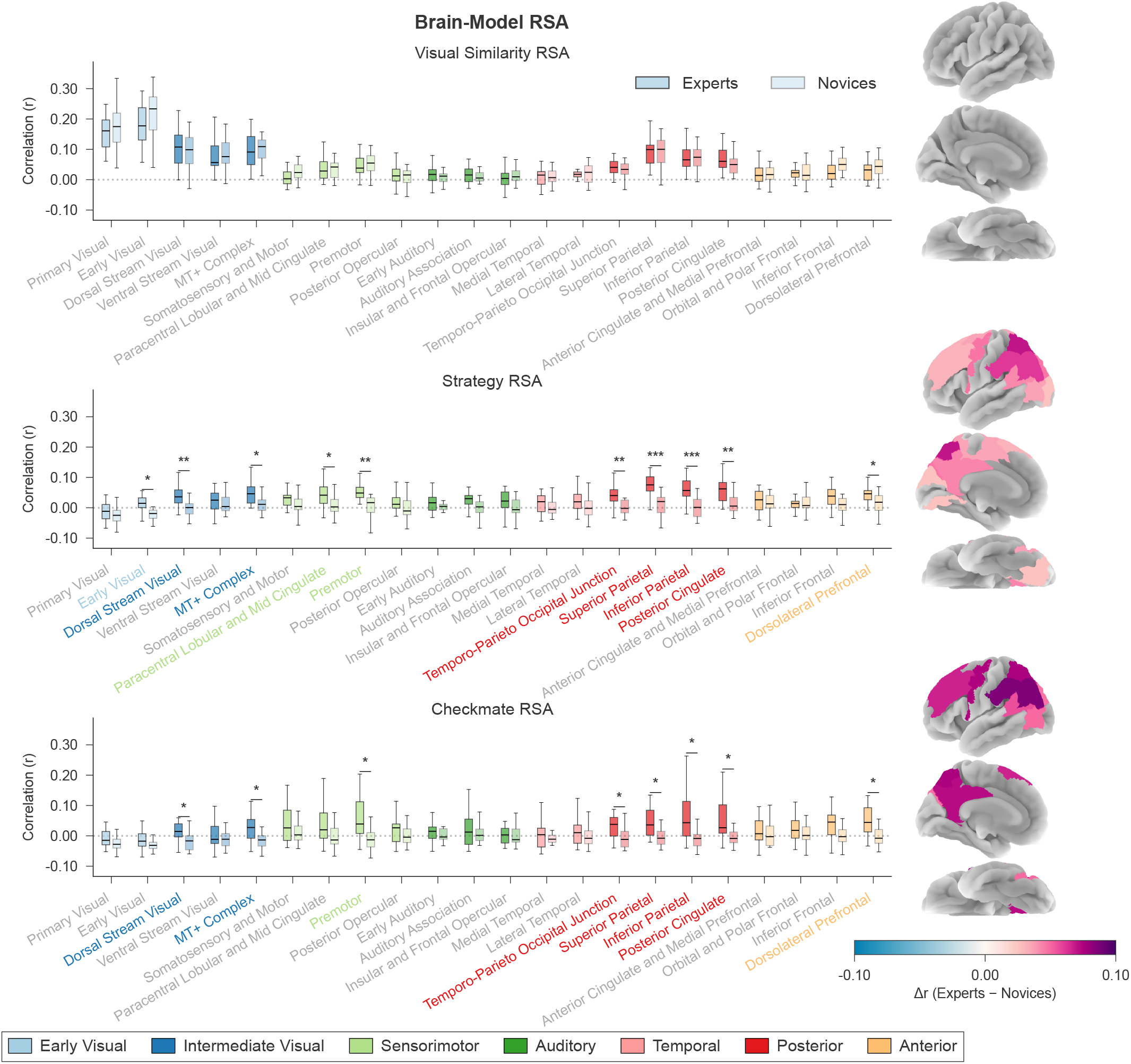
ROI-based RSA results. Representational similarity analysis between theoretical model RDMs and brain RDMs derived from Glasser coarse ROIs. Box plots show the distribution of per-subject Pearson correlations between model RDMs and neural RDMs for Experts (*n* = 20 independent participants) and Novices (*n* = 20 independent participants) across 22 ROIs. Boxes span the interquartile range (IQR; 25th–75th percentile), centre lines mark the median, and whiskers extend to the furthest data point within 1.5 × IQR of the box edges. Box colours group ROIs into cortical families (key at bottom). Stars indicate Experts vs. Novices group differences from a two-sided Welch’s *t*-test (`scipy.stats.ttest_ind` with `equal_var=False`), FDR-corrected (Benjamini–Hochberg) across the 22 ROIs separately within each model: ^∗^ *p*_FDR_ *<* .05, ^∗∗^ *p*_FDR_ *<* .01, ^∗∗∗^ *p*_FDR_ *<* .001. Within-group correlations were assessed with a one-sample one-tailed *t*-test against zero (`alternative=‘greater’`). Group difference maps (Experts − Novices) projected onto the pial surface are shown next to each plot; only ROIs with FDR-significant group differences are coloured. Per-ROI *t*-statistics, Welch–Satterthwaite degrees of freedom, mean differences, 95% confidence intervals, Cohen’s *d*, raw and FDR-corrected *p*-values for all 22 ROIs and three models are reported in Supplementary Table 4. Source data are provided as a Source Data file.

In contrast, the *Strategy* model revealed markedly larger expert–novice differences across a distributed network (Fig. 4, middle). Effect sizes were strongest in dorsal/posterior regions, including Superior Parietal (Δ*r* = .070, *t*(36.6) = 5.42, *p*_FDR_ *<* .001, Cohen’s *d* = 1.71, 95% CI [.044,.096]), Inferior Parietal (Δ*r* = .062, *t*(36.2) = 5.14, *p*_FDR_ *<* .001, Cohen’s *d* = 1.62, 95% CI [.037,.086]), Posterior Cingulate (Δ*r* = .043, *t*(37.2) = 3.58, *p*_FDR_ = .004, Cohen’s *d* = 1.13, 95% CI [.019,.068]), and the Temporo-Parieto-Occipital junction (Δ*r* = .043, *t*(34.9) = 3.46, *p*_FDR_ = .005, Cohen’s *d* = 1.09, 95% CI [.018,.068]). Robust effects also appeared in Premotor cortex (Δ*r* = .047, *t*(36.8) = 3.73, *p*_FDR_ = .004, Cohen’s *d* = 1.18, 95% CI [.022,.073]), the Dorsal Visual Stream (Δ*r* = .046, *t*(32.1) = 4.20, *p*_FDR_ = .001, Cohen’s *d* = 1.33, 95% CI [.024,.069]), MT+ / neighbouring visual areas (Δ*r* = .037, *t*(37.7) = 2.88, *p*_FDR_ = .021, Cohen’s *d* = 0.91, 95% CI [.011,.062]), and dorsolateral prefrontal cortex (Δ*r* = .031, *t*(38.0) = 2.65, *p*_FDR_ = .026, Cohen’s *d* = 0.84, 95% CI [.007,.055]). The difference maps (right panel) highlight bilateral dorsal patches consistent with stronger abstraction- and tactic-related structure in expert representations.

A similar pattern was observed for the *Checkmate vs. Non-checkmate* model, with larger effects overall: Superior Parietal (Δ*r* = .076, *t*(21.9) = 3.38, *p*_FDR_ = .015, Cohen’s *d* = 1.07, 95% CI [.029,.122]), Inferior Parietal (Δ*r* = .086, *t*(22.2) = 3.51, *p*_FDR_ = .014, Cohen’s *d* = 1.11, 95% CI [.035,.137]), Posterior Cingulate (Δ*r* = .072, *t*(22.1) = 2.85, *p*_FDR_ = .031, Cohen’s *d* = 0.90, 95% CI [.020,.125]), temporo-parieto-occipital junction (Δ*r* = .056, *t*(25.9) = 3.03, *p*_FDR_ = .024, Cohen’s *d* = 0.96, 95% CI [.018,.094]), MT+ (Δ*r* = .050, *t*(27.2) = 3.47, *p*_FDR_ = .014, Cohen’s *d* = 1.10, 95% CI [.020,.079]), Premotor cortex (Δ*r* = .074, *t*(26.5) = 3.50, *p*_FDR_ = .014, Cohen’s *d* = 1.11, 95% CI [.031,.118]), Dorsal Visual Stream (Δ*r* = .047, *t*(25.8) = 2.55, *p*_FDR_ = .047, Cohen’s *d* = 0.81, 95% CI [.009,.085]), and dorsolateral prefrontal cortex (Δ*r* = .066, *t*(23.5) = 2.81, *p*_FDR_ = .031, Cohen’s *d* = 0.89, 95% CI [.018,.115]). These effect sizes converge on bilateral dorsal parietal–frontal and motion-sensitive visual regions, aligning with the hypothesis that expert neural geometry more strongly reflects abstract tactical structure and relational constraints.

RSA reveals strong expert-novice differences for dominant dimensions such as strategy and checkmate status, but this method is inherently limited to capturing features that drive the overall representational geometry ^46^. To assess whether subtler chess stimuli properties are also encoded, we applied multivariate decoding across eight dimensions (e.g., number of moves to checkmate, board difficulty; see Supplementary Sec. S3). RSA detected robust group differences in multiple ROIs for 3 dimensions, partial effects for 3 more, and none for the remaining 2. In contrast, decoding revealed expert advantages for 7 of 8 dimensions across widespread regions. These findings indicate that expertise supports a rich, sparser and more expressive representational space, where both dominant task-relevant properties and finer distinctions coexist.

### How Representations Are Structured: Compressed Geometry

We calculated the Participation Ratio (PR) ^47^–an effective dimensionality measure of the multivoxel covariance spectrum (higher PR = higher dimensionality)—to investigate whether expertise would result in lower-dimensional representations, which would manifest as a lower PR.

#### Participant classification based on dimensionality profiles

We next asked whether subjects’ full dimensionality profiles distinguish Experts from Novices. Dimensionality profiles for each participant across ROIs are shown as rows in Fig. 5A. When projected into two dimensions using PCA, each subject is represented as a point in Fig. 5B, with positioning determined by their full dimensionality profile. The classification analysis of these 2D projections revealed a clear separation of Experts and Novices (cross-validated accuracy = .800, permutation *p* = .0003, stratified 5-fold CV, *n* = 40 independent participants, 10,000 label permutations).

**Fig. 5.**
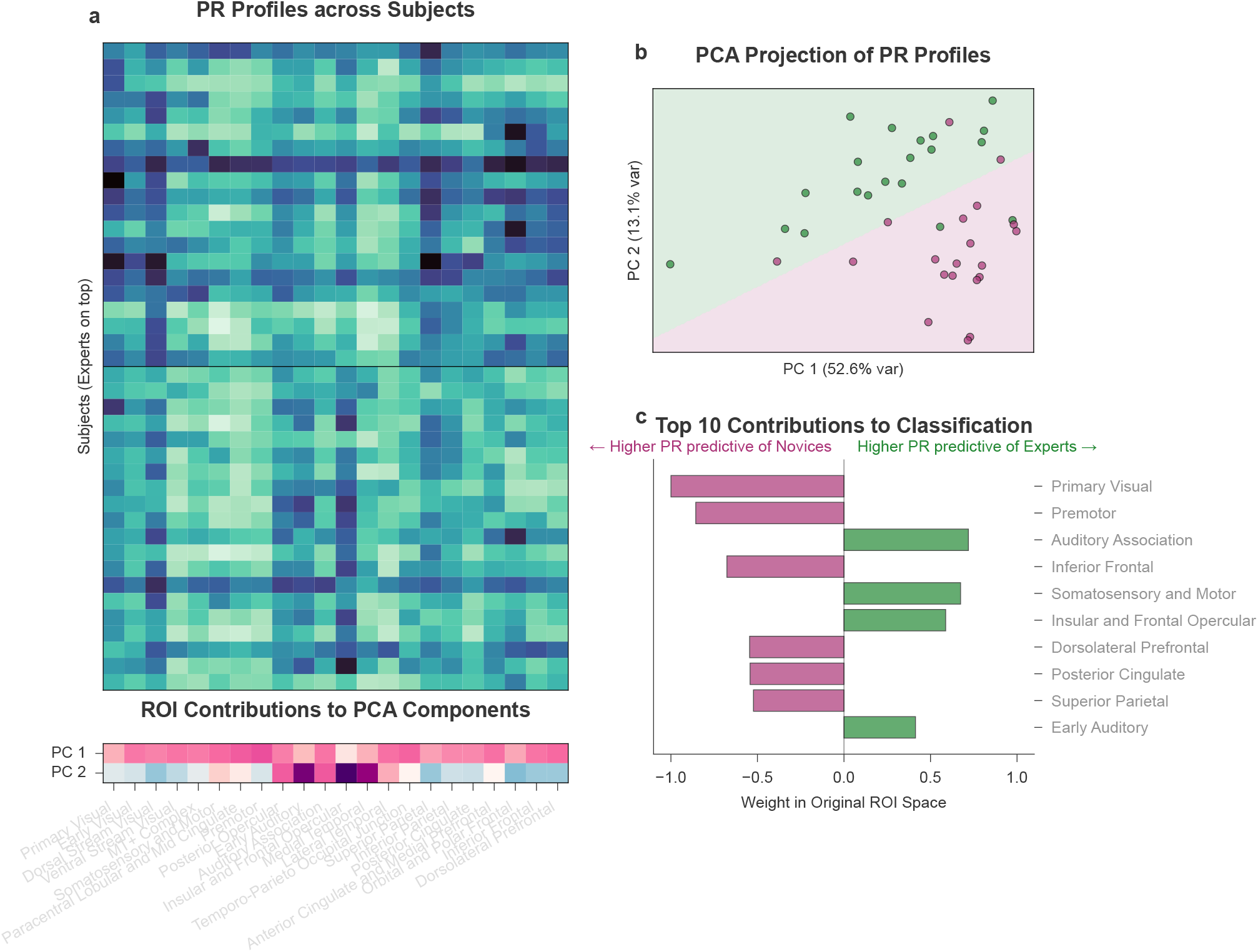
Participation Ratio (PR) analysis of representational dimensionality. **a**. Top: Heatmap of PR values across subjects and ROIs, sorted by group (Experts above and Novices below; *n* = 20 independent participants per group). Bottom: ROI loadings for the first two principal components derived from PCA on the 40 × 22 subject-by-ROI PR matrix (PC1: 52.6% variance explained; PC2: 13.1%). **b**. PCA projection of subjects based on their ROI-wise PR profiles. Each point represents one subject (green = Expert, *n* = 20; purple = Novice, *n* = 20). A logistic regression classifier was trained on the 2D projection; the overlaid decision boundary shows group-level differences in representational dimensionality. Cross-validated classification accuracy was assessed with stratified 5-fold cross-validation; significance was tested by a two-sided permutation test with 10,000 label permutations (cross-validated accuracy = .825, permutation *p* = .0002). **c**. Top 10 ROIs contributing most strongly to group classification, based on a logistic regression model trained on PR values in original ROI space (same cross-validation and permutation procedure as panel b). Positive weights (in green) indicate that higher PR values in a given ROI increase the likelihood of being classified as an Expert, whereas negative weights (in purple) indicate that higher PR values increase the likelihood of being classified as a Novice. Source data are provided as a Source Data file.

An exploratory assessment of the PCA loadings (Fig. 5A, bottom) indicated a strong influence of auditory and temporal regions on PC2, and conversely, a stronger influence of all other regions on PC1. This suggests that these sets of regions shape the geometry of the PCA space and drive the classifier’s decision. Consistent with this, the analysis of classifier weights in ROI space (Fig. 5C; cross-validated accuracy = .825, permutation *p* = .0002, stratified 5-fold CV, *n* = 40 independent participants, 10,000 label permutations) showed that regions with the lowest PR in Experts carried more negative classifier weights—indicating that higher dimensionality in these regions biased classification toward Novices—whereas regions with lower PR in Novices were associated with more positive weights. These classifier weights are purely descriptive and reflect the joint distribution of PR values across the full profile rather than isolated region-level effects. The interpretation of these findings is discussed further in Supplementary Sec. S4.

For statistics and more region-level inferences, Fig. 6 (top) shows the group-average PR values for Experts and Novices across cortical regions, obtained by averaging across participants column-wise in the matrix shown in Fig. 5A. The lower panel of Fig. 6 illustrates the group differences (Experts−Novices) along with their significance. A cross regions, most areas exhibit significantly lower PR values in Experts compared to Novices, suggesting that expert brains rely on more compressed representational structures. All reported *p*-values are FDR-corrected.

**Fig. 6.**
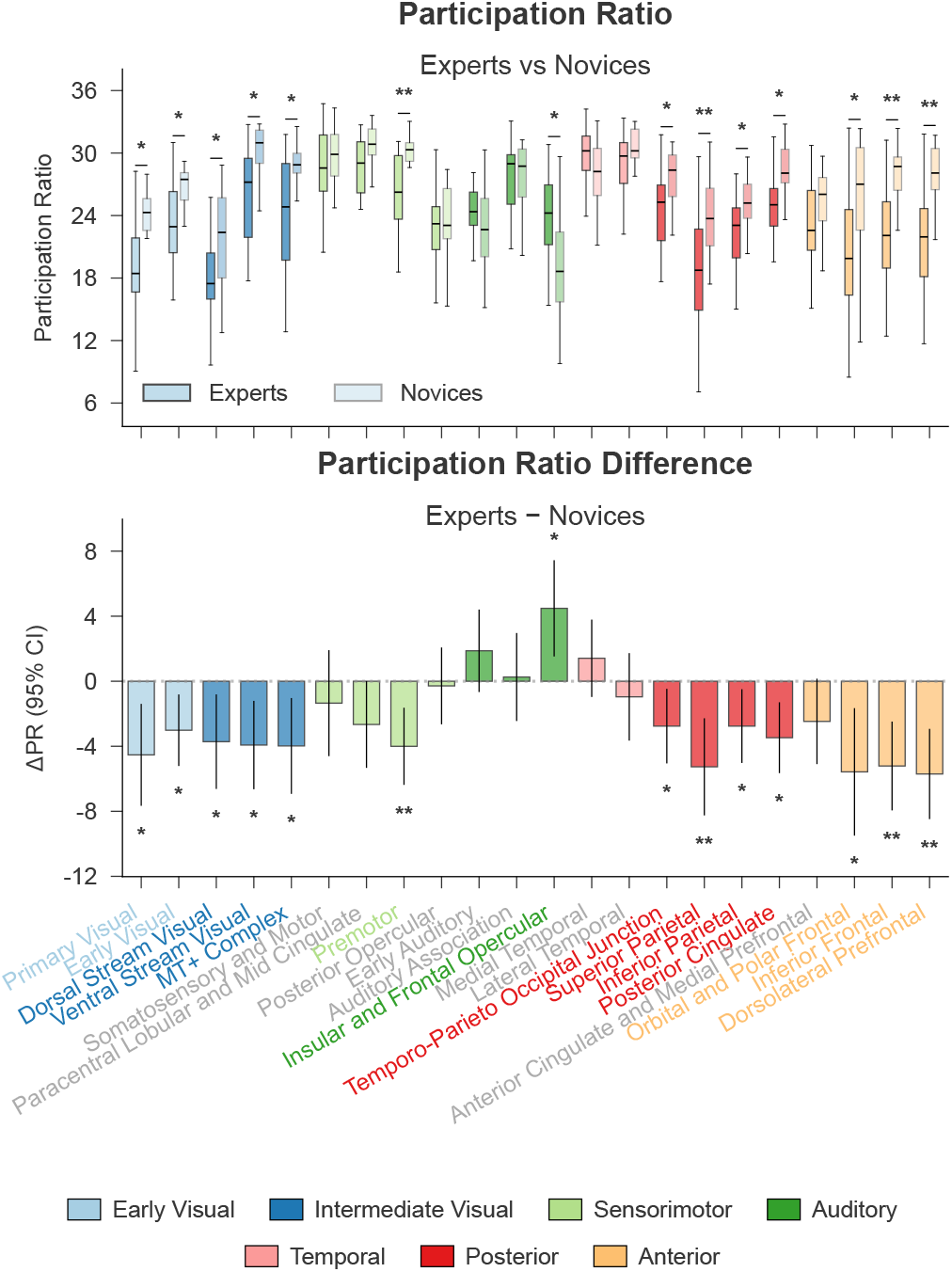
Participation Ratio (PR) analysis of representational dimensionality. Top: box plots show the distribution of per-subject PR values for Experts (*n* = 20 independent participants) and Novices (*n* = 20 independent participants) across 22 ROIs. Boxes span the interquartile range (IQR; 25th–75th percentile), centre lines mark the median, and whiskers extend to the furthest data point within 1.5 × IQR of the box edges. Lower values reflect reduced representational dimensionality in Experts. Bottom: ΔPR (Experts−Novices) per ROI; bar height is the mean group difference and error bars show the 95% confidence interval of that difference (Welch’s *t*-distribution with Welch–Satterthwaite degrees of freedom, computed via `scipy.stats.ttest_ind(equal_var=False`).`confidence_interval(0.95)`). Stars indicate two-sided Welch’s *t*-tests (`scipy.stats.ttest_ind` with `equal_var=False`), FDR-corrected (Benjamini–Hochberg) across the 22 ROIs at *α* = 0.05: ^∗^ *p*_FDR_ *<* .05, ^∗∗^ *p*_FDR_ *<* .01, ^∗∗∗^ *p*_FDR_ *<* .001. Per-ROI *t*-statistics, degrees of freedom, mean differences, 95% confidence intervals, Cohen’s *d*, raw and FDR-corrected *p*-values for all 22 ROIs are reported in Supplementary Table 8. Source data are provided as a Source Data file.

#### Dimensionality differences by ROI

Statistical comparisons revealed significant PR reductions in Experts in 14 of 22 regions after FDR correction for multiple comparisons (see Supplementary Table 8 for full statistics). These included early visual areas such as the Primary Visual (ΔPR = −4.53, *t*(30.1) = −2.95, *p*_FDR_ = .016, Cohen’s *d* = −0.93, 95% CI [−7.66, −1.39]) and Early Visual cortex (ΔPR = −3.01, *t*(29.4) = −2.80, *p*_FDR_ = .020, Cohen’s *d* = −0.89, 95% CI [−5.21, −0.81]), as well as higher-level visual processing areas such as the Ventral Stream Visual (ΔPR = −3.92, *t*(35.0) = −2.93, *p*_FDR_ = .016, Cohen’s *d* = −0.93, 95% CI [−6.64, −1.21]), MT+ Complex (ΔPR = −3.98, *t*(30.0) = −2.76, *p*_FDR_ = .020, Cohen’s *d* = −0.87, 95% CI [−6.93, −1.03]), and Dorsal Stream Visual cortex (ΔPR = −3.72, *t*(38.0) = −2.59, *p*_FDR_ = −.025, Cohen’s *d* = −0.82, 95% CI [−6.62, −0.81]). Significant effects extended to associative and executive areas, including the Premotor cortex (ΔPR = −4.00, *t*(31.7) = −3.44, *p*_FDR_ = .009, Cohen’s *d* = −1.09, 95% CI [−6.37, −1.63]), the Temporo-Parieto Occipital Junction (ΔPR = −2.76, *t*(36.9) = −2.44, *p*_FDR_ = .031, Cohen’s *d* = −0.77, 95% CI [−5.05, −0.47]), and parietal areas including Superior Parietal (ΔPR = −5.26, *t*(33.9) = −3.58, *p*_FDR_ = .008, Cohen’s *d* = −1.13, 95% CI [−8.25, −2.28]) and Inferior Parietal cortex (ΔPR = −2.76, *t*(30.2) = −2.49, *p*_FDR_ = −.031, Cohen’s *d* = −0.79, 95% CI [−5.02, −0.49]). Finally, effects were particularly robust in prefrontal areas: the Orbital and Polar Frontal cortex (ΔPR = −5.57, *t*(37.0) = −2.88, *p*_FDR_ = −.016, Cohen’s *d* = −0.91, 95% CI [−9.49, −1.65]), Inferior Frontal cortex (ΔPR = −5.21, *t*(36.7) = −3.87, *p*_FDR_ = −.005, Cohen’s *d* = −1.22, 95% CI [−7.94, −2.48]), and Dorsolateral Prefrontal cortex (ΔPR = −5.70, *t*(32.9) = −4.18, *p*_FDR_ = −.004, Cohen’s *d* = −1.32, 95% CI [−8.48, −2.93]). The only significant increase in dimensionality (Experts*>*Novices) was observed in the Insular and Frontal Opercular cortex (ΔPR = +4.49, *t*(37.3) = +3.07, *p*_FDR_ = −.015, Cohen’s *d* = +0.97, 95% CI [+1.53, +7.44]).

Strikingly, the reduction in PR is observed in each of the eight cortical regions that also showed an increase in representational content for the two tested high-level distinctions, Checkmate and Strategy (Fig. 4). Neural content (what) seems to be closely associated with neural geometry (how).

Together, the PR results indicate that expert representations are lower-dimensional across the very regions that carry relational/goal-relevant content. These dimensionality differences are not driven by differences in ROI size (Supplementary Sec. S5). It is important to note that this compression did not come out at the cost of information. The decoding analyses (Supplementary Sec. S3) revealed that experts achieved higher classification accuracy for fine-grained stimulus properties in many of the same regions that showed the largest dimensionality reductions. Expert representations are therefore paradoxically both more compact and more informative: a sparse, efficient code in which compression and discriminability coexist.

### Where Information Is Encoded: Domain-General Networks

To investigate which cognitive domains are preferentially engaged in chess experts, and how these domains relate to the representational changes that we observed, we examined the spatial correspondence between group-level brain activity and representational patterns and meta-analytic maps derived from Neurosynth. Specifically, we assessed the degree to which Experts and Novices recruited predefined functional networks by correlating group-level *z*-statistic maps with seven term-based meta-analytic association maps: *Working Memory, Memory Retrieval, Navigation, Language Network, Face Recognition, Object Recognition*, and *Early Visual*. These networks include both domain-general processes–most notably working memory and memory retrieval–and more domain-specific or perceptual systems that have been implicated in prior studies of visual and cognitive expertise (see Sec. 4.8.1 for a more detailed rationale).

We applied this correlation framework to both univariate second-level *t*-maps and multivariate RSA maps, offering complementary perspectives on how chess expertise modulates brain function. The univariate analysis captures differences in overall activation strength, while the RSA-based analysis reflects differences in local representational geometry (see Sec. 4.8) across three model-based stimulus dimensions: Checkmate, Strategy, and Visual Similarity. This procedure allowed us to characterise how differences in activation patterns and representational geometries align with canonical functional networks, thereby bridging the gap between computational and cognitive changes observed in expertise.

#### Univariate networks involvement

To assess group-level functional recruitment during board evaluation, we compared the positive (Experts *>* Novices) and negative (Novices *>* Experts) directional components of the group contrasts with meta-analytic term maps from Neurosynth (see Fig. 7 and Supplementary Table 10 for correlation summaries and 95% bootstrap confidence intervals). These analyses were conducted for two second-level contrasts: (i) All Boards *>* Rest, and (ii) Checkmate *>* Non-checkmate. All reported *p*-values are FDR-corrected at *α* = 0.05.

**Fig. 7.**
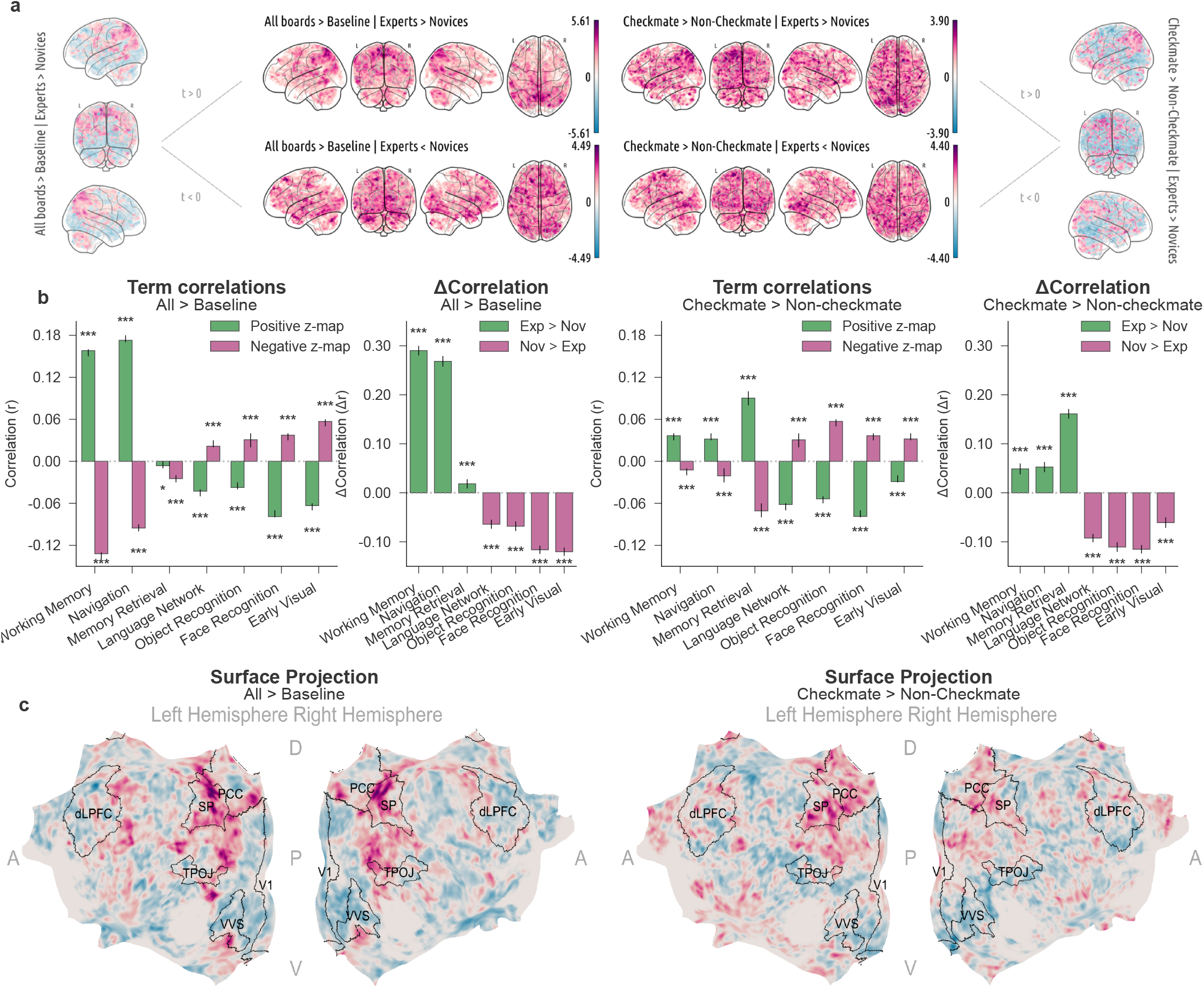
Neurosynth term correlations with univariate contrasts. **a**. Illustration of the preprocessing step applied to the group-level *t*-maps. For each contrast (All Boards *>* Baseline and Checkmate *>* Non-checkmate trials), the resulting second-level *t*-map was split into positive (Experts *>* Novices) and negative (Novices *>* Experts) components, converted to *z*-maps, and masked to retain only grey matter voxels. These maps were then used for the correlation analyses shown below. **b**. bar plots showing Pearson correlations (*r*) between each Neurosynth meta-analytic term map and the positive (Experts *>* Novices, green) and negative (Novices *>* Experts, purple) directional maps. Each bar is a single correlation across all grey-matter voxels (no per-subject distribution exists for this group-level summary; the underlying group *z*-maps were derived from *n* = 20 Experts and *n* = 20 Novices, independent participants). Error bars show 95% percentile bootstrap confidence intervals from 10,000 voxel-wise resamples. Significance was assessed with two-sided bootstrap *p*-values, FDR-corrected (Benjamini–Hochberg) across the 7 terms ×2 directions within each contrast: ^∗^ *p*_FDR_ *<* .05, ^∗∗^ *p*_FDR_ *<* .01, ^∗∗∗^ *p*_FDR_ *<* .001. Exact *p*-values are listed in Supplementary Table 10. **c**. surface projections of the bidirectional *z*-maps for each contrast. ROI contours are shown for reference (dlPFC: dorsolateral pre-frontal cortex; PCC: posterior cingulate; SP: superior parietal; TPOj: temporo-parieto-occipital junction; VVS: ventral visual stream; V1: primary visual). Source data are provided as a Source Data file.

For the All Boards *>* Rest contrast, Experts showed significantly stronger correlations with higher-order cognitive networks, including “Working Memory” (Δ*r* = .290, *p*_FDR_ *<* .001, 95% CI [.280,.300]), “Memory Retrieval” (Δ*r* = .018, *p*_FDR_ *<* .001, 95% CI [.009,.028]), and “Navigation” (Δ*r* = .268, *p*_FDR_ *<* .001, 95% CI [.258,.279]). In contrast, Novices exhibited stronger correlations with “Language Network” (Δ*r* = −.064, *p*_FDR_ *<* .001, 95% CI [−.073, −.055]), “Face Recognition” (Δ*r* = −.116, *p*_FDR_ *<* .001, 95% CI [−.125, −.108]), “Object Recognition” (Δ*r* = −.068, *p*_FDR_ *<* .001, 95% CI [−.078, −.058]), and “Early Visual” (Δ*r* = −.120, *p*_FDR_ *<* .001, 95% CI [−.129, −.112]).

For the more specific contrast of Checkmate *>* Non-checkmate boards, Experts again showed elevated similarity with maps associated with “Memory Retrieval” (Δ*r* = .161, *p*_FDR_ *<* .001, 95% CI [.151,.171]), “Navigation” (Δ*r* = .053, *p*_FDR_ *<* .001, 95% CI [.042,.063]), and “Working Memory” (Δ*r* = .049, *p*_FDR_ *<* .001, 95% CI [.038,.060]).

In this context, Novices displayed significantly stronger associations with “Language Network” (Δ*r* = −.092, *p*_FDR_ *<* .001, 95% CI [−.101, −.084]), “Object Recognition” (Δ*r* = −.111, *p*_FDR_ *<* .001, 95% CI [−.120, −.101]), “Face Recognition” (Δ*r* = −.115, *p*_FDR_ *<* .001, 95% CI [−.124, −.106]), and “Early Visual” (Δ*r* = −.061, *p*_FDR_ *<* .001, 95% CI [−.072, −.050]).

These findings suggest distinct neural strategies during board evaluation, with experts drawing more heavily on integrative, domain-general resources, and novices relying more on low-level visual and semantic processing. By comparing overlap between cognitive term maps and *t*-maps, we identified which functional networks are preferentially associated and geometrically aligned with each group’s activity, providing a computational and functional lens on spatial activation differences.

#### Multivariate representational shift

Our study was specifically designed to investigate the representational geometry underlying expertise. We extended the domain analysis to RSA searchlight maps, which capture local representational geometries based on three theoretical models: *Checkmate, Strategy*, and *Visual Similarity*. These group-level RSA maps were compared to the same set of meta-analytic Neurosynth term maps using the same procedure discussed in Sec. 4.8 (see Fig. 8, and Supplementary Table 9 for correlation summaries and 95% bootstrap confidence intervals). Unlike the correlation with univariate *t*-maps or coarser ROI-level RSA, this method reveals how small-scale activation patterns are organised relative to task-relevant properties, linking group-level differences in geometry to specific cognitive systems. All reported *p*-values are FDR-corrected at *α* = 0.05.

**Fig. 8.**
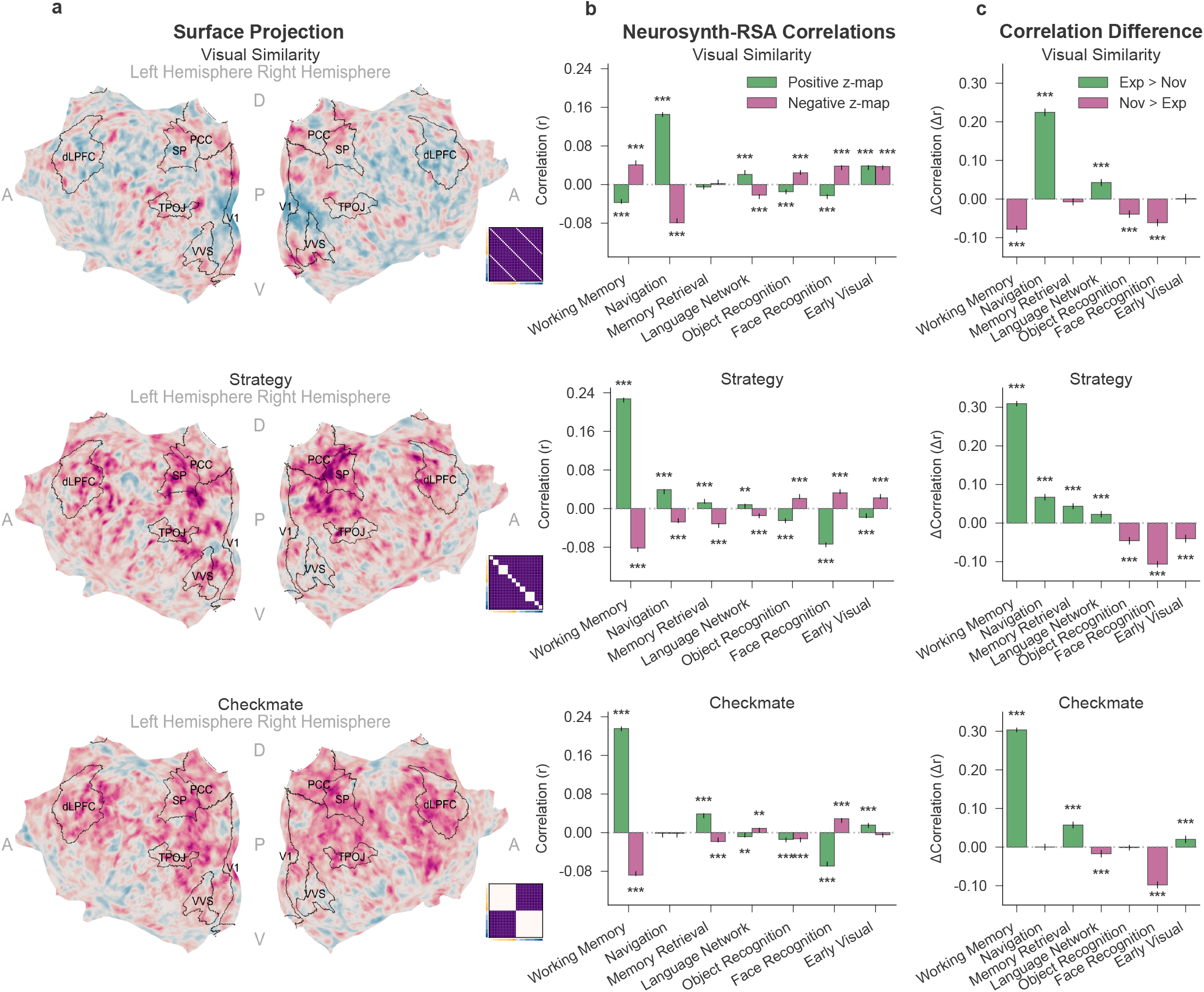
Group-level RSA correlations with Neurosynth term maps. For each of three model RDMs (Visual Similarity, Strategy, Checkmate; rows), the group-level RSA searchlight difference map (Experts − Novices) was split into positive (Experts *>* Novices) and negative (Novices *>* Experts) components and correlated with seven Neurosynth meta-analytic term maps selected *a priori*. Each RSA map reflects localised multivoxel dissimilarity patterns within grey matter. **a**. Flat left- and right-hemisphere surface projections of the RSA searchlight difference map for each model (warm: Experts *>* Novices; cool: Novices *>* Experts), with the corresponding model RDM as an inset; ROI contours are shown for reference (dlPFC: dorsolateral pre-frontal cortex; PCC: posterior cingulate; SP: superior parietal; TPOj: temporo-parieto-occipital junction; VVS: ventral visual stream; V1: primary visual). **b**. Bar plots of Pearson correlations between each term map and the positive (Experts *>* Novices, green) and negative (Novices *>* Experts, purple) directional components. **c**. Per-term difference between these two correlations (Δ*r* = *r*_positive_ −*r*_negative_); positive values (green) indicate a term more strongly associated with regions where Experts *>* Novices, negative values (purple) the reverse. In **b** and **c**, each bar is a single correlation across all grey-matter voxels (no per-subject distribution exists for this group-level summary; the underlying searchlight maps were derived from *n* = 20 Experts and *n* = 20 Novices, independent participants), and error bars show 95% percentile bootstrap confidence intervals from 10,000 voxel-wise resamples. Significance was assessed with two-sided bootstrap *p*-values, FDR-corrected (Benjamini–Hochberg) across the 7 terms separately within each series (the positive and negative components in **b** and their difference in **c**) and within each model: ^∗^ *p*_FDR_ *<* .05, ^∗∗^ *p*_FDR_ *<* .01, ^∗∗∗^ *p*_FDR_ *<* .001. Exact *p*-values are listed in Supplementary Table 9. Source data are provided as a Source Data file.

The *Checkmate* model revealed robust expert-related effects for “Working Memory” (Δ*r* = .304, *p*_FDR_ *<* .001, 95% CI [.298,.309]), indicating a stronger representational alignment between the working memory network and expert brain activity. A smaller expertise effect was observed for “Memory Retrieval” (Δ*r* = .057, *p*_FDR_ *<* .001, 95% CI [.049,.066]). Novices showed higher alignment for “Language Network” (Δ*r* = −.017, *p*_FDR_ *<* −.001, 95% CI [−.027, .008]) and “Face Recognition” (Δ*r* = −.098, *p*_FDR_ *<* −.001, 95% CI [−.107, .089]). A small expert-favouring effect was also found for “Early Visual” cortex (Δ*r* = .020, *p*_FDR_ *<* .001, 95% CI [.011,.030]), suggesting subtle representational organisations even in low-level perceptual areas. Overall, these patterns mirror the “goal achievement” nature of the Checkmate category.

The *Strategy* model showed even stronger expert-related domain effects. “Working Memory” again displayed a pronounced expert bias (Δ*r* = .309, *p*_FDR_ *<* .001, 95% CI [.302,.316]), followed by additional expertise effects for “Navigation” (Δ*r* = .067, *p*_FDR_ *<* .001, 95% CI [.058,.075]), “Memory Retrieval” (Δ*r* = .044, *p*_FDR_ *<* .001, 95% CI [−.036,.051]), and “Language Network” (Δ*r* = −.023, *p*_FDR_ *<* .001, 95% CI [−.014,.031]). Conversely, novices showed stronger alignment for “Object Recognition” (Δ*r* = −.046, *p*_FDR_ *<* .001, 95% CI [−.056, .036]), “Face Recognition” (Δ*r* = −.106, *p*_FDR_ *<* −.001, 95% CI [−.115, −.098]), and “Early Visual” (Δ*r* = −.040, *p*_FDR_ *<* .001, 95% CI [−.050, −.030]), possibly reflecting greater reliance on perceptual processes in novices. This supports the idea that experts encode multi-step relational motifs in memory-supported networks.

The *Visual Similarity* model presented a distinct profile. Experts showed stronger alignment with “Navigation” (Δ*r* = .225, *p*_FDR_ *<* .001, 95% CI [.215,.235]) and “Language Network” (Δ*r* = .043, *p*_FDR_ *<* .001, 95% CI [.034,.052]). Novices again exhibited stronger representational alignment in the “Object Recognition” network (Δ*r* = −.039, *p*_FDR_ *<* −.001, 95% CI [−.049, −.030]), “Face Recognition” (Δ*r* = .061, *p*_FDR_ *<* .001, 95% CI [−.071, −.052]), and notably for “Working Memory” (Δ*r* = .078, *p*_FDR_ *<* .001, 95% CI [−.087, −.069]). This divergence from the pattern in the other models highlights that when similarity is purely perceptual, novices rely more on perceptual systems. Experts’ representations, on the other hand, remain anchored to task-relevant spatial schemas (navigation) and verbal labelling.

Taken together, these results indicate that different RSA models capture partially distinct cognitive signatures. Because the representational approach has the power to differentiate between task-relevant dimensions (Checkmate and Strategy), it allows us to infer that task-relevant dimensions are mostly represented in regions involved in working memory and, to a lesser extent, memory retrieval. In contrast, the Visual Similarity model places greater weight on navigation in experts but diverges in its relationship to working memory, highlighting divergent information-processing priorities in the two groups. This provides a richer picture of how expertise shapes neural representations: not just where expertise effects emerge, but which cognitive systems are most closely tied to those computational changes, bridging local geometry with broader theories of domain-general and domain-specific reorganisation. Crucially, the three signatures do not arise in separate systems: the networks that encode relational content (**what**) and show compressed geometry (**how**) are the very systems experts preferentially recruit during evaluation (**where**).

### Skill-Gradient Analysis

To assess whether the observed representational differences scale continuously with chess skill rather than reflecting a purely categorical expert-novice distinction, we correlated neural metrics with two independent skill measures. Within the expert group, Elo rating correlated with mean Checkmate RSA model fit across ROIs (Pearson’s *r* = .47, *p* = .039, 95% CI [.03, .75], *n* = 20 Experts). Across all participants, familiarisation move accuracy on the very same board set used during fMRI correlated with multiple relational neural metrics (Supplementary Sec. S6). These results suggest that the representational reorganisation described above scales with chess-relevant competence.

## Discussion

How does extensive learning reshape neural representations? Rather than merely identifying *where* expertise resides in the brain, we asked *how* extensive training transforms the internal structure of neural codes and *what* (which content) is encoded in the expert brain. Using chess as a controlled, theory-rich domain, our findings reveal expertise-driven transformations in the content, structure, and neural substrates of representations, characterised by optimisation along high-level, goal-relevant dimensions. We interpret these transformations as a form of *network-code inversion*, providing a unified account that integrates classic cognitive theories, efficient coding, and modern manifold theory.

Experts prioritised relational content over surface features. Behaviourally, expert choices were more reliable (Supplementary Sec. S2) and better predicted by high-level, relational properties of the stimuli (Fig. 3). In-scanner board preference profiles reinforce this dissociation: within visually matched pairs, experts strongly preferred checkmate over non-checkmate boards, whereas novices preferred both at similar rates, with their choices governed by shared perceptual features (Supplementary Sec. S2). This divergence was mirrored in neural geometry: representational geometry in experts aligned more strongly with the same relational models in frontal and parietal cortices, whereas novices showed weaker or absent alignment (Fig. 4). The behavioural and neural results indicate a qualitative shift in ***what*** is encoded with expertise: away from appearance and toward high-level, relational properties. Rather than encoding perceptual properties, expert representational resources are allocated to behaviourally relevant dimensions (in line with efficient coding principles ^48^) to prioritise relational structure (in line with chunking and template theories ^17,43^). A perceptual-to-relational feature gradient analysis further confirmed this dissociation, with the vast majority of explained variance in expert preferences attributable to relational rather than perceptual features (Supplementary Fig. 3).

Classical chunking ^43^ and template theories ^17^ correctly predict tha experts focus on relational structure rather than surface features. But they do not specify the form or brain basis of this shift. Our results show that the shift is realised through low-dimensional, detail-preserving codes in domain-general frontoparietal systems. This adds neural and computational precision to existing theories.

Expertise alters not only what information is represented, but also ***how*** that information is structured in the brain. Using participation ratio as an effective dimensionality estimate, we found that expert neural codes were significantly more compressed across visual, parietal, and prefrontal regions compared to novices (Fig. 6; Sec. 2.2). Yet this compression did not entail information loss: regions with the largest drop supported the highest decoding of high-level, relational features (Supplementary Sec. S3). Fine-grained analyses showed reliable classification of subtle properties even within lower-dimensional manifolds (Supplementary Sec. S3). These findings suggest that compression is not incidental but instrumental in expertise: a representational strategy that improves separability, reduces noise, and enhances access to task-relevant features. By putting most of the useful signal into the few dimensions that matter, expertise yields representations that are paradoxically both more compact and more separable. A similar “dimensionality paradox” has recently been documented in short-term task learning, where practice simultaneously increases separability along task-relevant axes and compresses along irrelevant ones ^49^. Our findings suggest that this dual optimisation, initially observed over days of laboratory training, crystallises into a stable representational regime after years of real-world expertise.

This pattern matches manifold views of population coding and efficient-coding principles: learning rotates and compresses the representational space so that most variance lies along task-relevant dimensions, making behaviourally important states easier to separate while dampening nuisance variability ^48,50–52^. Just as the efficient coding principles predicted that V1 neurons would develop Gabor-like receptive fields optimised for natural image statistics ^53,54^, in cognitive domains like chess, a similar logic would lead to a representational basis that captures key structural properties of board configurations. The result is a flexible geometry that supports rapid readout, abstraction, and generalisation. It reflects a reorganisation from high-dimensional, entangled spaces to low-dimensional, task-tuned manifolds aligned with task demands.

Despite being more compact, expert representations retained fine-grained, task-specific detail. In focused analyses of checkmate positions (Supplementary Sec. S3), expert activity patterns reliably encoded subtle features (e.g., tactical motif, moves-to-mate) even though these properties did not dominate the global similarity structure (Supplementary Fig. 7). These variables were linearly decodable, indicating that detail is embedded in a sparse, multiplexed format: a few dominant dimensions capture abstract, global structure, while smaller, partly independent subspaces carry tactical motifs.

This organisation aligns with mixed-selectivity and factorised-coding accounts ^50,52^, and proves particularly helpful in resolving the breadth-depth trade-off in expertise: global dimensions support rapid evaluation, while local subspaces preserve detail for deeper inference. Put differently, it illustrates a pragmatic paradox of expertise–more is packed into less, yet the details that matter remain readily accessible.

Meta-analytic spatial correlations indicated that expertise reshapes both the engagement and representational role of large-scale brain systems (Sec. 4.8). Experts’ activation patterns showed stronger involvement of domain-general networks (working memory, navigation, retrieval–chiefly frontoparietal) whereas novices aligned more with sensory and language-related systems (face/object regions, early visual) (Fig. 7). Pattern-similarity analyses mirrored this divergence: in experts, domain-general systems carried high-level relational information, while in novices, task-relevant information remained anchored in domain-specific networks (Fig. 8). If stronger frontoparietal involvement merely reflected greater control demand, one would expect it to be more pronounced in novices, for whom the task was harder. Instead, novices aligned more strongly with perceptual and language-related systems, whereas the frontoparietal signature tracked expert relational coding. This frontoparietal locus converges with recent evidence that practice reshapes task representations toward conjunctive, task-tailored codes ^49,55^. The present findings extend this picture by showing that, in established expertise, the cortical systems carrying these restructured codes are domain-general frontoparietal networks that encode domain-specific relational content in a compressed, multiplexed format, a level of representational specificity that has not been characterised in practice-based paradigms.

This divergence likely reflects distinct cognitive and computational strategies. Experts appear to transition from a “lazy” learning regime, characterised by generic, sensory-bound coding, to a “rich” learning regime, where domain-general networks are actively reshaped into optimised, task-aligned representational systems ^56^. This richer representational regime need not replace the stronger engagement of domain-specific cortex observed in experts ^20–22^. Rather, improved perceptual templates in ventral/temporal regions might supply cleaner evidence to domain-general systems. Extracting the most informative latent (relational) variables probably depends on precise mid-level feature representations. This in turn would align with findings of greater activation in experts’ domain-specific regions ^6^.

In this sense, the change is less a switch of areas than a reallocation of representational content: domain-general networks encode domain-specific information in a structured, compressed, behaviourally optimised format, while domain-specific cortex continues to deliver high-quality inputs. Together with the *what* and *how* results, the ***where*** pattern shows that expertise involves a systematic reallocation and repurposing of neural resources. The end-result is a fundamental reorganisation in the anatomical substrates and computational architecture of expert cognition. It marks a shift from reactive, perceptual encoding to proactive, schematic reasoning. This reorganisation is based on tuned perceptual scaffolding and reconciles prior reports of higher activation in domain-specific cortex ^20,22^ with the present representational locus in domain-general systems. These results complement recent evidence suggesting that the rostrolateral prefrontal cortex supports relational categorisation during initial laboratory learning ^57^. Our results reveal the stable end-state of years of real-world expertise: a coordinated reorganisation of content, structure, and locus across the entire frontoparietal system, not just a single prefrontal focus during acquisition.

Several of these findings have no behavioural analogue and could not have been anticipated from existing theories of expertise. Effective dimensionality is a property of neural geometry, not of choice behaviour. The cortical locus of representational content (frontoparietal vs. sensory) is invisible to any behavioural measure. The multiplexed structure (global compression with local detail preservation in orthogonal subspaces) cannot be inferred from reaction times or accuracy. The co-occurrence and mutual constraint of these properties (the regions that carry relational content are the same ones that show compression and belong to domain-general networks) is what establishes the network-code inversion as a mechanistic account rather than a descriptive restatement of known behavioural effects. Novices, for their part, were actively engaged with the task and applied internally consistent evaluation strategies organised along perceptual rather than relational dimensions (Supplementary Sec. S2), confirming that the expert-novice contrast reflects a difference in representational content, not in task engagement. Moreover, estimated eye-movement patterns did not differ between groups (Supplementary Sec. S7), ruling out gaze strategy as a confound.

Our findings do not answer all questions, and further research will be needed. First, while we assessed robustness with cross-validated and convergent analyses (see SI), our inferences are based on crosssectional fMRI in a single, theory-rich model system, chess. It is known that cognitive mechanisms of expertise transfer well acros domains, including motor skills. The domain of chess has historically been a fertile ground for theories and hypotheses that have proven highly valuable in other domains. In the same spirit, the neural principles identified here (relational content, compact-yet-separable structure, targeted locus) remain to be tested for generality towards other domains beyond chess. Second, our ROI-based multivariate analyses focused on cortical representational geometry. An exploratory extension to nine subcortical ROIs (hippocampus, amygdala, caudate, putamen, pallidum, thalamus, nucleus accumbens, cerebellum, brain-stem) derived from the CAB-NP atlas ^58^ revealed no significant expert-novice differences in either RSA or decoding after FDR correction (Supplementary Sec. S8), suggesting that the representational reorganisation we report is predominantly cortical. Whether subcortical systems contribute to expertise through complementary mechanisms (e.g., procedural consolidation, reward-driven optimisation) remains an open question for future work. Third, we performed a cross-sectional study in which we compare experts with novices. The skill-gradient analysis (Sec. 2.4) provides initial evidence that the observed reorganisation scales with chess-relevant competence rather than reflecting a purely categorical distinction, but the restricted Elo range (1751–2269) limits power for detecting finer-grained within-expert gradients. It will be highly valuable to follow up this research with longitudinal designs and wider skill ranges that track expertise-related changes as expertise develops, in order to investigate the extent to which the changes in the what, how, and where of representational geometry follow the same temporal trajectory.

Keeping these limitations in mind, the current findings converge on a theoretical framework for understanding how expertise changes neural representations. Across behavioural, univariate, and multivariate analyses, expertise reshapes *what* is encoded (relational content over surface features), *how* it is structured (more compact yet more separable representations that preserve fine detail), and *where* it is expressed (domain-general frontoparietal involvement). We refer to this coordinated change as a ***network-code inversion***: novices rely on domain-specific sensory and semantic systems (network) to produce general-purpose, unstructured representations (code). They essentially apply generic perceptual and cognitive processes to the task of chess. Experts invert this pattern, recruiting domain-general systems associated with abstraction, working memory, and planning and repurposing them to represent domain-optimised content in a compact, structured, and task-optimised format.

Put simply, expertise packs more into less. This shift reflects a fundamental change in the internal representational geometry. Instead of increasing activation within existing circuits, the expert brain constructs a domain-aligned coordinate system tuned to the latent structure of the task. This system is low-dimensional and task-tuned: its dominant dimensions capture abstract structure for fast gist while smaller, partly independent subspaces preserve the tactical detail needed for appropriate responses.

This representational realignment grounds classic chunk/template accounts in measurable geometry. It bridges chunking theory through geometric compression, efficient coding through optimisation and dimensionality reduction, and flexible reasoning through low-dimensional, multiplexed manifolds. Variability is rotated and pooled onto meaningful dimensions, improving readout and reducing interference ^48,50–52^. This representational account yields clear, falsifiable predictions: with learning, alignment to relational models should strengthen, effective dimensionality should decrease while readout of task-relevant variables is maintained or improved, and expression should consolidate in frontoparietal systems.

We propose that this conceptual framework, anchored in chess as a model system, offers a general lens for understanding cognitive expertise as the emergence of structured internal representations optimised for the demands of the domain. The same metrics (e.g., geometry, separability, dimensionality, and locus) provide a portable assay for other domains (e.g., mathematics, language, complex motor skills) and enable evaluating computational models by how well their representations match expert structure. More broadly, the principle of packing more into less (i.e. compressing variability onto meaningful dimensions while keeping critical details readily retrievable) offers a common language for designing training, probing transfer, and aligning human and machine learning under the same representational criteria.

## Methods

All procedures complied with all relevant ethical regulations and were conducted in accordance with the latest version of the Declaration of Helsinki and the Belgian Law of 7 May 2004 on experiments on human persons. The study protocol was approved by the Ethics Committee Research UZ/KU Leuven (University Hospitals Leuven / KU Leuven; protocol number S62131; Belgian registration number B322201939337). All participants gave written informed consent prior to participation and received a fixed compensation of C25 per hour.

### Participants

Forty healthy adult volunteers took part and were assigned to two age- and education-matched groups based on chess expertise quantified by the Elo rating system ^59^. Elo is an interval-like metric updated after rated games (population mean ≈ 1500, SD ≈ 200) and tracks skill across the range (beginners ~ 800; novices ~ 1100; club players ~ 1500; experts ~ 2000; grandmasters ≥ 2500).

The expert group comprised twenty individuals (17 males, 3 females; mean age = 32.4 years, SD = 10.7, range = 18–52). All twenty had Elo ratings (mean = 2036, SD = 147, range = 1751–2269); nineteen were official tournament ratings and one (rating = 1824) was based on regular online play (e.g., Lichess, Chess.com).

The novice group comprised twenty participants (16 males, 4 females; mean age = 32.3 years, SD = 11.0, range = 22–55) who knew the rules of chess well (i.e., could identify legal moves and checkmates) but had no official Elo rating. Novices were competent hobby players and capable of completing the experimental task. They reported playing chess less than once per month, indicating that their skills, while sufficient for the study, were substantially lower than those of the experts.

Given the very large Expert−novice effects typically observed in chess-specific tasks (often Cohen’s *d* ≳ 1), a sample of 20 per group affords high power to detect group differences of this magnitude (two-tailed *α* = .05; e.g., ≈ .90–.95 for *d* ≈ 1.0).

All participants had completed at least high school education, gave written informed consent in accordance with ethical guidelines approved by the local research ethics committee, and completed a standardised MRI safety screening to ensure eligibility for scanning.

Sex was self-reported during the MRI safety screening. Sex was not a factor in study design — inclusion and group assignment were determined solely by chess expertise (Elo) and standard MRI eligibility criteria — and the sample is not statistically powered to test sex effects on the reported expert−novice contrasts. Sex distributions are reported above for each group; all subsequent analyses pool male and female participants within group, in keeping with the study’s primary aim of contrasting representational geometry by expertise.

### Stimuli

We constructed a set of 40 visually and categorically controlled chessboard stimuli designed to vary systematically along three dimensions of interest (Fig. 2A). The first dimension, *Checkmate vs. Non-checkmate*, reflects a high-level, relational property of the board, distinguishing positions that depict an impending checkmate (resolved in four moves or fewer) from those that do not. The second dimension, *Strategy*, captures differences in the tactical motifs and piece involved in the resolution of the game. Boards were grouped into ten strategy types (five for checkmate, five for their non-checkmate counterparts): queen-rook checkmates, queen-rook supported by minor pieces, knight-bishop combinations, bishop-forcing moves, and single-move easy checkmates, each paired with a corresponding non-checkmate strategy category. This dimension reflects both relational structure and visual identity of the involved pieces. The third dimension, *Visual similarity*, isolates low-level perceptual variation by matching each of the 20 checkmate boards with a visually similar non-checkmate counterpart, yielding 20 matched pairs. The difference between checkmate and non-checkmate boards is a minimal intervention, typically the repositioning of one or two pieces, that disables the checkmate. Stimuli were constructed to highlight these dimensions and enable representational similarity analysis of relational (checkmate positions), strategic (tactical motifs), and perceptual content (non-checkmate positions; see Fig. 2. Refer to S9 for a full list).

### Experimental Task and Design

#### Familiarisation Task

To promote recognition of the stimuli, assess participants’ ability to detect checkmate configurations, and reduce eye movements during scanning, we implemented a two-stage familiarisation procedure (Fig. 2D). First, participants were asked to complete an online task hosted on Pavlovia 48 to 24 hours prior to the scanning session, in which each of the 40 chessboard stimuli was presented in randomised order. Participants were instructed to enter the correct move in standard chess notation if they identified a checkmate sequence, or to skip the board otherwise. This task, performed without time constraints, allowed us to verify participants’ understanding of the stimuli while encouraging detailed inspection of each board.

On the day of the scan, participants were again shown all 40 stimuli during a 10-minute pre-scan review period. Boards were presented one by one and could be viewed freely, without any task or time pressure. This second exposure was designed to reinforce memory of the stimuli’s visual layout and strategic content, thereby minimising the need for visual search during scanning and reducing potential oculomotor artefacts. Together, these procedures ensured that participants entered the scanner with well-encoded representations of each board, allowing us to more precisely examine neural responses associated with the encoded chess-relevant properties.

#### fMRI Task

During scanning, participants performed a 1-back board evaluation task designed to engage domain-specific cognitive processes while remaining accessible to both expert and novice players. On each trial, a single chessboard (approximately 8° of visual angle) was presented for 2.5 seconds, followed by a 1-second inter-stimulus interval. On every trial, participants judged which of the two most recent boards presented a more strategically advantageous position for White, indicating their choice by pressing one of two buttons (two-alternative forced choice): one for the current board and one for the previous board. The button-to-response mapping was counterbalanced across runs within participants to control for motor confounds. We chose an active, domain-relevant task rather than passive viewing, since task context is known to strongly modulate object representations across the cortex ^60^. This way we were able to capture value-based, expertise-related differences while using a task that novices can in principle perform–albeit less effectively–, thereby providing rich and informative domain-specific encoding.

Each scanning session consisted of 5 to 10 functional runs, depending on individual tolerance and scanner-related constraints (the number of runs did not differ significantly between Experts (*M* = 8.4, *SD* = 2.0) and Novices (*M* = 9.5, *SD* = 1.4); Welch’s *t*(33.3) = −1.94, *p* = −.061, Cohen’s *d* = −0.61, 95% CI [−2.15, −0.05], Experts Novices). Note that novices completed slightly more runs on average, providing greater statistical power for this group; the key expert advantages reported throughout the paper are therefore conservative estimates (a run-matched control analysis confirmed that equalising run distributions across groups preserves all significant results; Supplementary Sec. S10). Each run began and ended with a 10-second fixation baseline and included 80 trials (i.e., each of the 40 boards presented twice in randomised order), for a total duration of 300 seconds. The fixation cross remained onscreen throughout to reduce eye movements, and participants were explicitly instructed to maintain fixation between trials and across runs. Stimuli were presented on a rear projection screen and viewed through an angled mirror mounted on the head coil. All experimental tasks were coded in MATLAB (The MathWorks, Natick, MA) using Psychtoolbox-3, and responses were recorded using an MRI-compatible button box, with both response accuracy and reaction times logged on every trial.

### MRI Data Acquisition and Preprocessing

MRI data were collected at the Department of Radiology, Universitair Ziekenhuis Leuven, using a 3T Philips Achieva dStream scanner equipped with a 32-channel head coil. Functional images were acquired using a multiband-accelerated T2*-weighted echo-planar imaging (EPI) sequence (multiband factor = 2) with the following parameters: TR = 2000 ms, TE = 30 ms, flip angle = 90^◦^, voxel size 2.0 x 2.0 x 2.7 mm^3^, and anterior-to-posterior phase encoding. Each volume comprised 30 axial slices acquired in ascending order, with multiband acceleration effectively covering 60 slices per TR. Participants were positioned supine with their head stabilized using foam padding, and scanner noise was attenuated using earplugs and over-the-ear headphones.

After five functional runs, a high-resolution T1-weighted structural scan was acquired using an MPRAGE sequence (1 mm isotropic voxel size) to serve as an anatomical reference. All MRI data were initially saved in DICOM format and subsequently converted to NIfTI using `dcm2niix` ^61^. The resulting data were organised according to the BIDS standard ^62^ to facilitate reproducibility and downstream processing.

Results included in this manuscript come from preprocessing performed using *fMRIPrep* 24.0.1 (^63^; RRID:SCR_016216), which is based on *Nipype* 1.8.6 (^64^; RRID:SCR_002502). All details about anatomical and functional pre-processing are reported in Supplementary Sec. S11.

### Regions of Interest (ROIs)

The ROIs were defined using the volumetric MNI projection of the Glasser parcellation ^65^, publicly distributed as `MNI_Glasser_HCP_2019_v1.0` via `afni_atlases_dist` (AFNI). Although the original Glasser atlas was defined on the cortical surface, we opted for the AFNI volumetric projection for several reasons. All analyses in the present study (RSA, decoding, participation ratio, Neurosynth correlations) operate on volumetric MNI-space data; using a surface-based parcellation would require projecting BOLD signals onto the cortical surface, introducing interpolation artefacts that are avoided entirely by projecting the atlas into volume space instead. Furthermore, any localisation error introduced by the volume projection at the level of the 360 fine-grained parcels is largely mitigated by our grouping scheme: hemispheric partitions were merged into bilateral masks (180 ROIs), and grouped by the coarse labelling in the original parcellation (22 ROIs) ^65^ (see Fig. 4). At this spatial scale, the grouped ROIs are sufficiently large that residual projection error has negligible impact on the analyses. An exploratory extension to subcortical structures using the CAB-NP atlas is reported in Supplementary Sec. S8.

### fMRI Univariate Analysis

Preprocessed fMRI data (in MNI space, 2 mm resolution) were analysed using a first-level General Linear Model (GLM) implemented in SPM12. For each participant, the GLM was estimated twice: onc using spatially smoothed functional images (with a Gaussian kernel with full-width at half-maximum in all directions: [4, 4, 4] mm), and once using un-smoothed images. The smoothed dataset was used for second-level univariate group analyses and neurosynth map correlations (see Sec. 4.8), while the un-smoothed dataset was reserved for multivariate and dimensionality analyses, including representational similarity analysis (RSA), decoding and participation ratio (PR) estimation.

The design matrix for each run included 40 regressors of interest– one for each unique chessboard stimulus. In addition, eight nuisance regressors (previously estimated with fMRIprep) were included to account for global signal fluctuations, six rigid-body motion parameters, and frame-wise displacement. Each trial was modeled according to the time the corresponding chessboard was on screen. Inter-stimulus intervals and fixation blocks were not explicitly modeled, and thus served as an implicit baseline. This approach produced one beta image per board (i.e., 40) per run for each participant.

For analyses involving un-smoothed data, no contrasts were applied at the first level, as the beta estimates themselves were used to extract multivoxel response patterns. For the smoothed data, two first-level contrasts were specified: *Checkmate > Non-checkmate*, identifying regions more active during checkmate boards relative to non checkmate boards, and *All boards > Rest*, identifying task-engaged voxels relative to baseline. These first-level contrasts were entered into second-level group analyses in SPM. For each contrast, we performed a between-group comparison (*Experts > Novices*) to identify regions where neural activations differed between the two groups, which provided the second-level *t*-maps used in the univariate Neurosynth maps correlation (Sec. 4.8) and ROIs analysis (Supplementary Sec. S12).

### Representational Similarity Analysis (RSA)

We used representational similarity analysis (RSA) ^33^ to characterise the geometry of stimulus representations in both behaviour and brain. This approach allowed us to assess how neural patterns in different brain regions reflect the structure of the experimental dimensions, and whether this alignment differs across groups. Specifically, we constructed representational dissimilarity matrices (RDMs) at the level of individual participants and regions of interest (ROIs), and compared them to theoretical model RDMs derived from stimulus properties.

#### Theoretical RDMs

To formalise the predicted representational structure associated with each experimental condition, we constructed binary theoretical RDMs for *Checkmate, Strategy*, and *Visual similarity*. Each model RDM encoded pairwise dissimilarities between stimuli based on their categorical labels. For instance, in the Checkmate model, all boards labelled as checkmate were considered similar to each other and dissimilar from all non-checkmate boards, resulting in a matrix with zeros along within-category pairs and ones for across-category comparisons. The same logic was applied to the Strategy and Visual similarity models, where board identities were defined according to shared tactical motifs or matched visual configurations, respectively. These RDMs thus provide idealised predictions of how stimuli should be organised in representational space if neural or behavioural responses are structured primarily along a given dimension (see Fig. 2C). Interpreting alignment with a model RDM therefore reveals which stimulus feature–relational, strategic, or perceptual–dominates the underlying representational geometry.

#### Behavioural Representational Similarity Analysis

From the raw trial-level behavioural data obtained during scanning, we first derived pairwise preference comparisons between stimuli. Each time a participant judged one board more favorable than another, we recorded the preferred and non-preferred stimuli. Using these pairwise ecomparisons pooled across all participants within each group, we computed two group-level matrices (one for experts, one for novices): a directional preference matrix and a symmetric behavioural RDM (See Fig. 3A).

The directional preference matrix captured the signed difference in how often one board was chosen over another. Specifically, cell (*i, j*) reflected the net number of times stimulus *i* was preferred over stimulus *j*. A positive value indicated a preference for *i* over *j*, while a negative value indicated the reverse. Because the 1-back task compares only consecutive boards, different stimulus pairs are compared different numbers of times depending on the randomised presentation sequence. To prevent pairs compared more often from accumulating higher raw counts independently of preference consistency, we normalised each entry by the total number of comparisons for that pair:

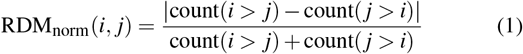

Values range from 0 (perfectly tied preferences) to 1 (perfectly consistent preference direction); pairs with zero comparisons were set to 0. The directional preference matrix is reported in raw counts in Fig. 3A to convey the group-level preference structure, while the count-normalised behavioural RDM is what we use for all subsequent behavioural RSA analyses.

To further explore stimulus-level preference profiles, we computed the selection frequency for each stimulus in all comparisons (Fig. 3C). Specifically, we counted how often each board was chosen as the preferred in a pair. These frequencies were visualised as bar plots, with the bars coloured by checkmate status and shaded by strategy identity. This visualisation allowed us to detect stimuli that were consistently favoured or avoided (i.e., they had a higher or lower perceived value), regardless of the specific board they were paired with.

Finally, we assessed the degree to which behavioural preferences aligned with theoretically motivated models of board similarity (Fig. 3D). For each theoretical RDM (checkmate, strategy, visual similarity), we computed Pearson correlations between the lower triangles of the group-level symmetric behavioural RDM and each model RDM. To quantify the reliability of each correlation, we used a nonparametric bootstrap procedure (10,000 resamples over RDM pairwise measures) to compute 95% confidence intervals around the correlation coefficient. This procedure was repeated separately for experts and novices, enabling direct comparisons of the extent to which each group’s preferences reflected structural properties of the stimuli. All *p*-values were corrected for multiple comparisons across the three model tests within each group using the Benjamini-Hochberg false discovery rate (FDR) procedure at *α* = 0.05.

#### fMRI Representational Similarity Analysis

Representational similarity analysis (RSA) was performed in MATLAB R2022b using the CoSMoMVPA toolbox ^66^. As detailed in Sec. 4.6, we used unsmoothed beta estimates for each of the 40 experimental conditions. For each subject, multivoxel activity patterns for each unique stimulus were averaged across runs to increase reliability, yielding one beta pattern per stimulus per subject. Neural RDMs were then computed from these run-averaged patterns extracted from predefined ROIs based on the volumetric projection of the Glasser parcellation ^65^ (see Sec. 4.5) or spherical neighborhoods. Neural RDMs were built by computing pairwise correlation distances (1 − *r*), which capture angular pattern similarity independently of response magnitude ^67^, between multivoxel patterns for all conditions, separately for each subject. Model RDMs for categorical variables (checkmate status, strategy, visual similarity, motif) were constructed as binary dissimilarity matrices (0 if same category, 1 if different), while model RDMs for continuous features were based on pairwise feature differences. Following CoSMoMVPA recommendations, we applied voxel-wise mean centering prior to computing correlation distances. Model RDMs reflected high-level conceptual dimensions (e.g., Checkmate, Strategy) and lower-level perceptual features (e.g., Visual similarity), as described in Sec. 4.7.1.

##### RSA on Regions of Interest

ROI-based RSA was conducted using the `©cosmo_target_dsm_corr_measure` function in CoSMoMVPA. For each subject, ROI, and model regressor, the neural RDM was correlated with the corresponding model RDM using Pearson correlation, yielding a single similarity value per combination. This quantified the degree to which the neural representational geometry aligned with the hypothesized category structure.

To assess statistical significance, subject-level RSA values were tested against zero using one-tailed one-sample *t*-tests within each group (Experts and Novices). Group differences were evaluated using two-tailed Welch’s *t*-tests. All *p*-values were corrected for multiple comparisons across ROIs using a False Discovery Rate (FDR) procedure with *α <* .05.

##### Whole-brain Searchlight RSA

To extend the analysis beyond predefined ROIs, we conducted whole-brain searchlight RSA using spherical neighborhoods (radius = 3 voxels, 6 mm) in CoSMoMVPA. At each voxel, a local RDM was computed from the surrounding multivoxel activity pattern using correlation distance and compared (Pearson) to each model RDM. This yielded one unthresholded RSA map per subject and model, indicating how well local representational geometry matched the predicted structure. No voxelwise multiple-comparison correction was applied to the searchlight maps, as they were not used for voxelwise inference; instead, they served as inputs to the spatial correlation analysis with Neurosynth term maps (Sec. 4.8), where statistical testing was performed at the level of map-wise correlations with FDR correction.

### Meta-Analytic Correlation Analysis

To place the observed expertise effects in a broader functional context, we quantified the spatial similarity between our group-level statistical maps and large-scale functional networks derived from automated meta-analysis. Specifically, we correlated (i) the second-level univariate *spmT* maps (Experts *>* Novices; see section 4.6) and (ii) group-level representational similarity analysis search-light maps (section 4.7.3) with term-based association maps from the Neurosynth database ^68^.

#### Neurosynth term maps

Seven terms, selected *a priori* for their relevance to chess and cognitive expertise more broadly, were included in the meta-analytic analysis. Working memory was included as a proxy for the frontoparietal multiple-demand network, which has been implicated in expertise across different domains, and reflects the role of efficient information storage and manipulation in expert performance ^19,69^. Memory retrieval was selected to capture expert-specific mechanisms for efficient information access ^69,70^. Navigation was included due to the involvement of spatial systems in processing complex board configurations, as established in chess-specific research ^26,71^. The language network term was selected to capture verbal and propositional reasoning processes, which are thought to be more prominent in novice chess players ^24,72^. Face recognition was included given prior evidence from both chess-specific studies and broader expertise research implicating fusiform cortex and FFA in expertise-related visual processing ^45,73,74^. Early visual processing was assessed to examine reliance on low-level visual features, while object recognition was included as it is frequently associated with shape-based processing and perceptual expertise in domains requiring fine-grained discrimination ^25,75^ (see Supplementary Fig. 16).

For each term, we downloaded the z-scored association test map from the Neurosynth database and resampled it to the resolution of our group-level statistical image using linear interpolation. All subsequent voxelwise analyses were restricted to grey matter, defined using the ICBM152 2009c probabilistic grey matter template thresholded at *>* 0.5 and resampled to the target image space. This mask included both cortical and subcortical grey matter regions.

#### Univariate maps preparation

From the second-level GLM analysis (*All boards > Baseline* or *Checkmate > Non-checkmate* first-level contrasts; *Experts > Novices* second-level contrast, see Sec. 4.6), we extracted *t*-maps representing the difference in activation between Experts and Novices in the first-level contrasts. All *t*-maps were converted to *z*-maps before the group analysis following CoSMoMVPA recommendations, although preliminary tests showed almost identical results when using *t*-maps directly. The group-level *t*-maps (df = 38) were imported in Python, and converted to a signed, two-tailed *z*-map by (i) computing two-tailed voxel-wise *p*-values from the *t*-distribution and (ii) applying the inverse normal survival function while preserving sign. The resulting image was split into two non-overlapping one-tailed maps: a *positive* map (*Experts > Novices, z >* 0) and a *negative* map (*Novices > Experts, z <* 0, then multiplied by −1).

#### RSA maps preparation

To extend the meta-analytic correlation analysis beyond univariate activation patterns, we applied the same procedure to the group-level maps derived from the whole-brain RSA analysis (see Sec. 4.7.3.2).

For each RSA model (Checkmate, Strategy, Visual similarity) subject-level search-light correlation maps were Fisher-transformed (*z* = arctanh *r*). Fisher *z*-maps (20 experts, 20 novices) were entered into a second-level general linear model using `nilearn.glm.SecondLevelModel` with an intercept and a group regressor (Experts = +1, Novices = −1). The resulting *z*-map for the contrast *Experts > Novices* was split into positive and sign-flipped negative components as described above.

#### Correlation and statistical inference

Before computing correlations, voxels were cleaned using three criteria applied jointly across the positive map, negative map, and term map: (i) exclusion of voxels with non-finite values (NaN or Inf) in any map; (ii) exclusion of voxels with variance *<* 10^−5^ across the three maps (removing constant and near-constant voxels); and (iii) restriction to grey matter voxels.

For every term, Pearson’s correlation coefficient (*r*) was computed between the term map and (i) the positive and (ii) the negative directional map on the cleaned voxel set. Statistical inference was based on a non-parametric bootstrap executed on voxels (10000 iterations, percentile confidence intervals). In each iteration voxel indices were sampled with replacement, the correlation recomputed, and the distribution of bootstrap *r*-values used to derive two-sided 95% confidence intervals (CI) and empirical *p*-values. Benjamini–Hochberg false-discovery-rate correction (*α* = 0.05) was applied separately to (a) the set of positive correlations and (b) the set of negative correlations.

To determine whether a term was differentially associated with experts versus novices, the difference Δ*r* = *r*_positive_ − *r*_negative_ was bootstrapped (10000 iterations) to obtain the percentile-based 95% confidence interval and a two-tailed *p*-value using the same voxel resampling scheme. Two-sided *p*-values were computed as 2 min {*P*(Δ*r* ≤ 0), *P*(Δ*r* ≥ 0)} and FDR-corrected across the seven terms. Results are reported as Δ*r* with 95% CI and FDR-corrected *p*-values.

### Participation Ratio

We assessed the effective dimensionality of multivoxel representations within each ROI using the Participation Ratio (PR) ^47^, a metric that captures how variance is distributed across principal components. Higher PR values indicate that variance is more evenly distributed across components, reflecting a higher-dimensional, less compressed representational space. This approach allows us to probe the extent to whic neural representations differ in compactness across groups, with the hypothesis that expertise might be associated with more compressed or structured representational geometries.

We computed the PR separately for each subject and each ROI. To do so, we extracted voxel-wise beta estimates from un-smoothed first-level GLM results obtained in SPM12. For each subject, we first identified all beta images corresponding to individual chess boards across all runs, and averaged them within each condition to produce one beta image per unique board.

Within each ROI, we constructed a matrix of shape (*n*_boards_ × *n*_voxels_), where each row corresponds to the voxel activity pattern evoked by a specific chess board. PCA was performed on the resulting matrix, retaining all available components (min(*n* −1, *v*)), where *n* is the number of beta images (i.e., unique board conditions) and *v* is the number of voxels in the ROI. The PR was computed from the explained variance of these components, using the formula:

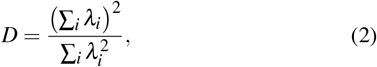

where *λ*_*i*_ denotes the eigenvalues of the PCA-decomposed covariance matrix. This value quantifies the effective number of components contributing substantially to the variance in the data. This procedure yielded one PR value per ROI, per subject.

Group-level comparisons of PR values between Experts and Novices were performed using Welch’s two-sample *t*-tests (unequal variance assumption) for each ROI. Significance was determined via two-tailed *t*-tests, and *p*-values were corrected for multiple comparisons using a False Discovery Rate (FDR) procedure with *α* = .05.

To explore whether subject-wise PR profiles contained group-level structure, we applied PCA to the standardised PR matrix (subjects × ROIs). This dimensionality reduction enabled us to visualise the dominant axes of variance across individuals in a 2D space. The resulting projection (Fig. 5B) is shown for visualisation purposes, and was used to assess whether Experts and Novices form separable clusters in PR space.

A logistic regression classifier trained on the 2D coordinates was overlaid to provide a reference linear decision boundary between groups. Classification accuracy was evaluated using stratified 5-fold cross-validation. Statistical significance was assessed via permutation testing with 10,000 iterations, in which group labels were randomly shuffled and cross-validated accuracy recomputed to build a null distribution. P-values were computed as the proportion of permuted accuracies exceeding the observed accuracy. This procedure was applied to both the full 22-dimensional PR space and the 2D PCA projection (using a pipeline comprising standardisation, PCA to 2 dimensions, and logistic regression, with all steps fit within each training fold).

To complement the ROI-wise group comparisons of PR values, we trained a second logistic regression classifier on the full set of ROI-wise PR values (before 2-D projection, without cross-validation) to explore which regions were most informative for distinguishing Experts from Novices. The resulting weight vector was interpreted descriptively as an exploratory ranking of ROI contributions to group classification. In this ranking, a PR increase in ROIs with positive weights would shift the classifier prediction toward the Expert group, while higher values in ROIs with negative weights would predict Novices’ PR profiles. We visualised the top 10 contributing ROIs in Fig. 5C to qualitatively assess which regions most strongly aligned with the separation between groups. This ranking is not used for statistical inference but serves to support and contextualise the group-level *t*-tests, highlighting regions more predictive of compressed representational geometry.

### Software

Behavioural and stimulus presentation: MATLAB R2022b (The Math-Works, Natick, MA) with Psychtoolbox-3 (^76^; RRID:SCR_002881). MRI preprocessing: *fMRIPrep* 24.0.1 (^63^; RRID:SCR_016216), built on *Nipype* 1.8.6 (^64^; RRID:SCR_002502). First-level GLM and contrasts: SPM12 (Wellcome Centre for Human Neuroimaging; RRID:SCR_007037). Multivariate pattern analyses (RSA, decoding): CoSMoMVPA (^66^; RRID:SCR_014519) under MATLAB R2022b. Group-level statistical analyses, manifold dimensionality, Neurosynth correlations, and figure generation: Python 3.11.6 with NumPy 1.26.4, SciPy 1.15.1, pandas 2.3.1, statsmodels 0.14.4, scikit-learn 1.7.1, Ni-Babel 5.3.2, Nilearn 0.11.1, Matplotlib 3.10.5, and seaborn 0.13.2. Eye-tracking decoding: scikit-learn 1.7.1 (RRID:SCR_002577). All package versions are pinned in the project’s `environment.yml` (Code Availability).

## Data Availability

Raw neuroimaging data (BIDS format), behavioural logs, stimuli, preprocessed MRI, and all subject-level analysis derivatives are deposited in the KU Leuven Research Data Repository (RDR) at 10.48804/VVCEWP ^77^. Access is granted through formal request via the repository, in compliance with ethical approval and GDPR requirements. Source data are provided with this paper.

## Code Availability

All analysis code, figure-generation scripts, and statistical pipelines are publicly available at costantinoai/chess-expertise-2025 and archived on Zenodo at 10.5281/zenodo.19392282 (concept DOI) ^78^. The code repository contains every pipeline needed to reproduce the paper, together with the group-level statistics, tables, and publication figures. Raw MRI, preprocessed data, and subject-level analysis derivatives are available from the RDR dataset cited above.

## Acknowledgements

We gratefully acknowledge Dr. Ronald Peeters (Radiology Department, UZ Leuven University Hospital) for his support in the preparation of the scanning protocol. We also thank Esna Mualla Gunay and Laura Van Hove (KU Leuven) for their help with recruitment and data collection. We are grateful to Ilkka Muukkonen, Shany Grossman and Luca Turella for their comment on an earlier version of the manuscript.

## Funding

H.O.B. was supported by the Research Foundation – Flanders (FWO; grant G0D3322N) and the Flemish Government (Methusalem grant METH/24/003).

## Author Contributions

A.I.C., H.O.B., M.B., and A.P. conceived the research idea and designed the experimental framework. A.I.C. and H.O.B. determined the hypotheses and supervised the project. A.I.C. and H.O.B. developed the methodology and experimental procedures. A.I.C. and E.V.H. performed the experiments and collected the data. F.F.V, H.O.B. and A.P. created the stimuli and datasets. Pilot testing was conducted by A.I.C., A.P., F.F.V. and E.V.H. A.I.C. processed and analysed the data, developed analysis scripts, and generated figures. A.I.C., H.O.B. and M.B. interpreted the results. A.I.C. wrote the initial draft of the manuscript and the Supplementary Information. A.I.C., H.O.B., M.B. and E.V.H. revised the manuscript. H.O.B., M.B. and A.I.C. supervised the work.

A.I.C. coordinated activities between institutions. H.O.B. acquired funding and provided laboratory resources. All authors discussed the results and commented on the manuscript at all stages.

## Declaration of Competing Interests

The authors declare no competing interests.

## Supplementary Information

Low-Dimensional and Optimised Representations of High-Level Information in the Expert Brain

Andrea I. Costantino, Artem Platonov, Felipe Fontana Vieira, Emily Van Hove, Merim Bilalić, Hans Op de Beeck

## Supplementary Material

**Supplementary Figure 1.**
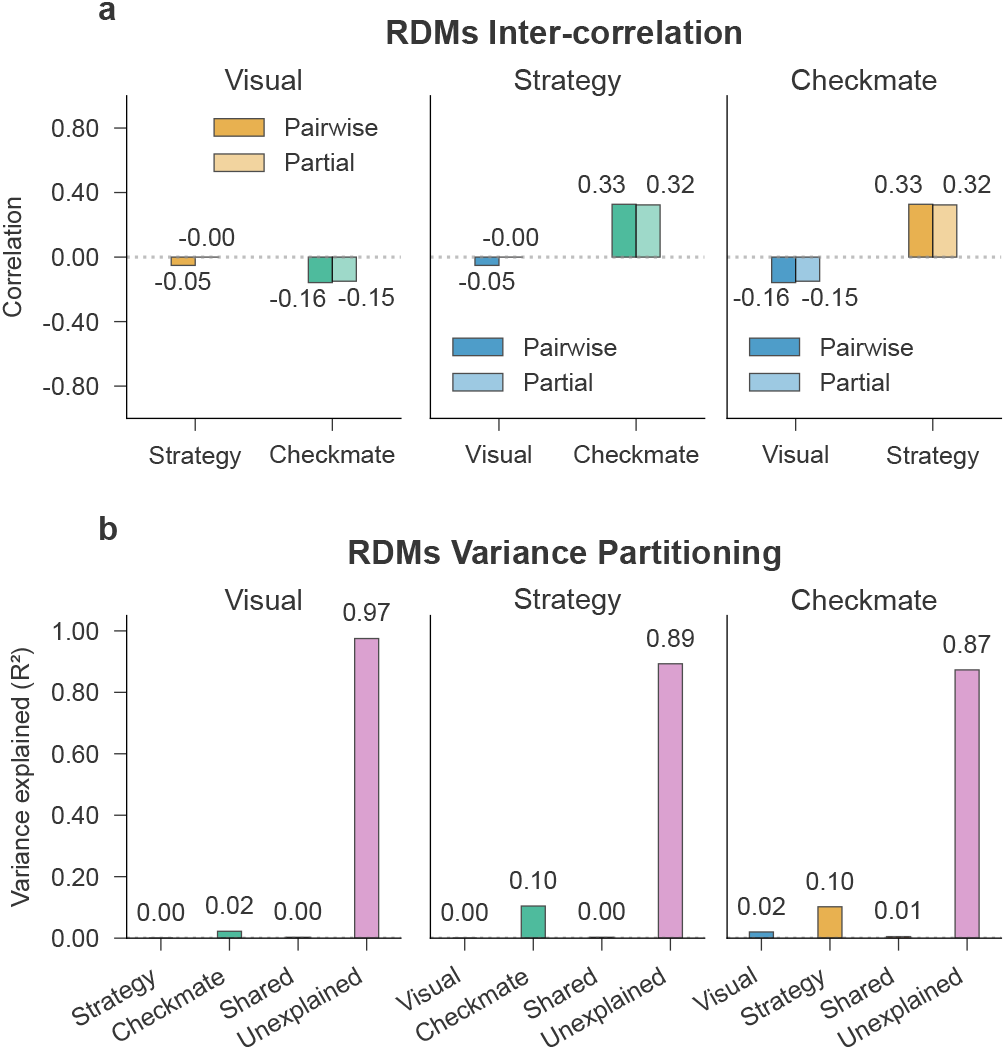
Orthogonality of representational dissimilarity matrices. **a**. Bar plots showing the Spearman correlation between each target RDM and the other two predictors, both as pairwise correlations and partial correlations (controlling for the third RDM). Each bar reflects the strength of association (*ρ*), with partial correlations labeled as “(partial X)” to indicate the variable that was controlled for. **b**. Variance partitioning results for each target RDM. Bars indicate the proportion of variance uniquely explained by each predictor RDM, variance shared between predictors, and residual (unexplained) variance. Larger unique and smaller shared contributions suggest greater orthogonality among representational structures. All values are computed on the lower triangle of the 40 × 40 stimulus RDMs (*n* = 780 stimulus pairs); analyses are descriptive (Spearman correlations and ordinary-least-squares variance partitioning) with no inferential test applied. Source data are provided as a Source Data file.

### S1 Orthogonality Across RDMs

To quantify the degree of overlap and separability among representational structures in our dataset, we first computed pairwise Spearman correlations between the flattened upper triangle of each model RDM (excluding the diagonal), providing a measure of shared variance in representational structure. We then computed partial Spearman correlations, which estimate the relationship between two RDMs while controlling for variance explained by the third. For instance, the partial correlation between the Checkmate and Strategy RDMs, controlling for Visual Similarity, reflects their unique association independent of the shared structure with the Visual RDM.

To further disentangle representational overlap, we further conducted a variance partitioning analysis based on nested linear regression models. Each RDM was treated in turn as a dependent variable, with the remaining two serving as predictors. We fit a full multiple linear model using both predictors to explain variance in the target RDM, and two reduced models each omitting one predictor. The unique variance explained by each predictor was calculated as the difference in *R*^2^ between the full model and the corresponding reduced model.

Shared variance was defined as the portion of *R*^2^ explained by both predictors jointly, beyond their individual contributions, and residual variance captured the unexplained portion. Formally, given a target RDM *Y* and two predictors *X*_1_ and *X*_2_, we define:

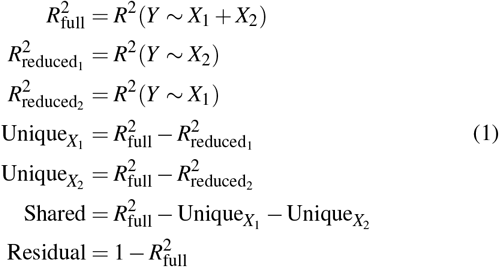

Any negative values arising from this decomposition due to noise or multicollinearity were set to zero for interpretability. This approach provides a formal quantification of the unique and shared variance attributable to each representational model, allowing us to assess the degree of orthogonality between theoretical representational geometries.

The variance partitioning analysis revealed that the three RDMs are well-separated and largely independent (see Supplementary Fig. 1), with each predictor explaining predominantly unique rather than shared variance. This orthogonality validates the theoretical distinctiveness of the representational structures captured by the *Check, Strategy*, and *Visual* RDMs, indicating that they reflect genuinely different aspects of chess board representation. The predominance of unique variance over shared variance supports the use of these RDMs as independent predictors in subsequent analyses, as they capture complementary rather than redundant information about stimulus similarity structure.

### S2 Behavioural Diagnostics: Task Engagement and Preference Structure

To assess whether expert-novice differences in neural representations reflect genuine differences in representational content rather than artefacts of differential task engagement, we conducted a series of diagnostic analyses on the in-scanner behavioural data and on a pre-scan familiarisation task.

#### Response rate

Both groups responded on the vast majority of trials (Experts: *M* = 0.86, *SD* = 0.20; Novices: *M* = 0.94, *SD* = 0.11; Welch’s *t*(27.8) = −1.63, *p* = .114, Cohen’s *d* = −0.53, 95% CI [−0.19, 0.02], Experts −Novices; *n* = 19 experts after excluding one participant whose button-box responses were not recorded), confirming active engagement throughout the scanning session.

#### Checkmate preference

For consecutive 1-back trials comparing a checkmate board to its visually matched non-checkmate counterpart (total *N* = 610 comparisons; per-subject *M* = 15.6), experts preferentially selected checkmate boards (*M* = 0.86, *SD* = 0.17; *t*(18) = 9.33, *p <* .001, Cohen’s *d* = 2.14, 95% CI for *M* [0.78, 0.95], vs. 50%), whereas novices performed at chance (*M* = 0.51, *SD* = 0.13; *t*(19) = 0.37, *p* = .713, Cohen’s *d* = 0.08, 95% CI for *M* [0.45, 0.57]). The group difference was significant (*t*(33.8) = 7.24, *p <* .001, Cohen’s *d* = 2.34, 95% CI [0.25, 0.45], Experts−Novices).

#### Within-subject transitivity

Both groups showed comparable levels of within-subject transitivity (Experts: *M* = 0.41, *SD* = 0.13; Novices: *M* = 0.38, *SD* = 0.08; Welch’s *t*(29.3) = 0.87, *p* = .389, Cohen’s *d* = 0.28, 95% CI [−0.04, 0.10], Experts − Novices), indicating that novices applied similarly consistent evaluation strategies to experts, albeit based on different criteria.

#### Board preference profiles within visual pairs

To test whether novice preferences are driven by perceptual similarity, we examined whether selection frequencies for checkmate boards and their visually matched non-checkmate counterparts were correlated across the 20 visual pairs. In novices, the group-level within-pair correlation was strongly positive (Pearson’s *r* = 0.86, *p <* .001, 95% CI [0.67, 0.94]), and the mean selection frequencies for checkmate (*M* = 0.544) and non-checkmate (*M* = 0.526) boards were nearly identical. At the individual level, novices showed a mean per-subject within-pair correlation of *M*_*r*_ = 0.67 (*SD* = 0.20; *t*(19) = 14.87, *p <* .001, Cohen’s *d* = 3.33, 95% CI [0.58, 0.77], vs. zero). In experts, the group-level within-pair correlation was negative and significant (Pearson’s *r* = −0.54, *p* = .013, 95% CI [−0.79, −0.13]); experts strongly preferred checkmate boards over their non-checkmate counterparts (*M*_*C*_ = 0.815 vs. *M*_*NC*_ = 0.195). The group difference in per-subject within-pair correlation was large (Welch’s *t*(33.2) = −9.75, *p <* .001, Cohen’s *d* = −3.15, 95% CI [−0.91, −0.60], Experts−Novices).

#### Familiarisation classification accuracy

Prior to scanning, participants completed a self-paced online task in which all 40 boards were presented and participants indicated whether a forced checkmate exists, typing the first move if so (*n* = 38; 19 Experts, 19 Novices; data from two participants were unavailable). Classification accuracy was computed as the proportion of all 40 boards correctly classified as checkmate (responded) or non-checkmate (left empty). Both Experts (*M* = 0.88, *SD* = 0.11; *t*(18) = 15.09, *p <* .001, Cohen’s *d* = 3.46, 95% CI for *M* [0.82, 0.93]) and Novices (*M* = 0.59, *SD* = 0.05; *t*(18) = 8.05, *p <* .001, Cohen’s *d* = 1.85, 95% CI for *M* [0.57, 0.62]) performed above chance (50%), though the group difference was large (Welch’s *t*(25.5) = 10.27, *p <* .001, Cohen’s *d* = 3.33, 95% CI [0.23, 0.34], Experts −Novices). Correct first-move responses on the 20 checkmate boards were analysed separately below as a stimulus-specific skill proxy.

#### Pairwise RDM split-half reliability

To assess the reliability of behavioral RDMs, we computed split-half correlations using repeated random split-half resampling. For each group, participants were randomly divided into two non-overlapping halves (9 per half for Experts, *n* = 19; 10 per half for Novices, *n* = 20), separate RDMs were computed with the same method as described in Sec. 4.7.2, and Spearman correlations were calculated between the lower triangles of the two RDMs across 1,000 random splits. The Spearman-Brown formula was then applied to estimate full-sample reliability. Two-sided empirical *p*-values were computed from the resampled *r*_full_ distribution; for the group difference, we evaluated 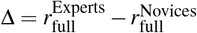 across matched resamples (see Supplementary Table 3). Expert reliability was significant (*r*_full_ = 0.459, *p* = .002, 95% CI [0.392, 0.515]), while Novices did not reach significance (*r*_full_ = −0.040, *p* = .488, 95% CI [0.069, 0.151]). The group difference was significant (Δ*r*_full_ = 0.419, *p* = .002, 95% CI [0.288, 0.536]). Between-group correlations were near zero (*r*_full_ = −0.040, *p* = .589, 95% CI [−0.178, 0.078]), indicating that expert and novice representations are organised differently rather than reflecting a simple strengthening of a common layout.

#### Reconciling pairwise and marginal reliability

The pairwise RDM split-half analysis shows non-significant reliability for novices (*r*_full_ = 0.040, *p* = .488, 95% CI [−0.069, 0.151]), suggesting that their finegrained pairwise dissimilarity structure is not consistent across individuals. However, the marginal (board-level) split-half analysis reveals significant reliability for both groups (Experts: *r*_full_ = 0.928, *p* = .002, 95% CI [0.892, 0.950]; Novices: *r*_full_ = 0.565, *p* = .014, 95% CI [0.175, 0.808]). These two results are not contradictory. The pairwise RDM captures the full 40 × 40 matrix of relative preferences between specific boards; because the 1-back task compares only consecutive stimuli, each specific pair is observed a limited number of times, making the pairwise structure inherently noisy. The marginal preference, by contrast, aggregates each board’s selection frequency across all its pairings, averaging out opponent-specific variability. At this coarser level, novices clearly agree: boards that share visual features with their matched counterpart are preferred at similar rates regardless of checkmate status (within-pair *r* = 0.86, *p <* .001, 95% CI [0.67, 0.94]), and this shared perceptual evaluation is consistent across individuals. The expert-novice difference therefore lies not in the presence or absence of a shared strategy, but in *which dimension* organises that strategy. The board preference feature analysis below confirms this directly: novice preferences are systematically predicted by visual complexity features (officer count, image entropy, edge density), whereas expert preferences are predicted exclusively by checkmate status. This dissociation between reliable marginal and unreliable pairwise structure in novices reflects a deeper property of their evaluative strategy. Expert evaluation is governed by a single categorical feature (checkmate status), which determines the outcome of every head-to-head comparison in the same way, producing consistent pairwise orderings across individuals. Novice evaluation, by contrast, depends on multiple continuous visual features (officer count, image entropy, edge density) that agree on the global board ranking (busier boards are generally preferred) but can conflict on specific pairwise comparisons: two novices may weight different features when resolving a close comparison, leading to divergent pairwise outcomes despite convergent marginal preferences. The pairwise RDM split-half is therefore more sensitive to low-dimensional, categorical strategies than to shared but higher-dimensional continuous strategies. This asymmetry parallels the neural dimensionality finding: expert neural codes are compressed onto fewer, task-relevant dimensions, while novice codes are spread across a higher-dimensional, perceptually-driven feature space.

#### Board preference feature drivers

To identify which objective board properties drive preferences in each group, we extracted 8 features for each of the 40 boards, ordered along a gradient from purely perceptual to deeply relational: (1) image entropy (Shannon entropy of the grayscale pixel intensity histogram, *H* = −∑ *p*_*i*_ log_2_ *p*_*i*_, 256 bins), (2) edge density (proportion of pixels exceeding the 90th percentile of Sobel gradient magnitude), (3) piece count (total pieces on the board), (4) officer count (non-pawn, non-king pieces), (5) centre occupation (pieces within the central 16 squares), (6) king advantage (opponent king exposure minus own king exposure), (7) attack advantage (white attack coverage minus black attack coverage), and (8) checkmate status (binary indicator of forced checkmate). Image-level features (1–2) were extracted from the stimulus images; board-level features (3–8) were computed from the board position via FEN notation using python-chess.

Spearman rank correlations between each feature and mean board selection frequency were computed per group. Significance was assessed via permutation testing (10,000 iterations: feature values randomly shuffled, correlation recomputed; two-tailed *p* = proportion of permuted |*ρ*|≥ observed |*ρ*|). 95% confidence intervals were obtained via bootstrap resampling (10,000 iterations, percentile method). Benjamini-Hochberg FDR correction (*α* = 0.05) was applied to permutation *p*-values across the 8 features within each group. To isolate unique contributions, partial Spearman correlations were computed by residualising both feature and preference on the remaining 7 features via OLS, then correlating residuals (permutation *p*-values and bootstrap CIs computed analogously, re-residualising within each bootstrap/permutation iteration). Explained variance was decomposed via hierarchical linear regression, entering features in three ordered blocks: Perceptual (image entropy, edge density), Structural (piece count, officer count, centre occupation), and Strategic-Relational (king advantage, attack advantage, checkmate status); Δ*R*^2^ per block quantifies unique variance added. These features were not balanced in the stimulus design; results are exploratory.

Across the 40 boards, only checkmate status predicted expert selection frequency (*ρ* = 0.87, 95% CI [0.81, 0.87], *p*_FDR_ *<* .001; Supplementary Fig. 3a); no other feature reached significance (all *p*_FDR_ *>* .38). In novices, three perceptual-structural features survived FDR: officer count (*ρ* = 0.59, 95% CI [0.36, 0.75], *p*_FDR_ = .002), edge density (*ρ* = 0.41, 95% CI [0.14, 0.64], *p*_FDR_ = .032), and image entropy (*ρ* = 0.38, 95% CI [0.12, 0.61], *p*_FDR_ = .049); advantage features and checkmate status were non-significant (Supplementary Fig. 3b). Partial correlations confirmed checkmate status as the sole independent expert predictor (partial *ρ* = 0.94, 95% CI [0.91, 0.99], *p*_FDR_ *<* .001) and officer count as the sole independent novice predictor (partial *ρ* = 0.55, 95% CI [0.27, 0.77], *p*_FDR_ = .006). Image entropy and edge density, significant bivariately, collapsed to non-significant partial correlations (CIs crossing zero), indicating full mediation by piece counts. Hierarchical variance partitioning quantified this dissociation: for experts, the Strategic-Relational block explained 92.7% of variance (Δ*R*^2^ = 0.927; total *R*^2^ = 0.96), with negligible perceptual (0.7%) and structural (2.3%) contributions. For novices, variance was concentrated in the Perceptual (12.8%) and Structural (29.2%) blocks, with only 3.4% from the Strategic-Relational block (total *R*^2^ = 0.45; Supplementary Fig. 3c,d).

This double dissociation is corroborated by the extreme boards in each group’s preference ranking (Supplementary Fig. 2a,c): experts’ three most preferred boards were all single-move checkmate positions (S18, S19, S20; all *f >* 0.89), while their three least preferred were all non-checkmate boards (*f <* 0.12). Novices’ top and bottom boards, by contrast, were a mix of checkmate and non-checkmate positions; notably, the most preferred novice board (S24, NC, *f* = 0.71) and the second most preferred (S4, C, *f* = 0.69) are members of the same visual pair, confirming that novices cannot distinguish between them.

#### Response rate by stimulus condition

To assess whether response behaviour varied systematically across stimulus categories, we computed the proportion of trials on which participants chose the current board (P(choose current)) separately for three trial types: *Advantageous* (the current board is a checkmate position and the previous is not), *Non-advantageous* (the reverse), and *Same* (both boards share the same checkmate status). Total response rates were comparable across conditions for both groups (Supplementary Fig. 5), confirming the absence of condition-specific response biases. Experts showed clear sensitivity to board advantage, preferring the current board on advantageous trials (P(choose current) = 0.87, 95% CI [0.79, 0.95]) and the previous board on non-advantageous trials (P(choose current) = 0.15, 95% CI [0.07, 0.22]). Novices, by contrast, chose near chance across all conditions (P(choose current) = 0.53–0.55; 95% CIs [0.50, 0.56] to [0.52, 0.58]). Because the button-to-response mapping was counterbalanced across runs, any residual association between stimulus category and motor response was balanced out across the data entering the neural analyses.

Together, these results rule out disengagement (high response rates, above-chance familiarisation, comparable transitivity; Supplementary Fig. 4) and support the interpretation that novices were engaged and internally consistent, but relied on a perceptual evaluation strategy that is shared in its broad outlines but noisy in its fine-grained pairwise application. The behavioural dissociation (novices driven by visual similarity, experts by relational structure) directly parallels the neural RSA findings, strengthening the conclusion that the observed neural differences reflect genuine representational reorganisation.

### S3 Brain Decoding

To assess whether local voxel patterns encoded linearly decodable information about chess-relevant categories, we conducted multivariate classification analyses using support vector machines (SVMs). These analyses evaluated the discriminability of activation patterns across different conditions within each region of interest (ROI), providing a functional readout of category-selective information.

Classification analyses were implemented in MATLAB R2024b using the CoSMoMVPA toolbox ^1^. As explained in Sec. 4.6, we used the unsmoothed beta estimates of the 40 conditions as the input of these analyses. For each subject and run, multi-voxel patterns were extracted from predefined ROIs, including bilateral cortical mask from the Glasser atlas ^2^ (See section 4.5 for more details). All analyses were performed separately for each ROI and participant.

For each ROI, a linear SVM classifier (`©cosmo_classify_svm`) was trained to discriminate between condition labels derived from the experimental design. Classification targets were constructed from categorical regressors: Checkmate (2 classes: checkmate vs. non-checkmate), Strategy (10 classes: 5 checkmate-specific and 5 non-checkmate-specific strategy types), and Visual similarity (20 classes: one per matched pair). The data were partitioned using an *n*-fold crossvalidation scheme across runs (leave-one-run-out), with balanced folds ensured via `cosmo_balance_partitions`. Within each fold, voxel patterns were labeled with their corresponding class and used to train and test the classifier.

Decoding accuracy was defined as the mean classification performance across cross-validation folds. ROIs with fewer than 6 usable voxels were excluded to ensure minimum feature dimensionality. Accuracy scores were computed independently for each participant, regressor, and ROI.

Supplementary Table 5 and Supplementary Fig. 6 report SVM decoding accuracy across ROIs for each main model dimension. Decoding based on the *Visual Similarity* model yielded a restricted set of group differences, with significant effects limited to paracentral/midcingulate (Δ*acc* = .013, *t*(34.0) = 2.73, *p*_FDR_ = .044, Cohen’s *d* = 0.86, 95% CI [.003,.023]), premotor (Δ*acc* = .013, *t*(33.8) = 2.87, *p*_FDR_ = .038, Cohen’s *d* = 0.91, 95% CI [.004,.022]), auditory association (Δ*acc* = .015, *t*(36.0) = 3.61, *p*_FDR_ = .007, Cohen’s *d* = 1.14, 95% CI [.006,.023]), superior parietal (Δ*acc* = .028, *t*(36.1) = 3.84, *p*_FDR_ = .005, Cohen’s *d* = 1.21, 95% CI [.013,.042]), and posterior cingulate regions (Δ*acc* = .028, *t*(35.2) = 4.64, *p*_FDR_ = .001, Cohen’s *d* = 1.47, 95% CI [.015,.040]). The majority of visual and high-level areas did not exhibit significant expert-novice differences for this perceptual dimension, and decoding in early and primary visual cortex remained low and non-significant.

In contrast, decoding of the *Strategy* dimension (accuracy −chance) revealed robust group-level differences (Experts −Novices) across a broad network of ROIs. Significant effects included dorsal stream visual cortex (Δ*acc* = .018, *t*(35.1) = 2.87, *p*_FDR_ = .015, Cohen’s *d* = 0.91, 95% CI [.005,.031]), paracentral lobule/mid-cingulate cortex (Δ*acc* = .024, *t*(35.9) = 4.12, *p*_FDR_ *<* .001, Cohen’s *d* = 1.30, 95% CI [.012,.036]), premotor cortex (Δ*acc* = .033, *t*(26.9) = 5.10, *p*_FDR_ *<* .001, Cohen’s *d* = 1.61, 95% CI [.020,.046]), insular and frontal opercular cortex (Δ*acc* = .013, *t*(36.6) = 2.71, *p*_FDR_ = .019, Cohen’s *d* = 0.86, 95% CI [.003,.023]), lateral temporal cortex (Δ*acc* = .016, *t*(33.5) = 3.55, *p*_FDR_ = .003, Cohen’s *d* = 1.12, 95% CI [.007,.026]), temporo-parieto-occipital junction (Δ*acc* = .022, *t*(33.3) = 3.73, *p*_FDR_ = .002, Cohen’s *d* = 1.18, 95% CI [.010,.034]), superior parietal cortex (Δ*acc* = .046, *t*(32.9) = 6.72, *p*_FDR_ *<* .001, Cohen’s *d* = 2.13, 95% CI [.032,.060]), inferior parietal cortex (Δ*acc* = .035, *t*(32.3) = 5.13, *p*_FDR_ *<* .001, Cohen’s *d* = 1.62, 95% CI [.021,.049]), posterior cingulate cortex (Δ*acc* = .026, *t*(29.9) = 3.86, *p*_FDR_ = .002, Cohen’s *d* = 1.22, 95% CI [.012,.039]), anterior cingulate/medial prefrontal cortex (Δ*acc* = .015, *t*(35.1) = 2.84, *p*_FDR_ = .015, Cohen’s *d* = 0.90, 95% CI [.004,.025]), inferior frontal cortex (Δ*acc* = .021, *t*(37.4) = 3.39, *p*_FDR_ = .004, Cohen’s *d* = 1.07, 95% CI [.008,.033]), and dorsolateral prefrontal cortex (Δ*acc* = .027, *t*(31.4) = 4.33, *p*_FDR_ *<* .001, Cohen’s *d* = 1.37, 95% CI [.014,.040]). This pattern indicates that strategy-related distinctions were more robustly represented across higher-order association cortices in Experts than in Novices.

**Supplementary Figure 2.**
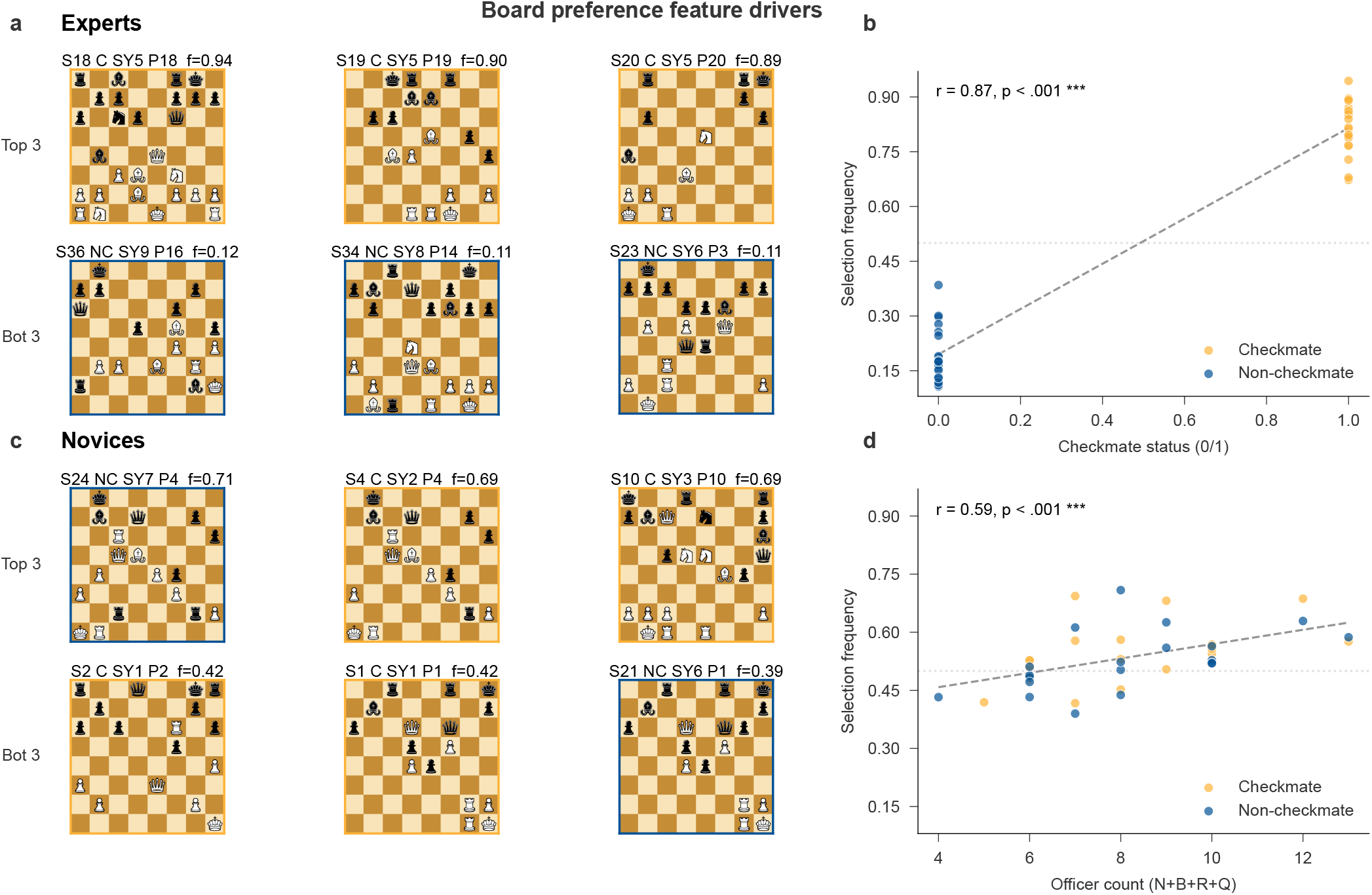
Board preference feature drivers. **a**. Three most and least preferred boards for experts, with selection frequencies (*f*). All top boards are checkmate positions; all bottom boards are non-checkmate. **b**. Expert selection frequency plotted against checkmate status (Spearman’s *ρ* = 0.87, *p*_FDR_ *<* .001, 95% CI [0.81, 0.87]; *n* = 40 boards). **c**. Three most and least preferred boards for novices. Top and bottom boards are a mix of checkmate (orange labels) and non-checkmate (blue labels). **d**. Novice selection frequency plotted against officer count (number of knights, bishops, rooks, and queens; Spearman’s *ρ* = 0.59, *p*_FDR_ = .002, 95% CI [0.36, 0.75]; *n* = 40 boards). Source data are provided as a Source Data file.

**Supplementary Figure 3.**
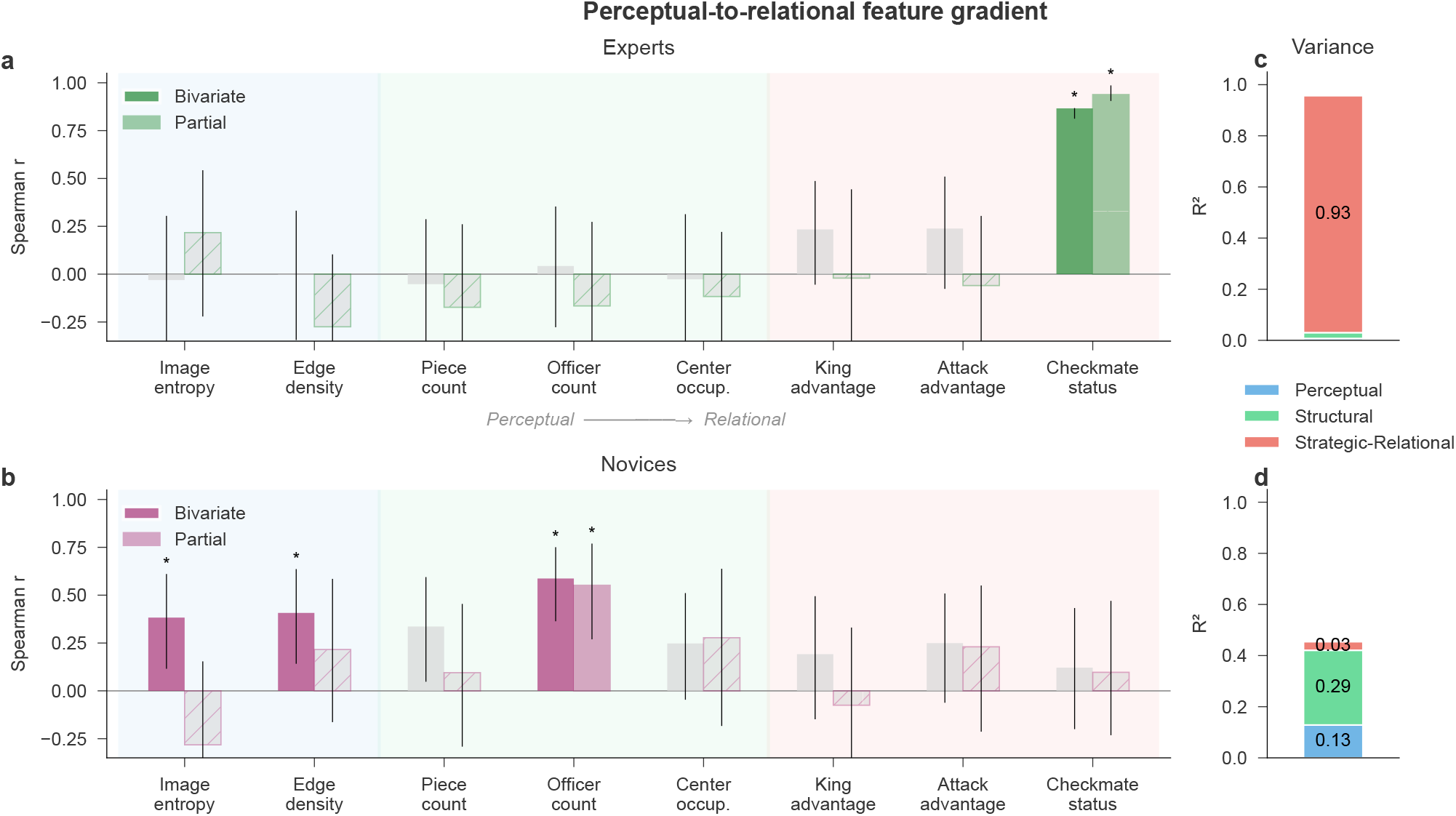
Perceptual-to-relational feature gradient. **a**. Bivariate (solid) and partial (hatched) Spearman correlations between 8 board features and expert selection frequency, with bootstrap 95% CIs (*n* = 40 boards). Background shading indicates the feature category: perceptual (blue), structural (green), strategic-relational (red). Only checkmate status is significant (bivariate and partial). **b**. Same analysis for novices. Officer count, edge density, and image entropy are significant bivariately; only officer count survives partialling. **c**. Hierarchical variance partitioning for experts: 92.7% of variance is explained by the Strategic-Relational block. **d**. For novices, variance is distributed across Perceptual (12.8%) and Structural (29.2%) blocks, with minimal Strategic-Relational contribution (3.4%). Asterisks indicate FDR-corrected significance (*p*_FDR_ *<* .05). Source data are provided as a Source Data file.

**Supplementary Figure 4.**
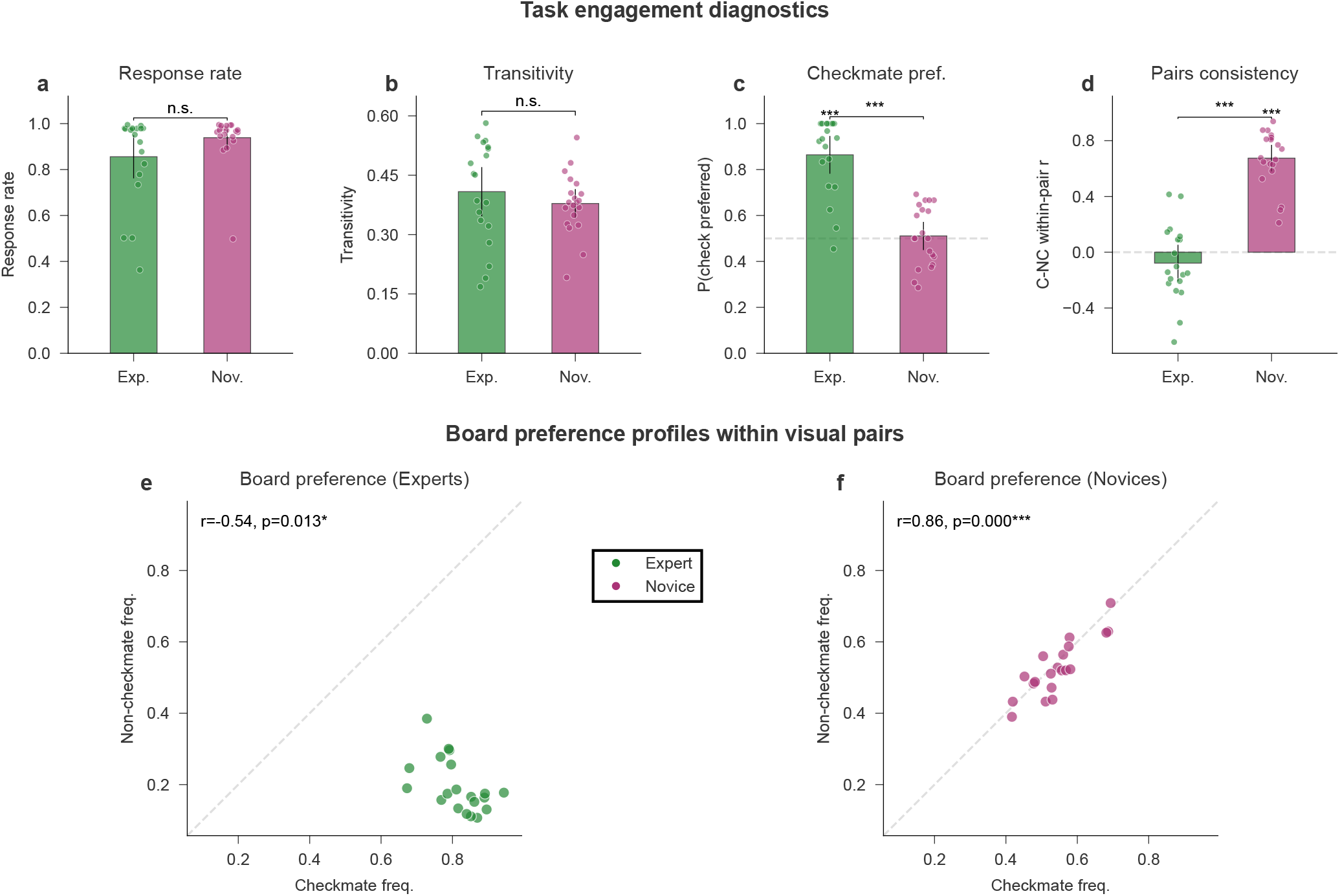
Behavioural diagnostics. Summary of behavioural diagnostic analyses assessing task engagement and preference structure in Experts (*n* = 19 independent participants, excluding one Expert without recorded button-box responses during scanning) and Novices (*n* = 20 independent participants). Top row: per-subject distributions for response rate (**a**), proportion of transitive triples (**b**), checkmate-preference rate (**c**; chance = 0.5), and within-pair checkmate–non-checkmate consistency (**d**; chance = 0); bars show group means with 95% confidence intervals (Student’s *t*-distribution); within-group asterisks above each bar indicate two-sided one-sample *t*-tests against chance for panels c and d; between-group brackets indicate two-sided Welch’s *t*-tests. Bottom row: group-aggregated checkmate vs. non-checkmate selection frequencies for matched stimulus pairs in Experts (**e**) and Novices (**f**); *r* and *p* refer to Pearson correlations across the 20 pairs. Inferential statistics, exact *p*-values, and the parallel response-rate-by-condition analysis are reported in Supplementary Sec. S2 and Supplementary Fig. 5. Source data are provided as a Source Data file.

**Supplementary Figure 5.**
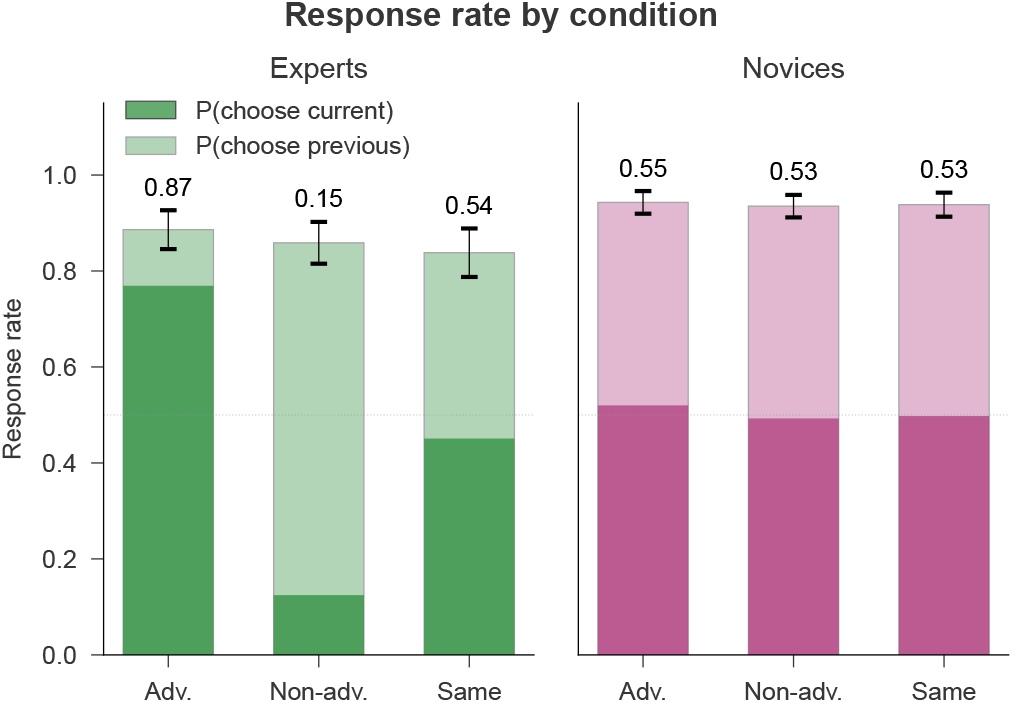
Response rate by stimulus condition. Stacked bars show the per-subject proportion of trials on which participants chose the current board (dark) or the previous board (light), shown for Experts (left, green; *n* = 19, excluding one participant without recorded button-box responses) and Novices (right, magenta; *n* = 20). Conditions reflect the relative advantage of the current board: Advantageous (current is checkmate, previous is not), Non-advantageous (reverse), and Same (both share checkmate status). Annotations indicate P(choose current). Error bars represent 95% confidence intervals on total response rate. Source data are provided as a Source Data file.

Decoding of *Checkmate vs. Non-checkmate* boards (accuracy −chance) yielded a similar spatial profile. Significant group differences (Experts −Novices) were observed in dorsal stream visual cortex (Δ*acc* = .025, *t*(37.9) = 2.47, *p*_FDR_ = .033, Cohen’s *d* = 0.78, 95% CI [.005,.046]), somatosensory/motor cortex (Δ*acc* = .033, *t*(27.5) = 2.64, *p*_FDR_ = .030, Cohen’s *d* = 0.83, 95% CI [.007,.059]), paracentral lobule/mid-cingulate cortex (Δ*acc* = .036, *t*(36.1) = 2.89, *p*_FDR_ = .016, Cohen’s *d* = 0.92, 95% CI [.011,.062]), premotor cortex (Δ*acc* = .052, *t*(25.7) = 4.35, *p*_FDR_ *<* .001, Cohen’s *d* = 1.38, 95% CI [.027,.077]), lateral temporal cortex (Δ*acc* = .052, *t*(35.1) = 5.22, *p*_FDR_ *<* .001, Cohen’s *d* = 1.65, 95% CI [.032,.072]), temporo-parieto-occipital junction (Δ*acc* = .028, *t*(37.3) = 2.48, *p*_FDR_ = .033, Cohen’s *d* = 0.78, 95% CI [.005,.052]), superior parietal cortex (Δ*acc* = .053, *t*(37.9) = 4.60, *p*_FDR_ *<* .001, Cohen’s *d* = 1.46, 95% CI [.030,.077]), inferior parietal cortex (Δ*acc* = .089, *t*(31.5) = 6.90, *p*_FDR_ *<* .001, Cohen’s *d* = 2.18, 95% CI [.063,.116]), posterior cingulate cortex (Δ*acc* = .035, *t*(26.6) = 3.55, *p*_FDR_ = .004, Cohen’s *d* = 1.12, 95% CI [.015,.056]), anterior cingulate/medial prefrontal cortex (Δ*acc* = .047, *t*(28.4) = 3.96, *p*_FDR_ = .002, Cohen’s *d* = 1.25, 95% CI [.023,.071]), orbital and polar frontal cortex (Δ*acc* = .026, *t*(30.9) = 2.33, *p*_FDR_ = .045, Cohen’s *d* = 0.74, 95% CI [.003,.048]), inferior frontal cortex (Δ*acc* = .053, *t*(29.1) = 3.88, *p*_FDR_ = .002, Cohen’s *d* = 1.23, 95% CI [.025,.081]), and dorsolateral prefrontal cortex (Δ*acc* = .080, *t*(35.4) = 5.62, *p*_FDR_ *<* .001, Cohen’s *d* = 1.78, 95% CI [.051,.108]). These effects indicate that goal-related distinctions such as checkmate status were more strongly represented in higher-order associative cortices for Experts than for Novices.

Aside our main decoding analysis (i.e., on the three main dimensions built into our dataset), we also performed a finer decoding analysis on five additional possible categorisations of the checkmate boards. This finer analysis allows us to assess what information can be decoded within the checkmate class, offering a proxy into how strategic information may be structured. Specifically, we performed a decoding analysis on: Strategy (the same as in the main analysis, but now using only the 20 checkmate chess boards), Motif, Total number of pieces, Total number of available legal moves, and Number of white moves to checkmate. All the results for this analysis (and their RSA counterparts) are reported in Supplementary Tables 7 and 6, and visualized in Supplementary Fig. 7.

### S4 Region-level interpretation of the dimensionality classifier

For most regions, the classifier weights in Fig. 5C converge with the findings from the PR analysis in Fig. 6: a clearly negative or positive classifier weight in Fig. 5C corresponds to a (significant) negative or positive group difference in PR in Fig. 6, respectively. A notable exception is the auditory cortex.

Functionally, the prominence of auditory cortex in this classifier emerges from a coherent secondary axis of the expertise effect rather than from any single region in isolation. The classifier weight on auditory cortex is positive (Early Auditory *w* = +0.41; Auditory Association *w* = +0.72, the third-largest absolute weight in Fig. 5C), meaning that higher PR in auditory cortex shifts the classifier toward an expert prediction. This direction is consistent with the per-ROI means: experts trend toward higher PR than novices in Early Auditory (ΔPR = +1.87) and Auditory Association (ΔPR = +0.26), although neither survives FDR correction at the per-region level (Fig. 6, Supplementary Table 8). The same anatomical inversion is recovered independently by the unsupervised PCA decomposition (Fig. 5A, bottom): PC1 (52.6% of variance) carries near-uniform positive loadings on the visual, parietal, and lateral prefrontal regions where experts compress, while PC2 (13.1% of variance) is bipolar, with positive loadings concentrated in insular and frontal opercular cortex (the largest PC2 loading, +0.49), auditory cortex (Early Auditory +0.43; Auditory Association +0.26), medial temporal cortex (+0.39), and posterior opercular cortex (+0.26). The single FDR-significant region in which experts show higher PR than novices — the insular and frontal opercular cortex (ΔPR = +4.49, *t*(37.3) = +3.07, *p*_FDR_ = .015, Cohen’s *d* = +0.97, 95% CI [+1.53, +7.44]) — is the strongest member of this same set; auditory cortex follows the same direction at smaller magnitude.

This secondary axis spans regions implicated in language and inner-speech (auditory, insular, frontal opercular), memory retrieval (medial temporal), and multi-sensory integration (posterior opercular). Reading our data conservatively, expert chessboard evaluation appears to engage these regions in a more differentiated, higher-dimensional regime — complementing the dominant compression we report in visual, parietal, and prefrontal cortex — even though the canonical chess task contains no auditory stimulation. The cognitive mechanism behind this secondary axis is not adjudicated by the present design (candidates compatible with the chess-expertise literature include verbal labelling of strategic patterns during evaluation and multi-sensory recruitment in long-term experts), and we therefore flag it as an observation that warrants targeted follow-up rather than as direct evidence for any specific account.

### S5 Participation ratio is not predicted by ROI size

To assess how regional dimensionality relates to anatomical size, we computed the participation ratio (PR; ^3,4^) within each cortical ROI for each participant. We then averaged PR values across participants within each group (Experts and Non-Experts) and calculated the difference (Expert minus Non-Expert) per ROI. Finally, we visualized and quantified the relationship between PR and ROI size by plotting these measures against each other and computing Pearson correlation coefficients across the 22 cortical ROIs (see Fig. 8). PR did not significantly scale with ROI size in Experts (*r* = 0.18, *p* = .426, 95% CI [−0.26, 0.56]), Novices (*r* = 0.33, *p* = .135, 95% CI [−0.11, 0.66]), or the Expert −Novice difference (*r* = −0.15, *p* = .506, 95% CI [−0.54, 0.29]), indicating that the reported PR effects are not explained by anatomical ROI extent.

### S6 Skill-Gradient Analysis

To assess whether the observed neural differences scale continuously with skill, we tested for graded relationships between skill measures and key neural metrics within the expert group and across all participants. For Elo analyses, we computed both Pearson and Spearman correlations. For familiarisation accuracy, we used Pearson correlations with neural metrics averaged across the 22 ROI groups. The “mean” neural metrics reported below were obtained by averaging each measure (PR, RSA model fit, or decoding accuracy) across the 22 bilateral Glasser ROI groups before correlating with skill. We also tested correlations at the individual ROI level (22 tests per measure); these were corrected for multiple comparisons using the Benjamini-Hochberg FDR procedure at *α* = 0.05. No individual ROI survived FDR correction for any measure.

#### Elo and neural measures (experts only)

Within the expert group (*n* = 20, Elo range: 1751–2269), Elo rating correlated with mean Checkmate RSA model fit (Pearson’s *r* = 0.47, *p* = .039, 95% CI [0.03, 0.75]; Spearman’s *ρ* = 0.43, *p* = .061, 95% CI [−0.02, 0.73]). Ten of 22 individual ROIs showed uncorrected positive trends, but no individual ROI survived FDR correction. Neither mean participation ratio (Pearson’s *r* = −0.03, *p* = .913, 95% CI [−0.46, 0.42]), Strategy RSA (Pearson’s *r* = 0.37, *p* = .113, 95% CI [−0.09, 0.70]), nor decoding accuracy showed significant Elo correlations. The restricted Elo range (518 points) limits statistical power for detecting continuous effects within this group.

#### Familiarisation move accuracy and neural measures (all participants)

Across the full sample (*n* = 38), familiarisation move accuracy on the 20 checkmate boards served as a stimulus-specific skill proxy, directly indexing how well each participant could evaluate the exact boards presented during fMRI. Move accuracy correlated strongly with checkmate decoding (Pearson’s *r* = 0.71, *p*_FDR_ *<* .001, 95% CI [0.50, 0.84]), strategy decoding (Pearson’s *r* = 0.54, *p*_FDR_ *<* .001, 95% CI [0.27, 0.73]), strategy RSA (Pearson’s *r* = 0.59, *p*_FDR_ *<* .001, 95% CI [0.33, 0.76]), checkmate RSA (Pearson’s *r* = 0.44, *p*_FDR_ = .008, 95% CI [0.14, 0.67]), and mean participation ratio (Pearson’s *r* = −0.39, *p*_FDR_ = .017, 95% CI [−0.63, −0.08]; negative because higher skill is associated with lower dimensionality). No correlation was observed with visual similarity RSA (Pearson’s *r* = −0.13, *p*_FDR_ = .431, 95% CI [−0.43, 0.20]), confirming that the skill gradient tracks relational but not perceptual neural structure. Within the expert group alone (*n* = 19), none of these correlations reached significance after FDR correction, consistent with the restricted skill range.

These results demonstrate robust skill-to-representation relationships at the group level, with a detectable trend within experts for Elo and Checkmate RSA (Supplementary Fig. 9). The convergence of Elo (a general, externally validated skill measure) and familiarisation accuracy (a stimulus-specific skill proxy) supports the interpretation that the neural reorganisation scales with chess-relevant competence rather than reflecting a categorical group difference.

**Supplementary Figure 6.**
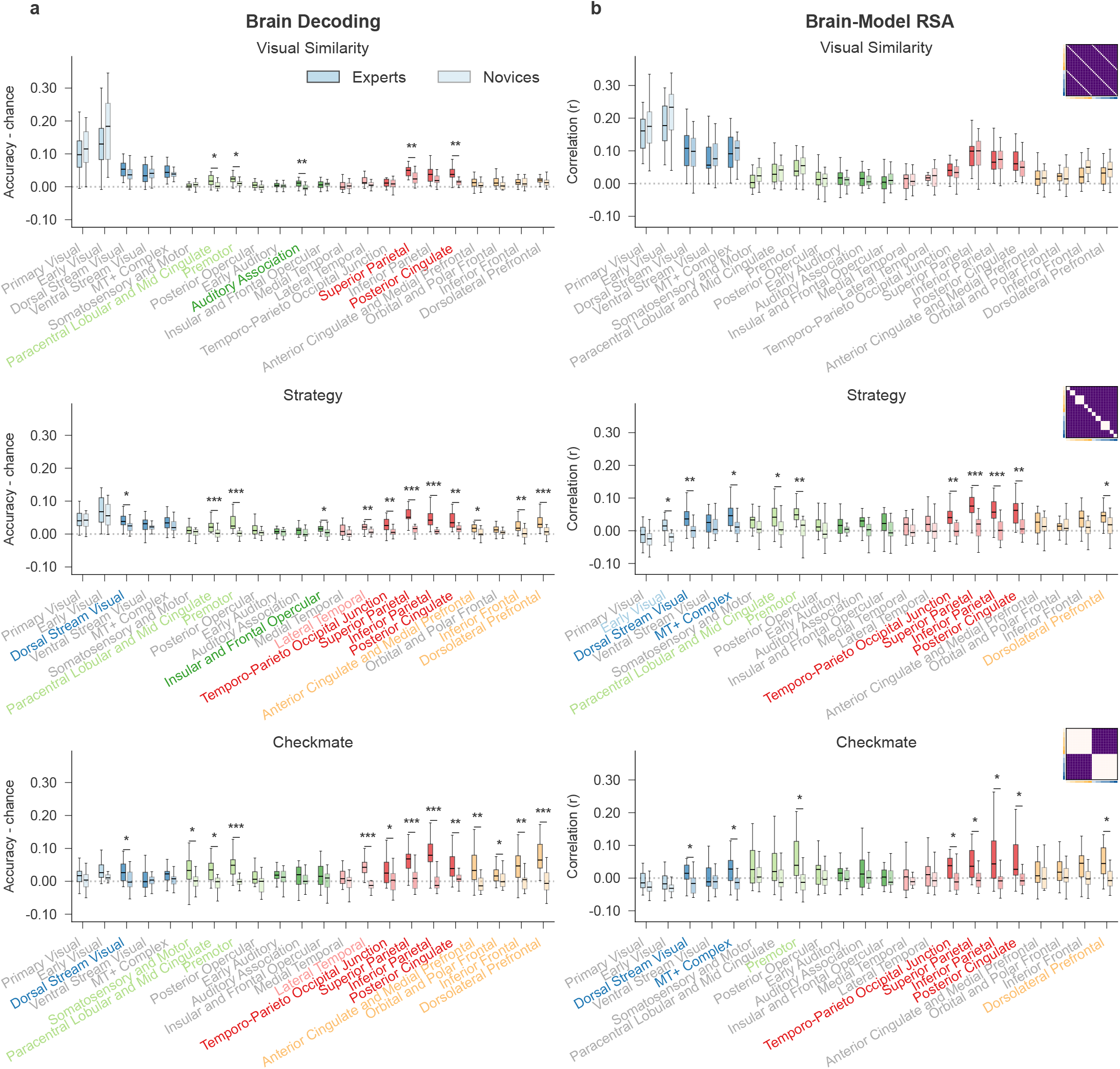
Comparison of decoding (a) and RSA (b) on the main model dimensions. ROI-wise summaries of model-brain correspondence (RSA) versus chance-corrected linear classification accuracy (SVM decoding; accuracy minus chance) for the three primary dimensions (Checkmate, Strategy, Visual similarity). Box plots show the distribution of per-subject values for Experts (*n* = 20 independent participants, solid) and Novices (*n* = 20 independent participants, lighter shade). Boxes span the interquartile range (IQR; 25th–75th percentile), with the median marked by a horizontal line; whiskers extend to the furthest data point within 1.5 × IQR of the box edges. Colours indicate ROI groups. Stars indicate two-sided Welch’s *t*-test (`scipy.stats.ttest_ind` with `equal_var=False`) Experts vs. Novices, FDR-corrected (Benjamini–Hochberg) across the 22 ROIs separately within each model: ^∗^ *p*_FDR_ *<* .05, ^∗∗^ *p*_FDR_ *<* .01, ^∗∗∗^ *p*_FDR_ *<* .001. Per-ROI *t*-statistics, Welch–Satterthwaite degrees of freedom, mean differences, 95% confidence intervals, Cohen’s *d*, and exact raw and FDR-corrected *p*-values are reported in Supplementary Tables 4 and 5. Source data are provided as a Source Data file.

**Supplementary Figure 7.**
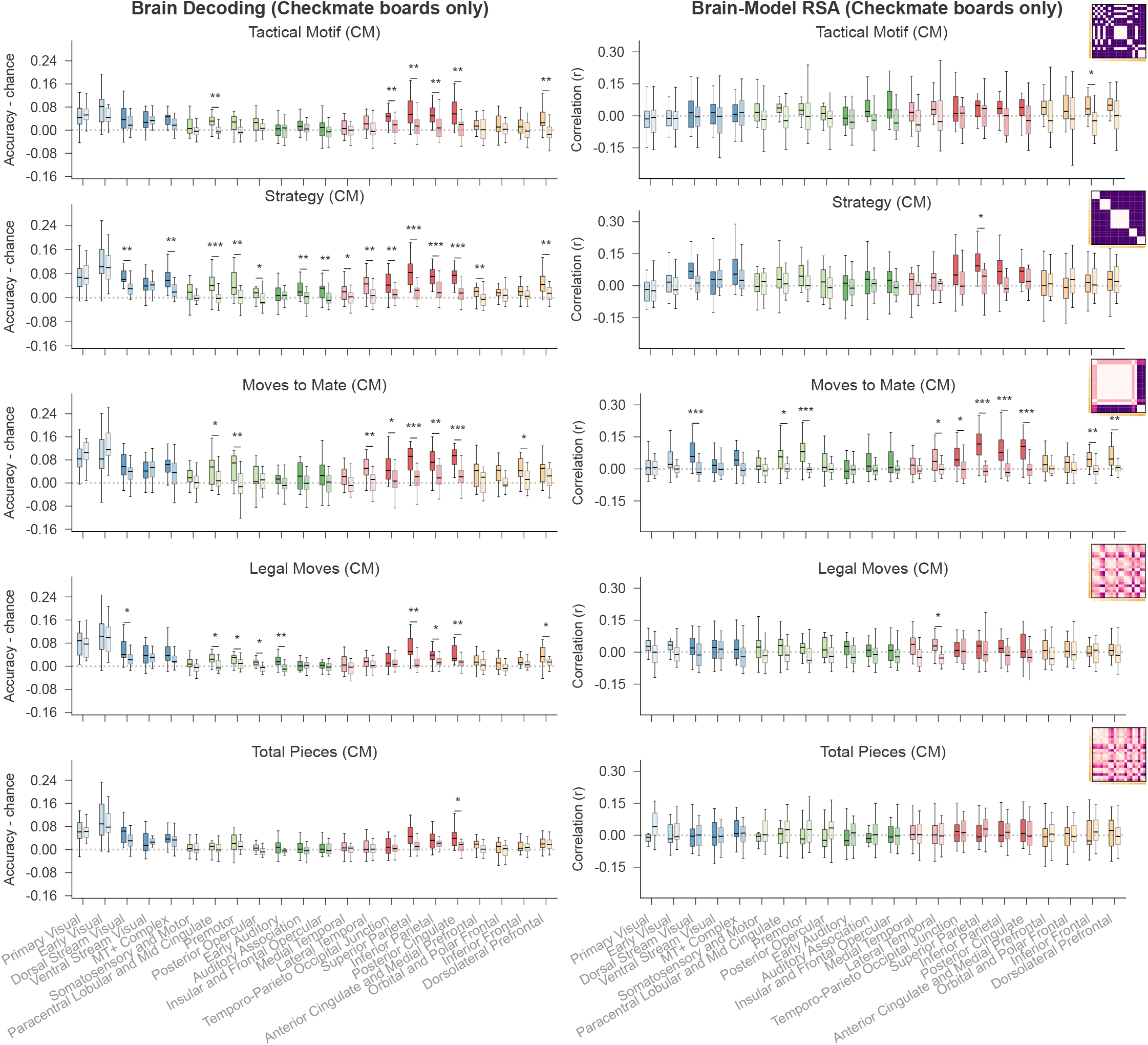
Comparison of RSA and decoding on finer dimensions (checkmate stimuli only). As in Supplementary Fig. 6, but restricted to checkmate boards and extended to finer categorisations (Strategy within checkmates, Motif, Total pieces, Legal moves, and White moves to mate). Per-subject distributions for Experts (*n* = 20 independent participants, solid) and Novices (*n* = 20 independent participants, lighter shade) are shown as box plots (IQR boxes, median centre line, whiskers within 1.5 × IQR of the box edges). RDMs derived from the finer dimensions are shown on the right. Stars indicate two-sided Welch’s *t*-test Experts vs. Novices, FDR-corrected (Benjamini–Hochberg) across the 22 ROIs separately within each model: ^∗^ *p*_FDR_ *<* .05, ^∗∗^ *p*_FDR_ *<* .01, ^∗∗∗^ *p*_FDR_ *<* .001. Per-ROI test statistics, degrees of freedom, mean differences, 95% confidence intervals, Cohen’s *d*, and exact *p*-values are reported in Supplementary Tables 6 and 7. Source data are provided as a Source Data file.

**Supplementary Figure 8.**
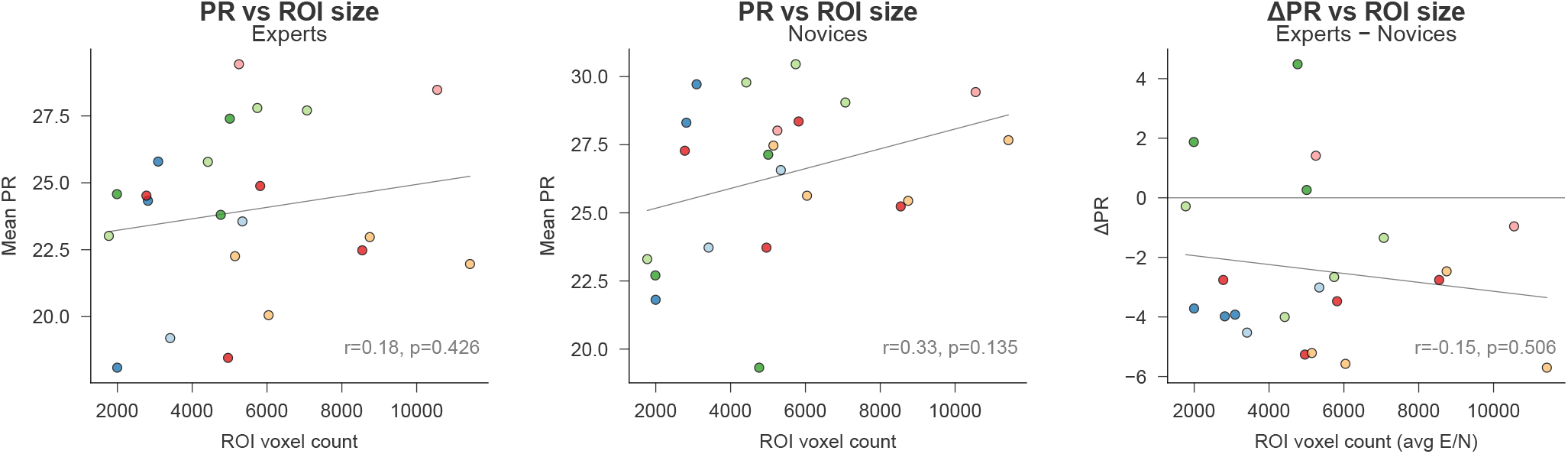
**Relationship between PR and ROI size** (number of voxels), plotted separately for Experts (left), Novices (centre), and group differences (right). Each dot represents one ROI (*n* = 22 cortical ROIs); lines indicate linear regression fits. Pearson correlations were small and non-significant for Experts (*r* = 0.18, *p* = .426, 95% CI [−0.26, 0.56]), Novices (*r* = 0.33, *p* = .135, 95% CI [−0.11, 0.66]), and group differences (*r* = −0.15, *p* = .506, 95% CI [−0.54, 0.29]). These analyses test whether PR scales with anatomical extent and whether this relationship differs between groups. Source data are provided as a Source Data file.

### S7 Groups cannot be inferred from estimated eye-movements

To investigate whether estimated eye-movement patterns differ systematically between chess Experts and Novices, we performed two separate decoding analyses using support vector machines (SVM). The first analysis used the estimated gaze coordinates (x, y), while the second used the corresponding displacements from the centre of the screen. These estimates were derived from functional MRI data using BidsMReye (), a BIDS-ready version of DeepMReye ^5^.

We included the full sample of 40 participants (20 Experts and 20 Novices). For each participant, all available task runs were used; each run was treated as an independent sample. The input features for each sample were constructed by flattening the time series of the gaze coordinates or displacement values within a run, resulting in a subject-run-level feature vector. This yielded a feature matrix *X* ∈ ℝ^*n*×*d*^, where *n* is the number of runs and *d* is the number of timepoints times two (for x and y) or one (for displacement). Binary class labels *y* ∈ {0, 1}^*n*^ indicated expertise (expert = 1, novice = 0), and subject identifiers were recorded to define cross-validation groups.

We assessed classification performance using stratified group *k*-fold cross-validation with *k* = 20 folds, as implemented in `StratifiedGroupKFold` from `scikit-learn`. This procedure ensured that samples from the same participant were never split between training and test sets (i.e., we used 19 subjects per group as training set and one subject per group as test set). Folds were stratified by class to preserve expert/novice balance.

Within each fold, we trained a support vector classifier with a linear kernel using a pipeline that included z-scoring of input features via `StandardScaler`. The model was trained on all runs from all training participants and evaluated on the runs from the held-out participants. For each test sample, we recorded the predicted class label and the posterior probability for being classified as an expert. This process was repeated for all folds, and the predicted labels and probabilities were aggregated across folds.

For each participant, runs from all folds in which that participant appeared in the test set were aggregated to produce a single subject-level out-of-fold accuracy. Statistical significance was assessed with a two-sided one-sample *t*-test on these 40 subject-level accuracies against a null hypothesis of 0.5 (chance). Pooled run-level accuracy with Wilson 95% confidence intervals is reported descriptively. Neither feature set yielded significant decoding: for gaze coordinates, mean subject-level accuracy was 0.534 (95% CI [0.475, 0.594], *t*(39) = 1.162, *p* = .252, Cohen’s *d* = 0.18); for displacement, 0.462 (95% CI [0.415, 0.509], *t*(39) = −1.620, *p* = .113, Cohen’s *d* = −0.26). Results are reported in Supplementary Fig. 10 and Supplementary Table 14.

### S8 Subcortical Representational and Decoding Analysis

To test whether the expertise-related representational effects observed in cortex extend to subcortical structures, we performed an exploratory analysis using the Cole-Anticevic Brain-wide Network Partition (CABNP; ^6^). We extracted the 358 subcortical parcels from the CAB-NP CIFTI atlas, grouped them into 9 bilateral anatomical ROIs (Hippocampus, Amygdala, Caudate, Putamen, Pallidum, Thalamus, Nucleus Accumbens, Cerebellum, Brainstem), and resampled the resulting atlas into MNI volumetric space (2 mm isotropic) using nearest-neighbour interpolation. Voxels overlapping with the cortical Glasser atlas were excluded to ensure strictly non-overlapping cortical and subcortical regions.

We then applied the same multivariate pipeline used for cortical ROIs (Sec. 4.7.3 and Supplementary Sec. S3): RSA correlations with the three model RDMs (Checkmate, Strategy, Visual Similarity) and SVM decoding of the same three dimensions, with identical cross-validation, balancing, and statistical procedures. Group differences were tested with Welch’s *t*-tests, FDR-corrected across the 9 subcortical ROIs.

No subcortical ROI showed a significant expert-novice difference in either RSA or decoding after FDR correction (Supplementary Fig. 11; smallest *p*_FDR_ = .090). Uncorrected trends were observed in the cerebellum for checkmate decoding (Δ*M* = 0.035, *t*(27.7) = 2.59, *p* = .015, *p*_FDR_ = .135, Cohen’s *d* = 0.82, 95% CI [0.007, 0.063]) and strategy decoding (Δ*M* = 0.013, *t*(35.1) = 2.72, *p* = .010, *p*_FDR_ = .090, Cohen’s *d* = 0.86, 95% CI [0.003, 0.023]), but neither survived correction for multiple comparisons.

These results suggest that the expertise-related representational reorganisation characterised in the main analyses is predominantly cortical. The null result in subcortical regions does not rule out subcortical contributions to chess expertise, but it indicates that the representational effects detectable with the present sample, task, and analytical approach are concentrated in cortical systems. Subcortical involvement may be more closely tied to procedural learning, memory consolidation, or reward processing, aspects that the current task (board evaluation rather than active play) may not strongly engage.

**Supplementary Figure 9.**
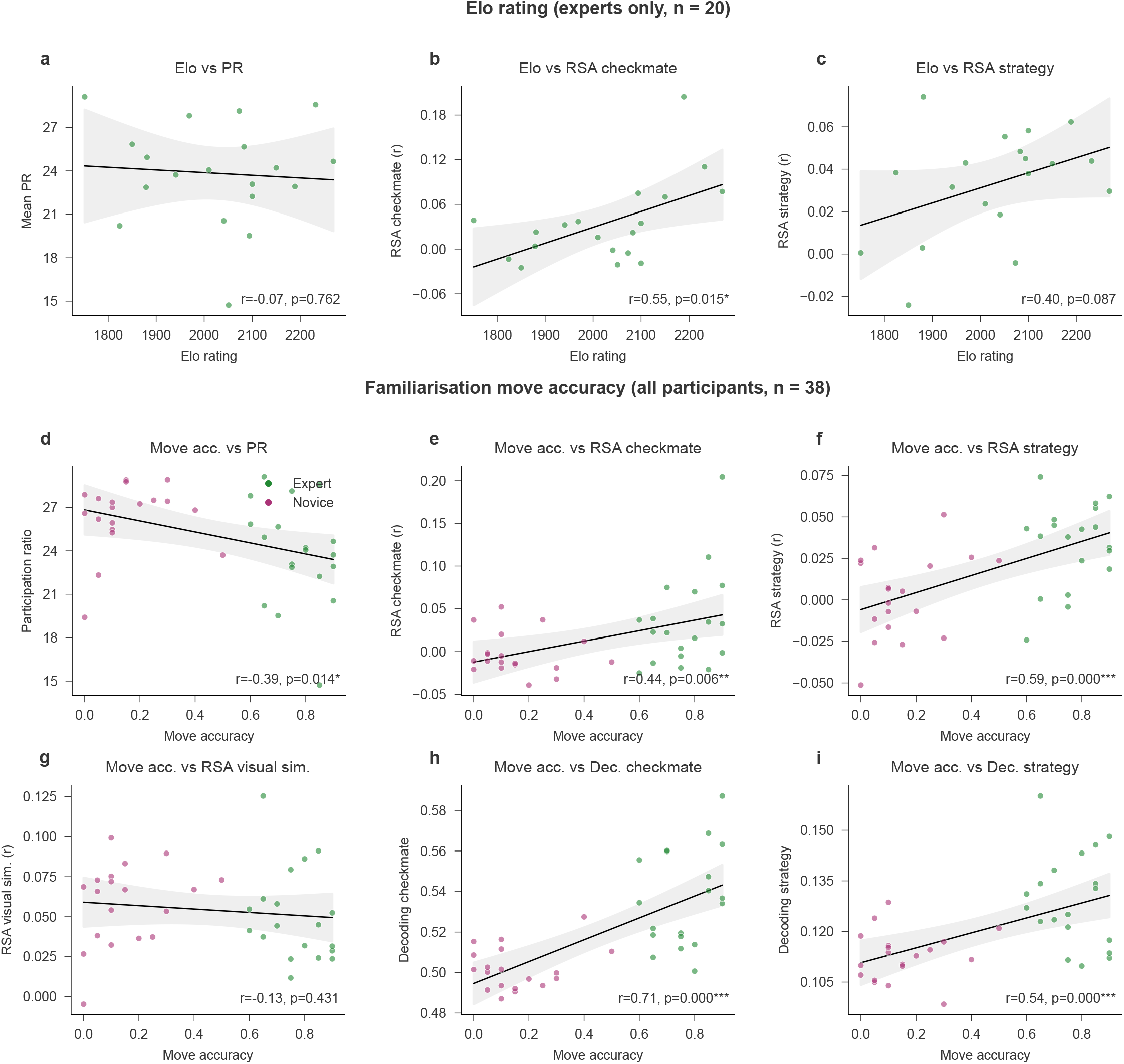
Skill-gradient analysis. Correlations between skill measures and neural metrics. **a**–**c**. Pearson correlations between Elo rating and three neural metrics in experts only (*n* = 20 independent participants): mean participation ratio (**a**), RSA checkmate fit (**b**), and RSA strategy fit (**c**). **d**–**i**. Pearson correlations between familiarisation move accuracy and six neural metrics across all participants who completed the online familiarisation (*n* = 38 independent participants): mean participation ratio (**d**), RSA checkmate fit (**e**), RSA strategy fit (**f**), RSA visual-similarity fit (**g**), checkmate decoding accuracy (**h**), and strategy decoding accuracy (**i**). Each point represents one participant; lines indicate linear fits with 95% confidence bands. Statistical tests are two-sided Pearson correlations with 95% confidence intervals computed by 10,000-resample bootstrap; *p*-values are FDR-corrected (Benjamini–Hochberg) across the 22 ROIs within each measure (no individual ROI survived FDR correction). Source data are provided as a Source Data file.

**Supplementary Figure 10.**
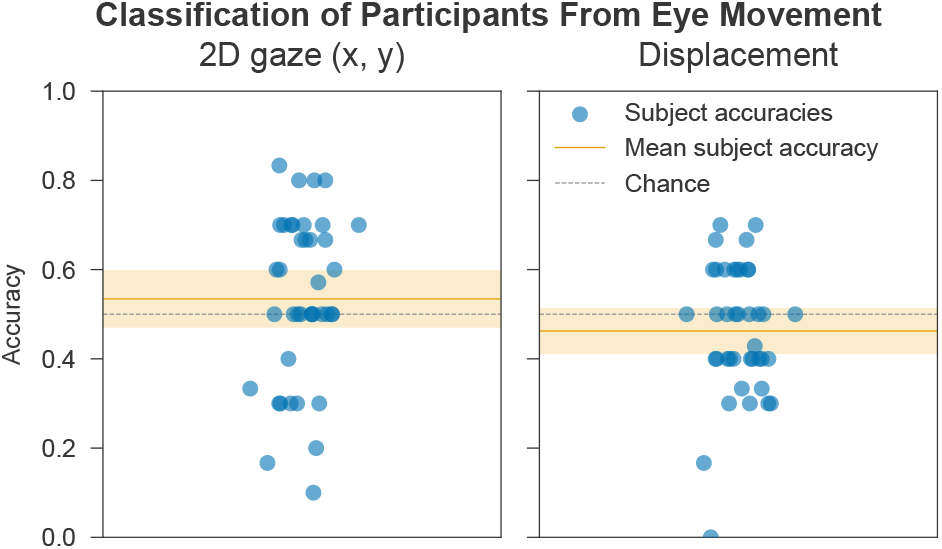
Decoding chess expertise from eye-tracking data. Each dot represents the subject-level out-of-fold accuracy for one participant (*n* = 40 independent participants). **Left:** decoding based on two-dimensional gaze coordinates (x, y). **Right:** decoding based on displacement from screen centre. The solid horizontal line shows the mean subject-level accuracy; the dashed line indicates chance level (0.5). The shaded band marks the 95% confidence interval around the mean. Group-level inference is via a two-sided one-sample *t*-test of subject-level accuracy against chance (0.5); exact *t, d f, p*, Cohen’s *d*, and 95% confidence intervals are reported in Supplementary Table 14. Source data are provided as a Source Data file.

### S9 Stimulus Set Details

This supplementary section provides a comprehensive overview of the 40 chessboard stimuli used in the experiment, each labeled and visualized in Supplementary Fig. 12 and reported in Supplementary Table 1. Stimuli were constructed to systematically vary along three key dimensions: whether they depicted an imminent checkmate (Checkmate vs. Non-checkmate), the tactical strategy leading to the checkmate (Strategy Type), and perceptual similarity (Visual Pairing). The chessboards shown throughout this manuscript and Supplementary Information were rendered from the stimulus positions (Forsyth–Edwards Notation; Supplementary Table 1) using the open-source *chessnut* piece set by Alexis Luengas, distributed under the Apache License 2.0.

*Checkmate* boards (C/NC) depicted configurations in which a forced mate sequence for White against Black existed, resolvable in four or fewer moves. Each checkmate board (denoted “C” in the figure) was paired with a non-checkmate counterpart (“NC”) that preserved the overall structure and piece configuration but introduced a minimal intervention—typically the repositioning of one or two pieces—to neutralize the mating sequence while maintaining close visual similarity.

*Strategy types* (SY) were defined according to both piece involvement and underlying tactical features, reflecting qualitative distinctions in how mating sequences are structured. These include queen-rook combinations (SY1 and SY6), involving coordinated linear tactics; supported attacks with minor pieces (SY2 and SY7), where auxiliary knights or bishops play a crucial role in sustaining the queen mate; minor piece mating nets (SY3 and SY8), relying on spatial confinement by knights and bishops; bishop-driven forcing moves (SY4 and SY9), emphasizing long-range control; and one-move checkmates (SY5 and SY10), which require minimal calculation and serve as structurally simple baselines.

*Visual pairing* (P) designates stimuli matched primarily for perceptual similarity while holding relational structure constant. For each position, we also recorded the total number of pieces, the count of legal moves, the side of the board occupied by the defending king (left = 0, right = 1), and the dominant tactical motif characterising the checkmate sequence. Where applicable, the number of moves required to reach checkmate is also noted.

### S10 Run-Matched Control Analysis

Although the number of functional runs did not differ significantly between groups (Experts: *M* = 8.4, *SD* = 2.0; Novices: *M* = 9.5, *SD* = 1.4; Welch’s *t*(33.3) = −1.94, *p* = .061, Cohen’s *d* = −0.61, 95% CI [−2.15, 0.05], Experts −Novices), novices completed slightly more runs on average. To rule out the possibility that this imbalance contributes to the reported expertise effects, we repeated the ROI-based RSA and participation ratio analyses after equalising the run distribution across groups. All expert data were preserved. For novices, 8 subjects were deterministically assigned a cap of 6 runs (matching the 8 experts who completed 6 runs), yielding identical run distributions in both groups (8 × 6 + 12 × 10 = 168 runs per group). All statistical procedures (Welch *t*-tests, FDR correction) were identical to the main analyses.

The participation ratio results were unchanged: all 14 originally FDR-significant ROIs remained significant, and classification of experts vs. novices from dimensionality profiles remained highly accurate (ROI space: cross-validated accuracy = 80.0%, permutation *p* = .0006; PCA-2D: cross-validated accuracy = 85.0%, permutation *p* = .0001; stratified 5-fold CV, *n* = 40 independent participants, 10,000 label permutations; Supplementary Table 16). For RSA, all originally significant ROIs retained significance, and several ROIs that had been borderline in the original analysis crossed the FDR threshold once the novice data advantage was removed: three additional ROIs for the Checkmate model and two for the Strategy model (Supplementary Table 15). No ROI showed the reverse pattern (significant in the original but not in the run-matched analysis). The Visual Similarity model, which does not differ between groups, remained entirely non-significant in both analyses (0 ROIs). This pattern is expected: because novices contributed more data to the original analysis, their within-group estimates were more precise, producing slightly lower *t*-statistics for the group contrast. Equalising run counts removes this advantage, yielding marginally stronger expert-novice differences. These results confirm that the original expertise effects are, if anything, conservative estimates, and that the slight run imbalance does not inflate or drive the reported findings.

### S11 fMRI data pre-processing

#### Anatomical data preprocessing

A total of 1 T1-weighted (T1w) images was found within the input BIDS dataset per subject. The T1w image was corrected for intensity non-uniformity (INU) with `N4BiasFieldCorrection` ^7^, distributed with ANTs 2.5.1^8^ RRID:SCR_004757, and used as T1w-reference throughout the workflow. The T1w-reference was then skull-stripped with a *Nipype* implementation of the `antsBrainExtraction.sh` workflow (from ANTs), using OASIS30ANTs as target template. Brain tissue segmentation of cerebrospinal fluid (CSF), white-matter (WM) and gray-matter (GM) was performed on the brain-extracted T1w using `fast` ^9^ (RRID:SCR_002823). Brain surfaces were reconstructed using `recon-all` FreeSurfer 7.3.2, RRID:SCR_001847, ^10^, and the brain mask estimated previously was refined with a custom variation of the method to reconcile ANTs-derived and FreeSurfer-derived segmentations of the cortical gray-matter of Mindboggle RRID:SCR_002438, ^11^. Volume-based spatial normalization to one standard space (MNI152NLin2009cAsym) was performed through nonlinear registration with `antsRegistration` (ANTs 2.5.1), using brain-extracted versions of both T1w reference and the T1w template. The following template was selected for spatial normalization and accessed with *TemplateFlow* 24.2.0, ^12^: *ICBM 152 Nonlinear Asymmetrical template version 2009c* ^13^ (RRID:SCR_008796; TemplateFlow ID: `MNI152NLin2009cAsym`).

#### Functional data preprocessing

For each of the BOLD runs found per subject, the following preprocessing was performed. First, a reference volume was generated, using a custom methodology of *fMRIPrep*, for use in head motion correction. Head-motion parameters with respect to the BOLD reference (transformation matrices, and six corresponding rotation and translation parameters) are estimated before any spatiotemporal filtering using `mcflirt` ^14^. The BOLD reference was then co-registered to the T1w reference using `bbregister` (FreeSurfer) which implements boundary-based registration ^15^. Co-registration was configured with six degrees of freedom. Several confounding time-series were calculated based on the *preprocessed BOLD*: framewise displacement (FD), DVARS and three region-wise global signals. FD was computed using two formulations following Power (absolute sum of relative motions, ^16^) and Jenkinson (relative root mean square displacement between affines, ^14^). FD and DVARS are calculated for each functional run, both using their implementations in *Nipype* following the definitions by ^16^. The three global signals are extracted within the CSF, the WM, and the whole-brain masks. Additionally, a set of physiological regressors were extracted to allow for component-based noise correction *CompCor* ^17^. Principal components are estimated after high-pass filtering the *preprocessed BOLD* time-series (using a discrete cosine filter with 128s cut-off) for the two *CompCor* variants: temporal (tCompCor) and anatomical (aCompCor). tCompCor components are then calculated from the top 2% variable voxels within the brain mask. For aCompCor, three probabilistic masks (CSF, WM and combined CSF+WM) are generated in anatomical space. The implementation differs from that of Behzadi et al. in that instead of eroding the masks by 2 pixels on BOLD space, a mask of pixels that likely contain a volume fraction of GM is subtracted from the aCompCor masks. This mask is obtained by dilating a GM mask extracted from the FreeSurfer’s *aseg* segmentation, and it ensures components are not extracted from voxels containing a minimal fraction of GM. Finally, these masks are resampled into BOLD space and binarized by thresholding at 0.99 (as in the original implementation). Components are also calculated separately within the WM and CSF masks. For each CompCor decomposition, the *k* components with the largest singular values are retained, such that the retained components’ time series are sufficient to explain 50 percent of variance across the nuisance mask (CSF, WM, combined, or temporal). The remaining components are dropped from consideration. The head-motion estimates calculated in the correction step were also placed within the corresponding confounds file. The confound time series derived from head motion estimates and global signals were expanded with the inclusion of temporal derivatives and quadratic terms for each ^18^. Frames that exceeded a threshold of 0.5 mm FD or 1.5 standardized DVARS were annotated as motion outliers. Additional nuisance timeseries are calculated by means of principal components analysis of the signal found within a thin band (*crown*) of voxels around the edge of the brain, as proposed by ^19^. The BOLD time-series were resampled onto the following surfaces (FreeSurfer reconstruction nomenclature): *fsnative*. All resamplings were performed with *a single interpolation step* by composing all the pertinent transformations (i.e. head-motion transform matrices, susceptibility distortion correction when available, and co-registrations to anatomical and output spaces). Gridded (volumetric) resamplings were performed using `nitransforms`, configured with cubic B-spline interpolation. Non-gridded (surface) resamplings were performed using `mri_vol2surf` (FreeSurfer).

**Supplementary Figure 11.**
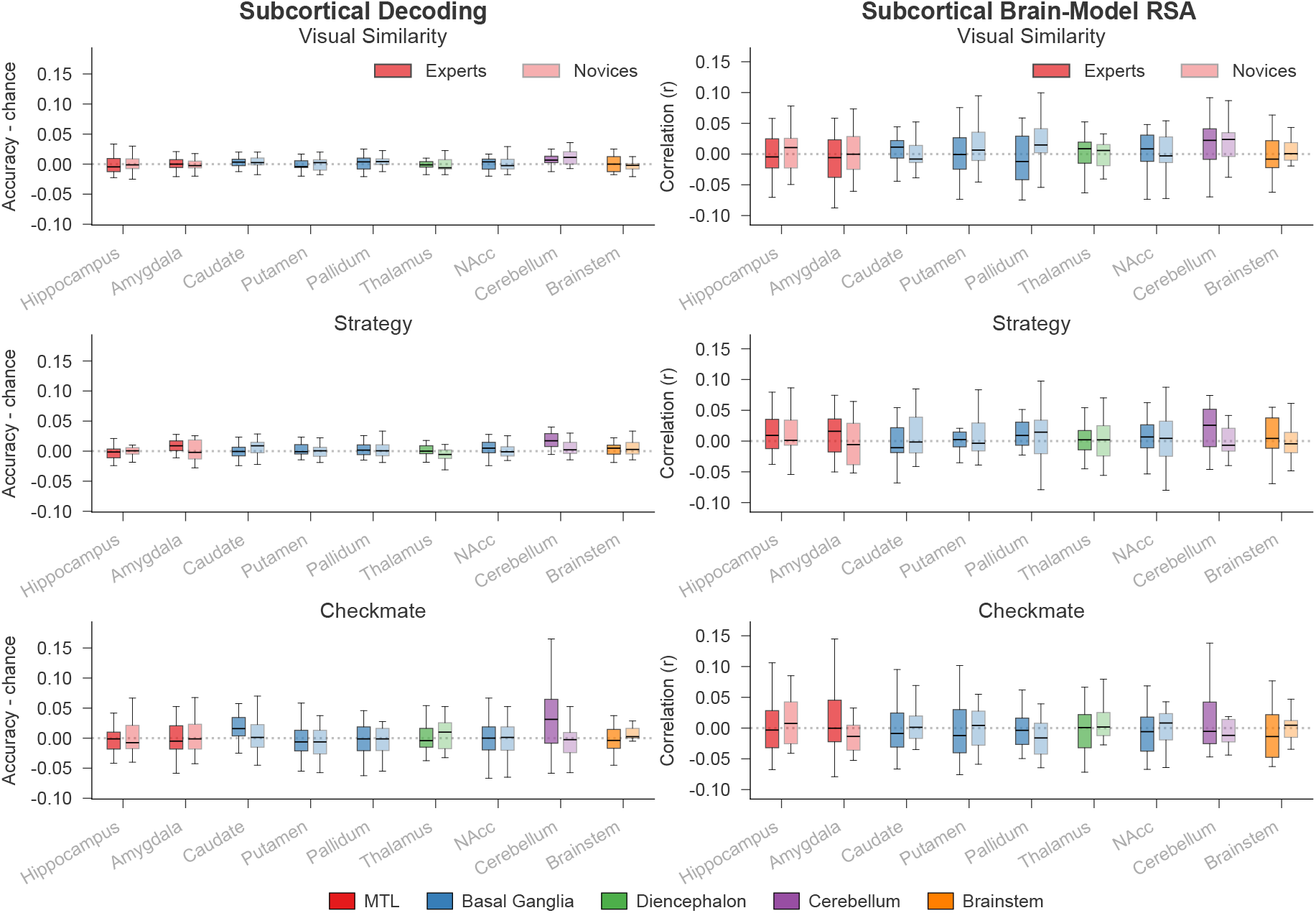
Subcortical decoding and RSA results. Left column: SVM decoding accuracy (minus chance) for Visual Similarity, Strategy, and Checkmate dimensions. Right column: RSA correlation (*r*) with the corresponding model RDMs. Box plots show the distribution of per-subject values for Experts (*n* = 20 independent participants, solid) and Novices (*n* = 20 independent participants, lighter shade). Boxes span the interquartile range (IQR; 25th–75th percentile), with the median marked by a horizontal line; whiskers extend to the furthest data point within 1.5 × IQR of the box edges. ROIs were derived from the CAB-NP subcortical atlas ^6^, grouped into 9 bilateral structures. Stars indicate two-sided Welch’s *t*-test (`scipy.stats.ttest_ind` with `equal_var=False`) Experts vs. Novices, FDR-corrected (Benjamini–Hochberg) across the 9 ROIs separately within each model: ^∗^ *p*_FDR_ *<* .05, ^∗∗^ *p*_FDR_ *<* .01, ^∗∗∗^ *p*_FDR_ *<* .001. No expert-novice difference survived FDR correction. Source data are provided as a Source Data file.

Many internal operations of *fMRIPrep* use *Nilearn* 0.10.4^20^ (RRID:SCR_001362), mostly within the functional processing workflow. For more details of the pipeline, see the section corresponding to workflows in *fMRIPrep*’s documentation.

**Supplementary Figure 12.**
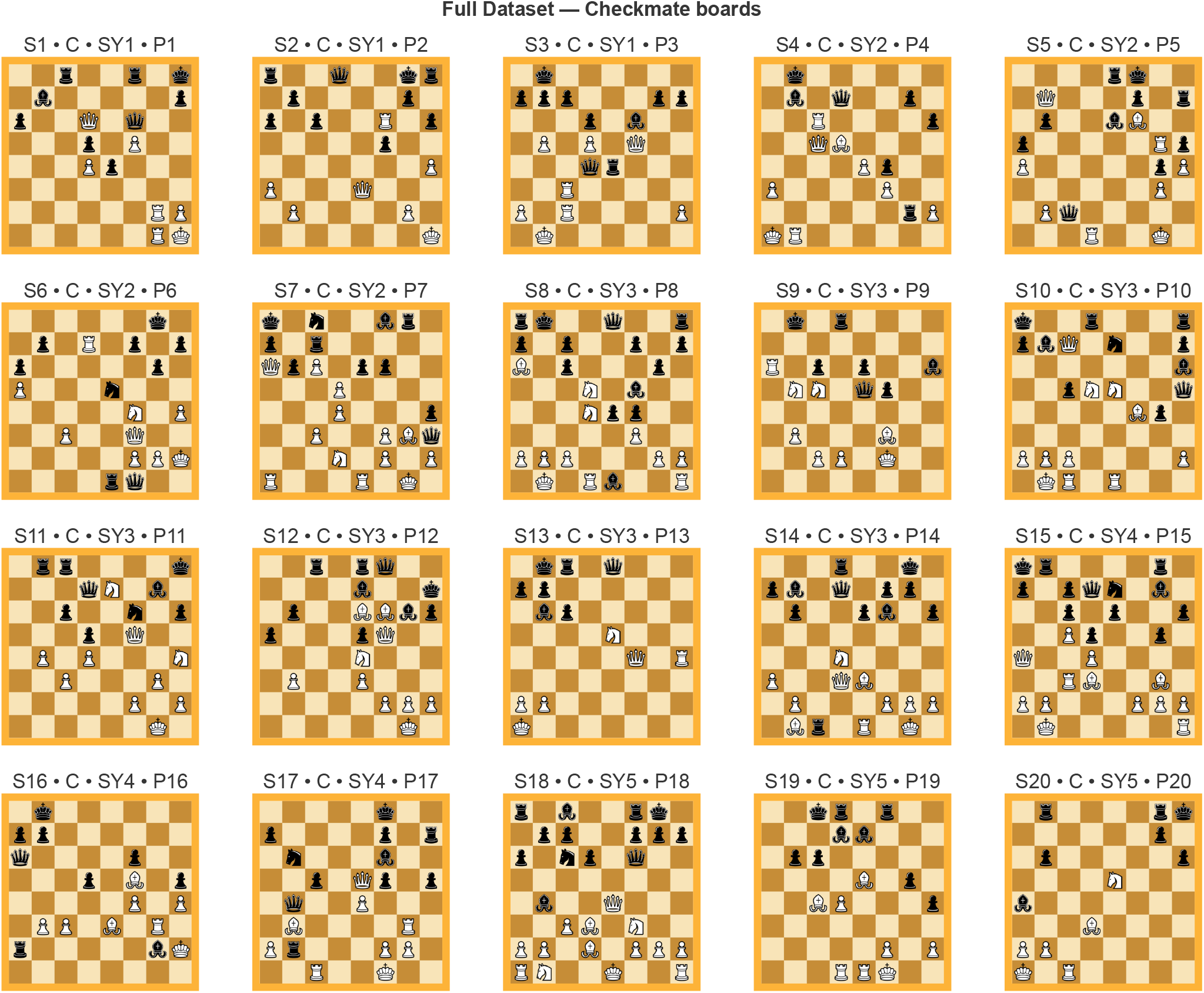
Full stimulus set used in the experiment — checkmate boards. Each chessboard represents one of the 20 unique checkmate stimuli from the dataset. The label above each board encodes: **S** (stimulus number), **C** for checkmate, **SY** (strategy index), and **P** (visual pair). Borders are shaded in orange tones, indicating checkmate status. The 20 non-checkmate counterparts are shown in Supplementary Fig. 13.

### S12 ROI analysis from uni and multivariate brain maps

To identify finer-grained anatomical regions showing significant differences in mean activation between Experts and Novices, we performed an exploratory second-level ROI analysis using the 180-region version of the Glasser parcellation described in Sec. 4.5. We conducted this analysis on two sets of brain maps: the univariate first-level GLM contrast maps (Sec. 4.6), and the searchlight RSA maps from Sec. 4.7.3.

To facilitate anatomical interpretation, each ROI was additionally assigned a secondary label from the Harvard-Oxford cortical atlas. This was done by computing the centre of mass of each ROI in MNI space and mapping the resulting coordinate to the nearest non-zero voxel in the 2 mm Harvard-Oxford atlas (thresholded at 25%). This combined approach allowed us to combine the spatial specificity of the Glasser atlas, which allows for more precise estimates of local activation patterns, with the broader and more familiar anatomical nomenclature of the Harvard-Oxford atlas.

#### Univariate contrast maps

For each participant and contrast (*Checkmate > Non-checkmate, All > Rest*), we imported the first-level contrast images and extracted mean signal values from each of the 180 cortical ROIs using the `NiftiLabelsMasker` implementation in `nilearn`. This yielded a single scalar value per ROI per participant, resulting in group-level matrices for Experts and Novices that were used for statistical comparison. For each ROI, we performed an independent-samples *t*-test comparing Experts and Novices, and computed the mean group difference, *t*-statistic, FDR-corrected *p*-value, and the 95% confidence interval of the difference. Results for significant ROIs are reported in Supplementary Table 13 and visualized in Supplementary Figure 14.

Univariate analyses were suggestive of several activation spots at or near locations that have been previously investigated in expertise literature, but in many cases these differences did not survive correction for multiple comparisons. This strengthens the suspicion that was the major motivation for our study: namely, that the classic univariate approach is not sensitive to the many ways in which expertise changes neural processing.

#### RSA searchlight maps

To facilitate anatomical interpretation of the searchlight RSA results, and to complement the univariate group-level analysis with a “content-oriented” approach, we applied the same second-level ROI-based procedure described in the above paragraph. For each participant theoretical model (Checkmate, Strategy, Visual Similarity), we imported the individual whole-brain correlation maps obtained from the searchlight (see Sec. 4.8), extracted the mean correlation value within each of the 180 Glasser ROIs, and performed independent-samples *t*-tests comparing Experts and Novices ROI-wise. Anatomical labels from the Harvard-Oxford cortical atlas were assigned to each ROI based on their centre of mass.

Results from this analysis (see Supplementary Tables 11 and 12, and Supplementary Figure 15) showed more extensive networks involved in chess expertise than the univariate counterpart. On the one hand, this suggests that multi-variate approaches may tap onto complementary aspects than univariate ones, and on the other that expertise effects in the brain may involve broader networks than previously thought.

**Supplementary Figure 13.**
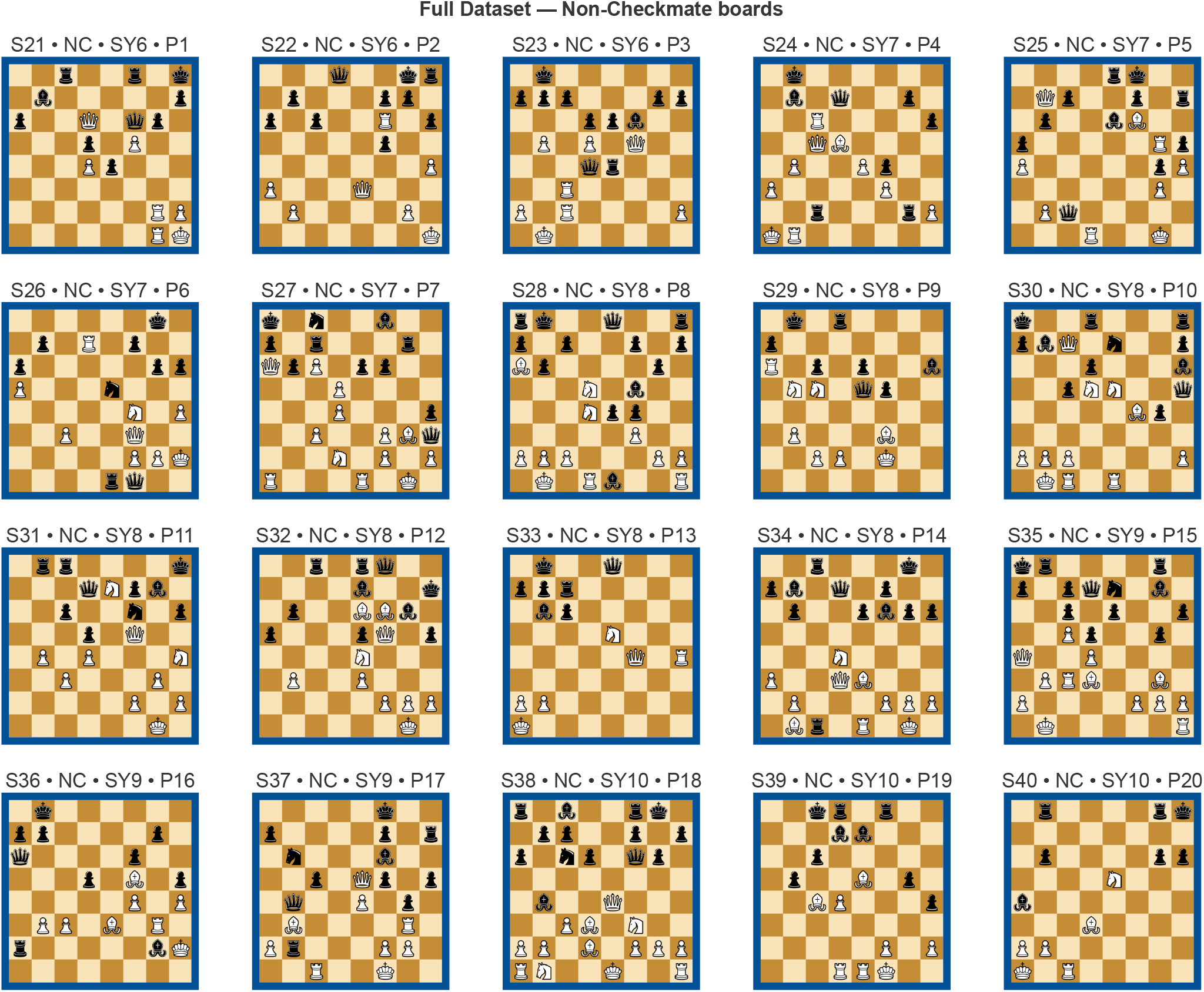
Full stimulus set used in the experiment — non-checkmate boards. Each chessboard represents one of the 20 unique non-checkmate stimuli from the dataset. The label above each board encodes: **S** (stimulus number), **NC** for non-checkmate, **SY** (strategy index), and **P** (visual pair). Borders are shaded in blue tones, indicating non-checkmate status. Each non-checkmate board pairs with the checkmate counterpart of the same visual pair P (see Supplementary Fig. 12).

**Supplementary Figure 14.**
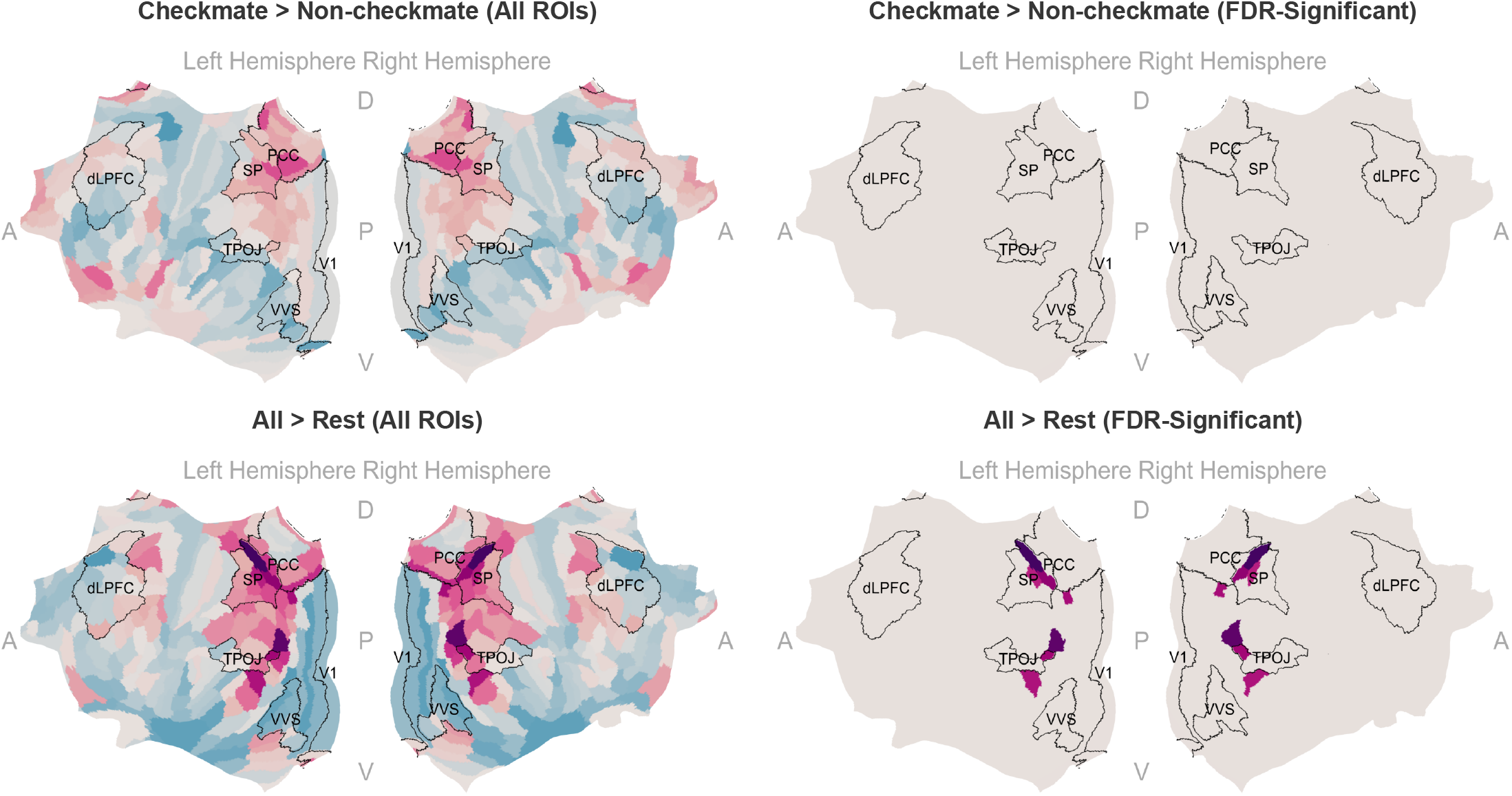
Univariate ROI differences (Experts vs. Novices). Cortical parcels from the 180-area Glasser atlas showing group differences in second-level GLM contrast maps (*Checkmate > Non-checkmate, All > Rest*) for *n* = 20 Experts and *n* = 20 Novices (independent participants in each group). Colours indicate the direction and relative magnitude of the Experts −Novices difference in mean contrast value (red: Experts > Novices; blue: Novices > Experts), saturated to the per-map maximum absolute value. Group differences were tested with two-sided independent-samples *t*-tests on per-subject mean contrast values within each parcel; only parcels surviving Benjamini–Hochberg FDR correction at *α* = 0.05 across the 180 parcels (within each contrast) are coloured. Per-parcel mean differences, *t*-statistics, raw and FDR-corrected *p*-values, and 95% confidence intervals are reported in Supplementary Table 13. Source data are provided as a Source Data file.

**Supplementary Figure 15.**
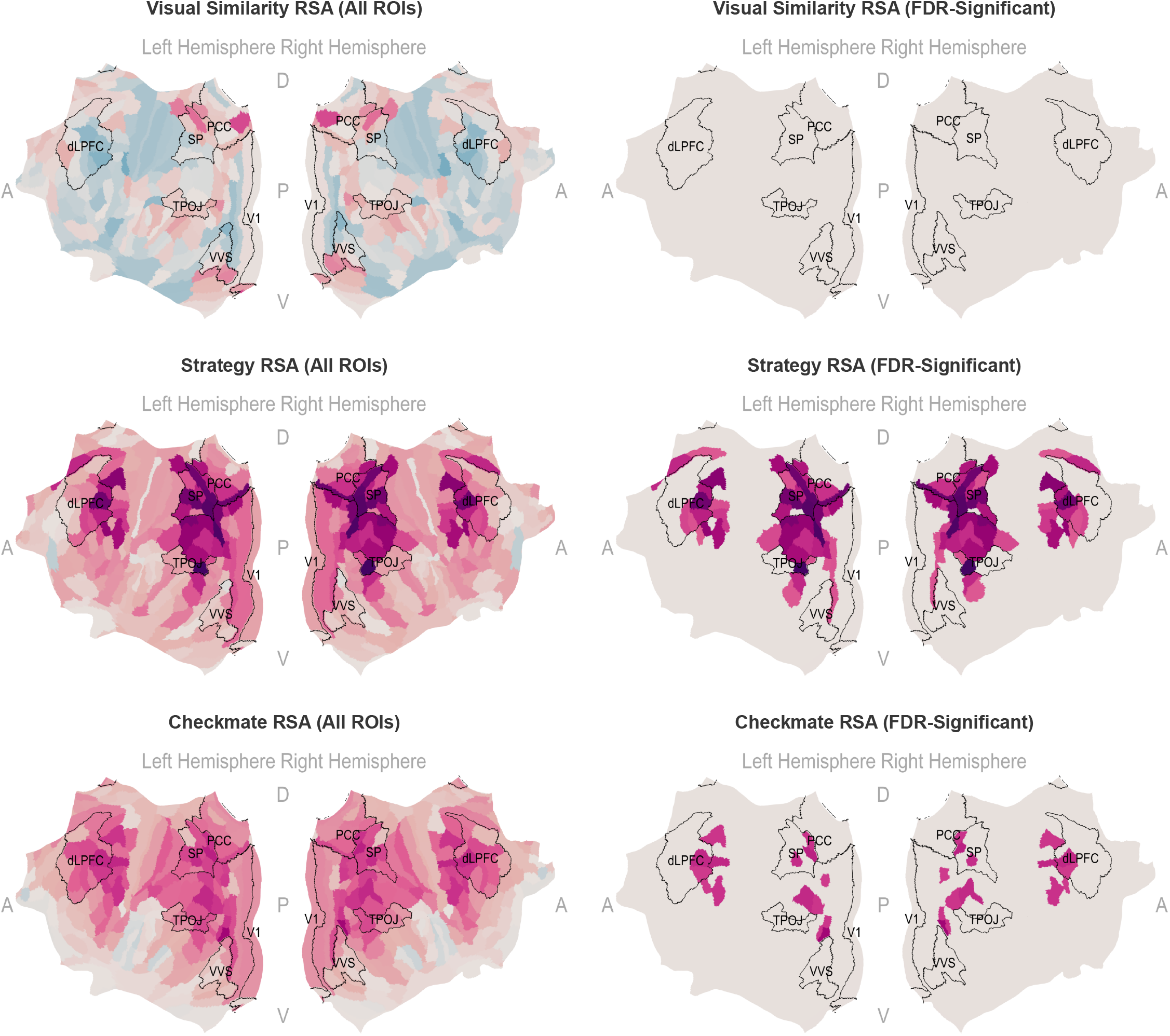
ROI summaries of searchlight RSA group differences. For each model RDM (Checkmate, Strategy, Visual similarity), whole-brain searchlight RSA maps were averaged within Glasser parcels to compare *n* = 20 Experts and *n* = 20 Novices (independent participants in each group). *Left*: uncorrected visualisation (mean Experts Novices difference per parcel; red: Experts *>* Novices; blue: Novices *>* Experts; saturated to the per-map maximum absolute value). *Right*: parcels surviving Benjamini–Hochberg FDR correction at *α* = 0.05 across the 180 parcels (within each model). Group differences per parcel were tested with two-sided independent-samples *t*-tests on per-subject mean RSA fit. Per-parcel mean differences, *t*-statistics, raw and FDR-corrected *p*-values, and 95% confidence intervals are reported in Supplementary Tables 11 and 12. Source data are provided as a Source Data file.

**Supplementary Figure 16.**
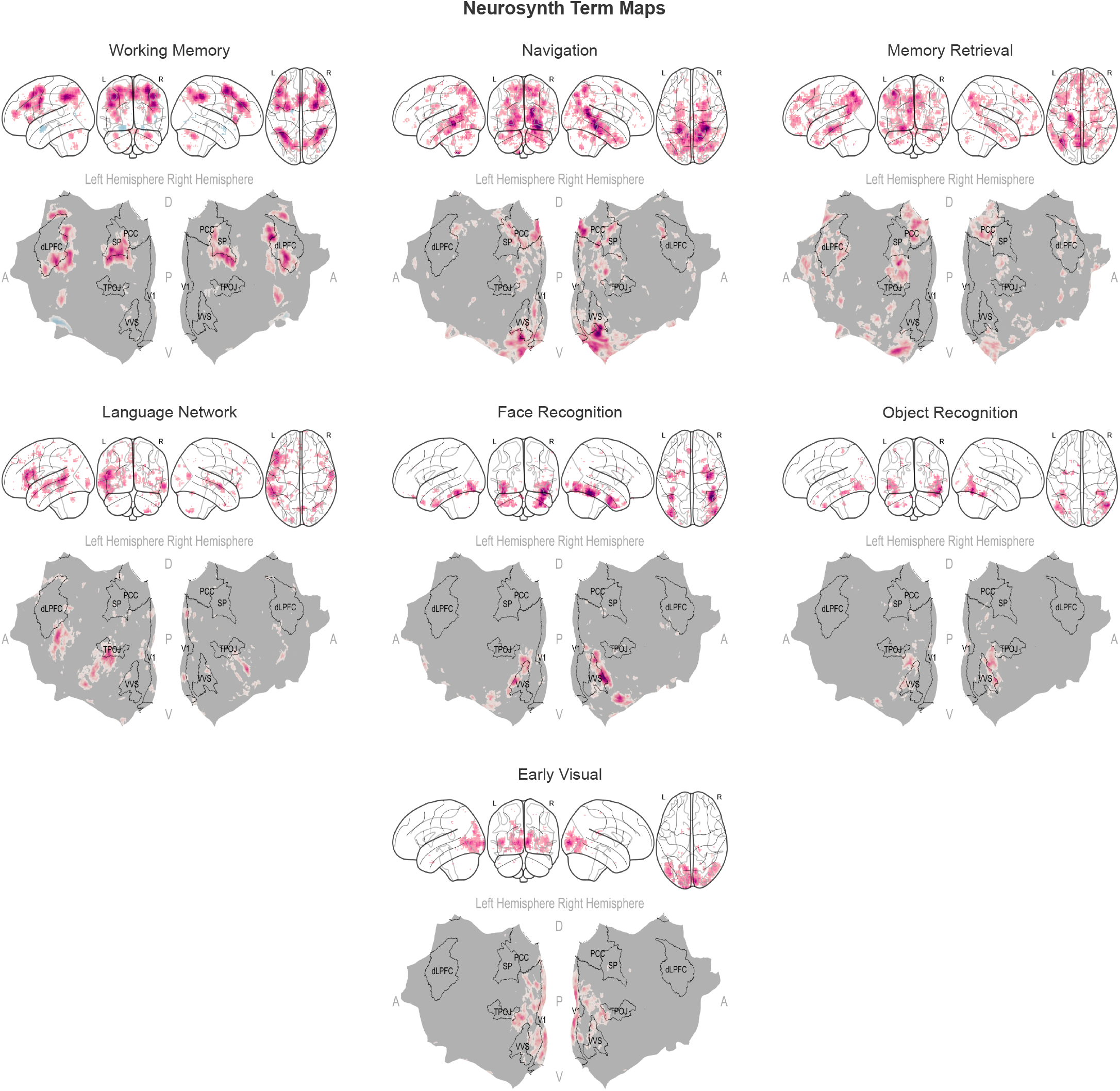
Neurosynth meta-analytic cognitive maps. The seven *a priori* terms (Working Memory, Memory Retrieval, Navigation, Language Network, Face Recognition, Object Recognition, Early Visual) used in Figs. 7 and 8 are shown in volume space along with the corresponding pial-surface projection. Each map is a Neurosynth association (*z*) map downloaded from https://neurosynth.org/ (accessed 2024-08; 14,371-study database) and resampled to the MNI152NLin2009cAsym template at 2 mm resolution. Display thresholds are the standard Neurosynth association threshold (*z >* 3.1). No statistical test is performed on this figure (the maps are reproduced for reference); inferential analyses combining these maps with our group-level *t*- and RSA-searchlight maps are reported in Figs. 7 and 8 and Supplementary Tables 10 and 9.

**Supplementary Table 1.** Complete listing of all 40 experimental stimuli with their Forsyth-Edwards Notation (FEN) strings. Each stimulus ID (S01–S40) corresponds to a unique chess board position used in the study.

| Stimulus ID | FEN Notation |
| --- | --- |
| S01 | 2r2r1k/1b5p/p2Q1q2/3p1P2/3Pp3/8/6RP/6RK w - - 0 1 |
| S02 | r2q2kr/1p4p1/p1p2R1p/5p2/7P/P3Q3/1P4P1/7K w - - 0 1 |
| S03 | 1k6/ppp3pp/3p1b2/1P1P1Q2/3qr3/2R5/P1R4P/1K6 w - - 0 1 |
| S04 | 1k6/1b1q2p1/2R4p/2QB4/4Pp2/P4P2/6rP/KR6 w - - 0 1 |
| S05 | 4rk2/1Q3p1r/1p2bB2/p5Rp/P5pP/6P1/1Pq5/3R2K1 w - - 0 1 |
| S06 | 6k1/1p1R1p1p/p5p1/P3n3/5N1P/2P2Q2/5PPK/4rq2 w - - 0 1 |
| S07 | k1n2br1/p1r5/QpP1pp2/3P4/3P3p/2P2PBq/3N1P1P/R3R1K1 w - - 0 1 |
| S08 | rk2q2r/p1p2p1p/B1p3p1/3N1b2/3Npp2/5P2/PPP3PP/1K1Rb2R w - - 0 1 |
| S09 | 1k1r4/8/R1p1p2b/1NN1qp2/8/1P3B2/2PP1K2/8 w - - 0 1 |
| S10 | k2r3r/pbQ1n2p/7b/2pNN2q/5Bp1/8/PPP4P/1KR1R3 w - - 0 1 |
| S11 | 1rr4k/3qN1b1/2p2n1p/3p1Q2/1P1P3N/2P3P1/5P1P/6K1 w - - 0 1 |
| S12 | 2r1rq2/4b2k/1p2BBbp/p3pQ2/4N3/1P2P3/5PPP/6K1 w - - 0 1 |
| S13 | 1kr1q3/pp6/1bp5/4N3/5Q1R/8/PP6/K7 w - - 0 1 |
| S14 | 3r2k1/pb1q1pp1/1p2pb1p/8/3N4/P2QB3/1P3PPP/1Br1R1K1 w - - 0 1 |
| S15 | kr4r1/p1pqn1b1/2p1p2p/2Pp2p1/Q2P4/2RB2B1/PP3PPP/1K5R w - - 0 1 |
| S16 | 1k6/pp6/q4p2/3p1B1p/5P1P/1PP1B1R1/r5bK/8 w - - 0 1 |
| S17 | 5k2/p4p1r/1n3b2/2p1Qp1p/1q2P3/1B4R1/Pr3PP1/2R2K2 w - - 0 1 |
| S18 | r1b2rk1/1pp2ppp/p1np1q2/8/1b2Q3/2PB1N2/PP1B1PPP/RN2K2R w - - 0 1 |
| S19 | 2kr1r2/3bb3/1pp5/4B1p1/2BP3p/8/5P1P/3RRK2 w - - 0 1 |
| S20 | 1r4rk/6p1/1p5p/4N3/b7/3B4/PP6/K1R5 w - - 0 1 |
| S21 | 2r2r1k/1b5p/p2Q1qp1/3p1P2/3Pp3/8/6RP/6RK w - - 0 1 |
| S22 | 3q2kr/1p3pp1/p1p2R1p/5p2/7P/P3Q3/1P4P1/7K w - - 0 1 |
| S23 | 1k6/ppp3pp/3ppb2/1P1P1Q2/3qr3/2R5/P1R4P/1K6 w - - 0 1 |
| S24 | 1k6/1b1q2p1/2R4p/2QB4/1P2Pp2/P4P2/2r3rP/KR6 w - - 0 1 |
| S25 | 4rk2/1Qp2p1r/1p2bB2/p5Rp/P5pP/6P1/1Pq5/3R2K1 w - - 0 1 |
| S26 | 6k1/1p1R1p2/p5pp/P3n3/5N1P/2P2Q2/5PPK/4rq2 w - - 0 1 |
| S27 | k1n2b2/p1r3r1/QpP1pp2/3P4/3P3p/2P2PBq/3N1P1P/R3R1K1 w - - 0 1 |
| S28 | rk2q2r/p1p2p1p/Bp4p1/3N1b2/3Npp2/5P2/PPP3PP/1K1Rb2R w - - 0 1 |
| S29 | 1k1r4/p7/R1p1p2b/1NN1qp2/8/1P3B2/2PP1K2/8 w - - 0 1 |
| S30 | k2r3r/pbQ1n2p/3p3b/2pNN2q/5Bp1/8/PPP4P/1KR1R3 w - - 0 1 |
| S31 | 1rr4k/3qNpb1/2p2n1p/3p1Q2/1P1P3N/2P3P1/5P1P/6K1 w - - 0 1 |
| S32 | 2r1rq2/4b2k/1p2BBb1/p3pQ1p/4N3/1P2P3/5PPP/6K1 w - - 0 1 |
| S33 | 1k2q3/ppr5/1bp5/4N3/5Q1R/8/PP6/K7 w - - 0 1 |
| S34 | 2r3k1/pb1q1p2/1p2pbpp/8/3N4/P2QB3/1P3PPP/1Br1R1K1 w - - 0 1 |
| S35 | kr4r1/p1pqn1b1/2p1p2p/2Pp2p1/Q2P4/1PRB2B1/P4PPP/1K5R w - - 0 1 |
| S36 | 1k6/pp4p1/q4p2/3p1B1p/5P1P/1PP1B1R1/r5bK/8 w - - 0 1 |
| S37 | 5k2/p4p1r/1n3b2/2p1Qp1p/1q2P1p1/1B4R1/Pr3PP1/2R2K2 w - - 0 1 |
| S38 | r1b2rk1/1pp2p1p/p1np1qp1/8/1b2Q3/2PB1N2/PP1B1PPP/RN2K2R w - - 0 1 |
| S39 | 2kr1r2/3bb3/2p5/1p2B1p1/2BP3p/8/5P1P/3RRK2 w - - 0 1 |
| S40 | 1r4rk/8/1p4pp/4N3/b7/3B4/PP6/K1R5 w - - 0 1 |

**Supplementary Table 2.** Behavioural-model RSA correlations (Experts vs Novices). Values are computed from count-normalised behavioural RDMs. Pearson correlations between the count-normalised group-level behavioural RDM and three categorical model RDMs, computed on the lower-triangle of the 40 × 40 RDM (*n* = 780 unique stimulus pairs per group). Group RDMs were aggregated across *n* = 19 Experts (one Expert excluded due to button-box malfunction during scanning) and *n* = 20 Novices (independent participants). 95% confidence intervals come from a 10,000-sample percentile bootstrap of stimulus pairs (`pingouin.corr`, two-sided). *p*-values are FDR-corrected (Benjamini–Hochberg) across the three models within each group at *α* = 0.05.

| Model | Experts |  |  |  | Novices |  |  |  |
| --- | --- | --- | --- | --- | --- | --- | --- | --- |
| | $r$ | 95% | $p$ | $p_{FDR}$ | $r$ | 95% | $p$ | $p_{FDR}$ |
| Checkmate | 0.733 | 0.700 to 0.760 | <.001 | <.001 | −0.036 | −0.110 to 0.030 | .320 | .479 |
| Visual Similarity | −0.116 | −0.180 to −0.050 | .001 | .001 | 0.049 | −0.020 to 0.120 | .174 | .479 |
| Strategy | 0.251 | 0.180 to 0.320 | <.001 | <.001 | 0.021 | −0.050 to 0.090 | .564 | .564 |

**Supplementary Table 3.** Pairwise RDM and marginal board-level split-half reliability. For pairwise analyses, *r*_*half*_ is the mean Spearman correlation between random half-sample estimates and *r* _*full*_ is the Spearman–Brown corrected mean reliability. Empirical two-sided *p*-values are computed from the resampled *r* _*full*_ distribution (1,000 random splits per group); 95% percentile CIs. Δ denotes Experts −Novices. *n* = 19 Experts (one Expert excluded due to button-box malfunction during scanning) and *n* = 20 Novices (independent participants).

| Analysis | Comparison | $r_{half}$ | $r_{full}$ | 95% CI ($r_{full}$) | $p$ |
| --- | --- | --- | --- | --- | --- |
| Pairwise RDM | Experts (within) | 0.299 | 0.459 | 0.392 to 0.515 | .002 |
| Pairwise RDM | Novices (within) | 0.022 | 0.040 | −0.069 to 0.151 | .488 |
| Pairwise RDM | Between groups | −0.019 | −0.040 | −0.178 to 0.077 | .589 |
| Pairwise RDM | $\Delta$ Experts – Novices | — | 0.419 | 0.288 to 0.536 | .002 |
| Marginal preference | Experts | 0.866 | 0.928 | 0.892 to 0.950 | .002 |
| Marginal preference | Novices | 0.411 | 0.565 | 0.175 to 0.808 | .014 |

**Supplementary Table 4.** ROI-level RSA. Expert–novice difference in correlation (Δ*r*) for Visual Similarity, Strategy, and Check vs Non-Check. Within-group correlations vs zero: one-sample *t*-test, one-tailed (`scipy.stats.ttest_lsamp(alternative=‘greater’)`). Between-group test (Experts vs Novices): Welch’s two-sample *t*-test, two-tailed (`scipy.stats.ttest_ind, equal_var=False`). 95% confidence interval on the mean difference: scipy `result.confidence_interval(0.95)` (Welch’s *t*-distribution, Welch–Satterthwaite df). *p*-values FDR-corrected (Benjamini–Hochberg) across 22 ROIs separately for each model and each test family at *α* = 0.05. *n* = 20 Experts and *n* = 20 Novices (independent participants).

| ROI | Visual Similarity |  |  |  | Strategy |  |  |  | Checkmate |  |  |  |
| --- | --- | --- | --- | --- | --- | --- | --- | --- | --- | --- | --- | --- |
| | $\Delta r$ | 95% CI | $p$ | $p_{FDR}$ | $\Delta r$ | 95% CI | $p$ | $p_{FDR}$ | $\Delta r$ | 95% CI | $p$ | $p_{FDR}$ |
| Primary Visual | −0.020 | −0.062 to 0.022 | .344 | .922 | .011 | −0.008 to 0.031 | .247 | .271 | .013 | −0.005 to 0.030 | .156 | .264 |
| Early Visual | −0.028 | −0.081 to 0.025 | .290 | .922 | .026 | 0.007 to 0.045 | .009 | .023 | .020 | 0.000 to 0.041 | .054 | .099 |
| Dorsal Stream Visual | .009 | −0.029 to 0.046 | .646 | .922 | .046 | 0.024 to 0.069 | <.001 | .001 | .047 | 0.009 to 0.085 | .017 | .047 |
| Ventral Stream Visual | −0.007 | −0.042 to 0.028 | .669 | .922 | .010 | −0.010 to 0.030 | .308 | .322 | .016 | −0.011 to 0.044 | .239 | .328 |
| MT+ Complex Visual | −0.006 | −0.039 to 0.027 | .723 | .922 | .037 | 0.011 to 0.062 | .007 | .021 | .050 | 0.020 to 0.079 | .002 | .014 |
| Somatosensory and Motor | −0.016 | −0.036 to 0.004 | .105 | .922 | .022 | −0.001 to 0.044 | .058 | .107 | .039 | 0.001 to 0.077 | .044 | .088 |
| Paracentral Lobular and Mid Cingulate | .002 | −0.022 to 0.026 | .875 | .922 | .035 | 0.009 to 0.062 | .009 | .023 | .048 | 0.005 to 0.090 | .030 | .074 |
| Premotor | −0.009 | −0.031 to 0.013 | .403 | .922 | .047 | 0.022 to 0.073 | <.001 | .004 | .074 | 0.031 to 0.118 | .002 | .014 |
| Posterior Opercular | .002 | −0.019 to 0.023 | .862 | .922 | .016 | −0.005 to 0.038 | .137 | .189 | .012 | −0.013 to 0.037 | .334 | .421 |
| Early Auditory | .008 | −0.012 to 0.028 | .412 | .922 | .013 | −0.007 to 0.034 | .194 | .237 | .007 | −0.016 to 0.030 | .535 | .558 |
| Auditory Association | .004 | −0.012 to 0.020 | .596 | .922 | .020 | −0.003 to 0.042 | .081 | .128 | .012 | −0.015 to 0.039 | .361 | .421 |
| Insular and Frontal Opercular | −0.007 | −0.031 to 0.017 | .548 | .922 | .020 | −0.002 to 0.042 | .078 | .128 | .010 | −0.017 to 0.038 | .450 | .495 |
| Medial Temporal | −0.002 | −0.022 to 0.019 | .876 | .922 | .015 | −0.008 to 0.038 | .192 | .237 | .009 | −0.021 to 0.039 | .558 | .558 |
| Lateral Temporal | .003 | −0.015 to 0.021 | .716 | .922 | .019 | −0.006 to 0.043 | .129 | .189 | .020 | −0.010 to 0.050 | .188 | .295 |
| Temporo-Parieto Occipital Junction | .008 | −0.015 to 0.031 | .505 | .922 | .043 | 0.018 to 0.068 | .001 | .005 | .056 | 0.018 to 0.094 | .006 | .024 |
| Superior Parietal | −0.001 | −0.032 to 0.029 | .922 | .922 | .070 | 0.044 to 0.096 | <.001 | <.001 | .076 | 0.029 to 0.122 | .003 | .015 |
| Inferior Parietal | −0.004 | −0.034 to 0.025 | .763 | .922 | .062 | 0.037 to 0.086 | <.001 | <.001 | .086 | 0.035 to 0.137 | .002 | .014 |
| Posterior Cingulate | .019 | −0.006 to 0.044 | .130 | .922 | .043 | 0.019 to 0.068 | <.001 | .004 | .072 | 0.020 to 0.125 | .009 | .031 |
| Anterior Cing. and Medial Prefrontal | .005 | −0.017 to 0.028 | .643 | .922 | .014 | −0.010 to 0.038 | .247 | .271 | .017 | −0.020 to 0.054 | .363 | .421 |
| Orbital and Polar Frontal | .002 | −0.021 to 0.024 | .894 | .922 | .002 | −0.022 to 0.025 | .877 | .877 | .019 | −0.011 to 0.049 | .214 | .313 |
| Inferior Frontal | −0.018 | −0.044 to 0.008 | .161 | .922 | .025 | 0.003 to 0.048 | .029 | .059 | .048 | 0.004 to 0.092 | .035 | .076 |
| Dorsolateral Prefrontal | −0.014 | −0.038 to 0.011 | .258 | .922 | .031 | 0.007 to 0.055 | .012 | .026 | .066 | 0.018 to 0.115 | .010 | .031 |

**Supplementary Table 5.**
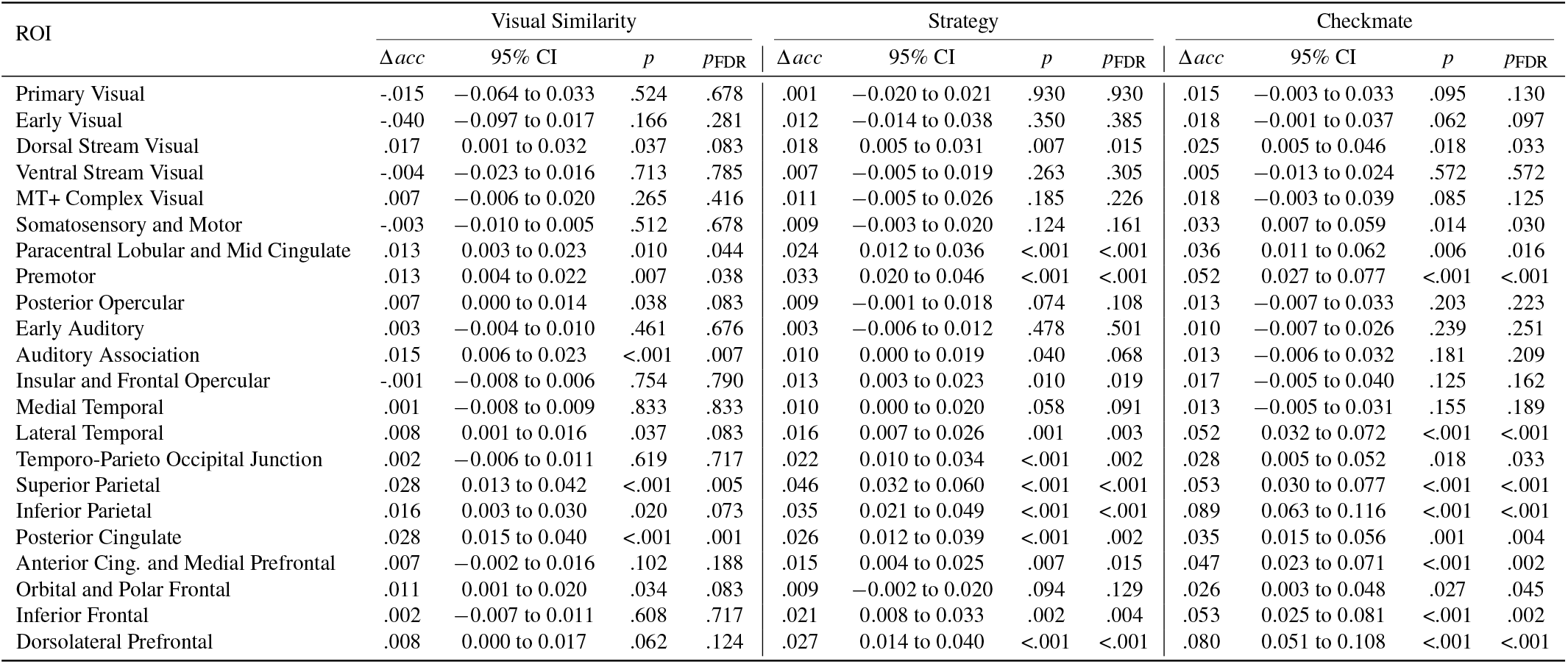
ROI-level decoding. Expert–novice difference in accuracy (Δ *acc*, chance-corrected) for Visual Similarity, Strategy, and Check vs Non-Check. Within-group test vs zero: one-sample *t*-test, one-tailed (`scipy.stats.ttest_lsamp(alternative=‘greater’)`). Between-group test: Welch’s two-sample *t*-test, two-tailed (`equal_var=False`). 95% CI on the mean difference: scipy `result.confidence_interval(0.95)`. *p*-values FDR-corrected (Benjamini–Hochberg) across 22 ROIs separately for each model and each test family at *α* = 0.05. *n* = 20 Experts and *n* = 20 Novices (independent participants).

**Supplementary Table 6.**
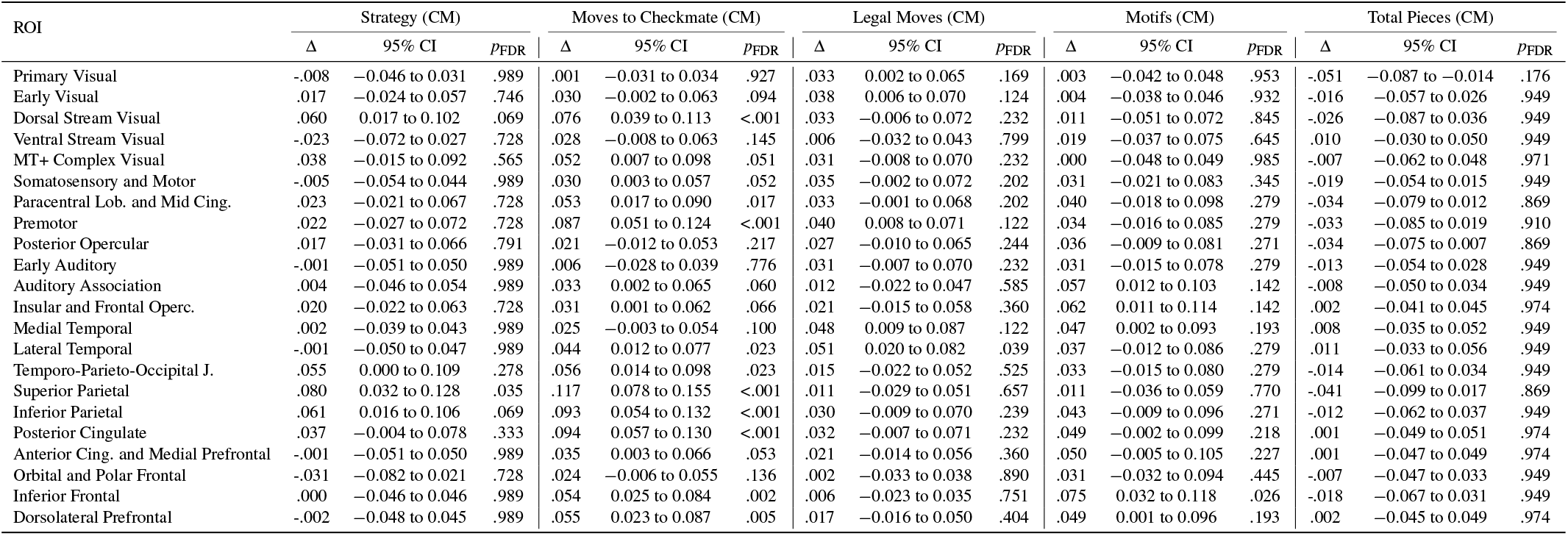
RSA – Experts–Novices difference across five finer regressors (Checkmate stimuli only). Restricted to the 20 checkmate boards. Within-group test vs zero: one-sample *t*-test, one-tailed (`alternative=‘greater’`). Between-group test: Welch’s two-sample *t*-test, two-tailed (`equal_var=False`). *p*-values FDR-corrected (Benjamini–Hochberg) across 22 ROIs separately for each regressor and each test family at *α* = 0.05. *n* = 20 Experts and *n* = 20 Novices (independent participants).

**Supplementary Table 7.**
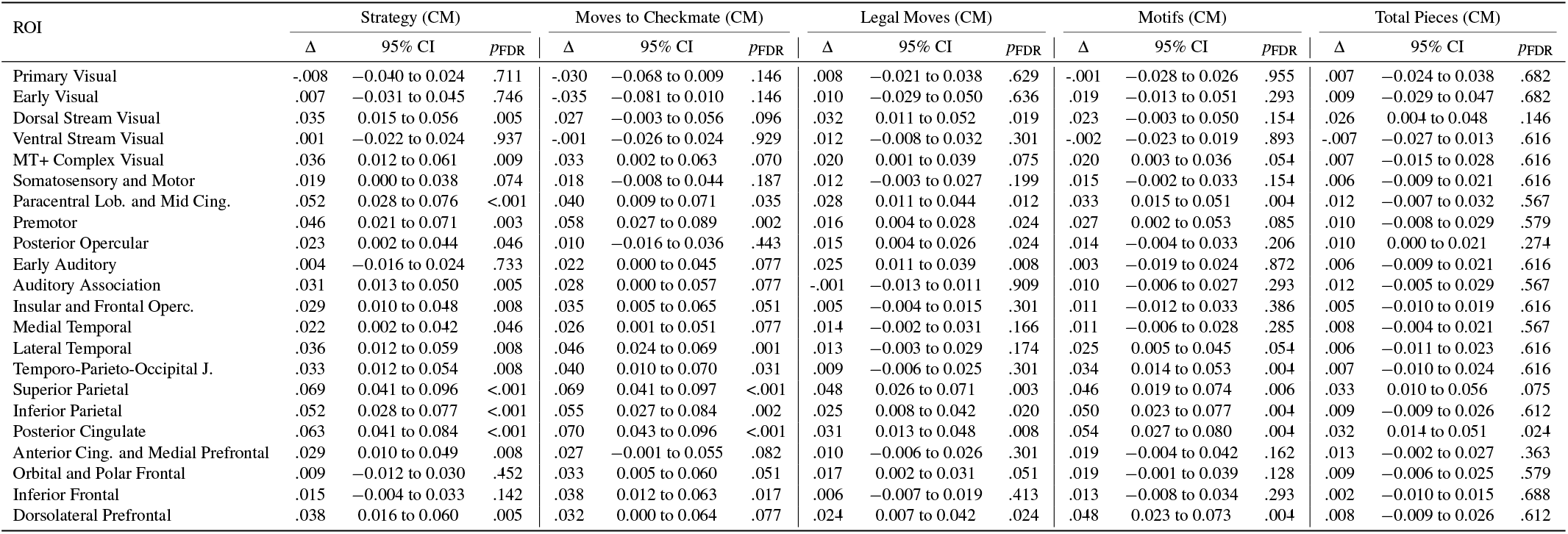
Decoding – Experts–Novices difference across five finer regressors (Checkmate stimuli only). Subject-level SVM cross-validated decoding accuracies (chance-corrected). Within-group test vs zero: one-sample *t*-test, one-tailed (`alternative=‘greater’`). Between-group test: Welch’s two-sample *t*-test, two-tailed (`equal_var=False`). *p*-values FDR-corrected (Benjamini–Hochberg) across 22 ROIs separately for each regressor and each test family at *α* = 0.05. *n* = 20 Experts and *n* = 20 Novices (independent participants).

**Supplementary Table 8.** Participation Ratio (PR) by ROI: group means with 95% confidence intervals, group differences (Experts −Novices) from Welch’s two-sample *t*-tests (scipy.stats.ttest_ind, equal_var=False; two-tailed) on per-subject PR values, with *t*-based 95% CIs (Welch–Satterthwaite df). Effect size: Cohen’s *d* (pooled SD). *p*-values FDR-corrected (Benjamini–Hochberg) across 22 ROIs at *α* = 0.05. *n* = 20 Experts and *n* = 20 Novices (independent participants).

| ROI | Experts |  | Novices |  | Experts – Novices |  |  |  |
| --- | --- | --- | --- | --- | --- | --- | --- | --- |
| | PR | CI95 | PR | CI95 | ΔPR | CI95 | *p* | *p*FDR |
| Primary Visual | 19.2 | 16.4 to 22.0 | 23.7 | 22.1 to 25.3 | -4.5 | -7.7 to -1.4 | .006 | .016 |
| Early Visual | 23.6 | 21.6 to 25.5 | 26.6 | 25.5 to 27.6 | -3.0 | -5.2 to -0.8 | .009 | .020 |
| Dorsal Stream Visual | 18.1 | 16.0 to 20.2 | 21.8 | 19.7 to 24.0 | -3.7 | -6.6 to -0.8 | .014 | .025 |
| Ventral Stream Visual | 25.8 | 23.5 to 28.0 | 29.7 | 28.1 to 31.4 | -3.9 | -6.6 to -1.2 | .006 | .016 |
| MT+ Complex Visual | 24.3 | 21.7 to 27.0 | 28.3 | 26.8 to 29.8 | -4.0 | -6.9 to -1.0 | .010 | .020 |
| Somatosensory and Motor | 27.7 | 25.0 to 30.4 | 29.1 | 27.1 to 31.0 | -1.3 | -4.6 to 1.9 | .407 | .471 |
| Paracentral Lob. and Mid Cing. | 27.8 | 25.4 to 30.2 | 30.5 | 29.1 to 31.8 | -2.7 | -5.3 to 0.0 | .051 | .074 |
| Premotor | 25.8 | 23.7 to 27.9 | 29.8 | 28.5 to 31.1 | -4.0 | -6.4 to -1.6 | .002 | .009 |
| Posterior Opercular | 23.0 | 21.2 to 24.8 | 23.3 | 21.6 to 25.0 | -0.3 | -2.6 to 2.1 | .808 | .846 |
| Early Auditory | 24.6 | 22.9 to 26.3 | 22.7 | 20.7 to 24.7 | 1.9 | -0.7 to 4.4 | .143 | .185 |
| Auditory Association | 27.4 | 25.4 to 29.4 | 27.1 | 25.1 to 29.1 | 0.3 | -2.4 to 3.0 | .846 | .846 |
| Insular and Frontal Operc. | 23.8 | 21.8 to 25.8 | 19.3 | 17.0 to 21.6 | 4.5 | 1.5 to 7.4 | .004 | .015 |
| Medial Temporal | 29.4 | 27.6 to 31.3 | 28.0 | 26.4 to 29.6 | 1.4 | -1.0 to 3.8 | .236 | .289 |
| Lateral Temporal | 28.5 | 26.4 to 30.5 | 29.4 | 27.6 to 31.3 | -1.0 | -3.6 to 1.7 | .475 | .522 |
| Temporo-Parieto-Occipital J. | 24.5 | 22.7 to 26.3 | 27.3 | 25.8 to 28.8 | -2.8 | -5.1 to -0.5 | .020 | .031 |
| Superior Parietal | 18.5 | 15.9 to 21.0 | 23.7 | 22.0 to 25.5 | -5.3 | -8.2 to -2.3 | .001 | .008 |
| Inferior Parietal | 22.5 | 20.5 to 24.5 | 25.2 | 24.1 to 26.4 | -2.8 | -5.0 to -0.5 | .019 | .031 |
| Posterior Cingulate | 24.9 | 22.9 to 26.8 | 28.4 | 27.2 to 29.5 | -3.5 | -5.7 to -1.3 | .003 | .012 |
| Anterior Cing. and Medial Prefrontal | 23.0 | 20.7 to 25.3 | 25.4 | 24.0 to 26.9 | -2.5 | -5.1 to 0.2 | .064 | .089 |
| Orbital and Polar Frontal | 20.1 | 17.0 to 23.1 | 25.6 | 23.0 to 28.2 | -5.6 | -9.5 to -1.7 | .007 | .016 |
| Inferior Frontal | 22.3 | 20.1 to 24.4 | 27.5 | 25.7 to 29.3 | -5.2 | -7.9 to -2.5 | <.001 | .005 |
| Dorsolateral Prefrontal | 22.0 | 19.6 to 24.3 | 27.7 | 26.1 to 29.2 | -5.7 | -8.5 to -2.9 | <.001 | .004 |

**Supplementary Table 9.** Neurosynth RSA correlations with 95% bootstrap confidence intervals. For each model RDM (Visual Similarity, Strategy, Checkmate), columns report Pearson correlations between group-level RSA searchlight difference maps (positive direction = Experts *>* Novices; negative direction = Novices *>* Experts) and seven Neurosynth meta-analytic term maps, computed across grey-matter voxels, plus their difference Δ = *r*_pos_ −*r*_neg_. Values are formatted as *r* [CI_low_, CI_high_]. 95% confidence intervals from a 10,000-sample percentile bootstrap of voxels (`pingouin.corr`, two-sided). *p*-values FDR-corrected (Benjamini–Hochberg) across 7 terms × 2 directions per model at *α* = 0.05. The underlying searchlight maps were derived from *n* = 20 Experts and *n* = 20 Novices (independent participants).

| Term | Visual Similarity |  |  | Strategy |  |  | Checkmate |  |  |
| --- | --- | --- | --- | --- | --- | --- | --- | --- | --- |
| | Pos+ | Neg- | $\Delta$ | Pos+ | Neg- | $\Delta$ | Pos+ | Neg- | $\Delta$ |
| Working Memory | -0.04 [-0.04, -0.03] | 0.04 [0.04, 0.05] | -0.08 [-0.09, -0.07] | 0.23 [0.22, 0.23] | -0.08 [-0.09, -0.08] | 0.31 [0.30, 0.32] | 0.22 [0.21, 0.22] | -0.09 [-0.09, -0.08] | 0.30 [0.30, 0.31] |
| Navigation | 0.15 [0.14, 0.15] | -0.08 [-0.08, -0.07] | 0.22 [0.21, 0.23] | 0.04 [0.03, 0.04] | -0.03 [-0.03, -0.02] | 0.07 [0.06, 0.08] | -0.00 [-0.01, 0.00] | -0.00 [-0.01, 0.00] | 0.00 [-0.01, 0.01] |
| Memory Retrieval | -0.01 [-0.01, 0.00] | 0.00 [-0.00, 0.01] | -0.01 [-0.02, 0.00] | 0.01 [0.01, 0.02] | -0.03 [-0.04, -0.03] | 0.04 [0.04, 0.05] | 0.04 [0.03, 0.04] | -0.02 [-0.02, -0.01] | 0.06 [0.05, 0.07] |
| Language Network | 0.02 [0.02, 0.03] | -0.02 [-0.03, -0.02] | 0.04 [0.03, 0.05] | 0.01 [0.00, 0.01] | -0.01 [-0.02, -0.01] | 0.02 [0.01, 0.03] | -0.01 [-0.01, -0.00] | 0.01 [0.00, 0.01] | -0.02 [-0.03, -0.01] |
| Object Recognition | -0.01 [-0.02, -0.01] | 0.02 [0.02, 0.03] | -0.04 [-0.05, -0.03] | -0.02 [-0.03, -0.02] | 0.02 [0.02, 0.03] | -0.05 [-0.06, -0.04] | -0.01 [-0.02, -0.01] | -0.01 [-0.02, -0.01] | -0.00 [-0.01, 0.01] |
| Face Recognition | -0.02 [-0.03, -0.02] | 0.04 [0.03, 0.04] | -0.06 [-0.07, -0.05] | -0.07 [-0.08, -0.07] | 0.03 [0.03, 0.04] | -0.11 [-0.11, -0.10] | -0.07 [-0.07, -0.06] | 0.03 [0.02, 0.03] | -0.10 [-0.11, -0.09] |
| Early Visual | 0.04 [0.03, 0.04] | 0.04 [0.03, 0.04] | 0.00 [-0.01, 0.01] | -0.02 [-0.02, -0.01] | 0.02 [0.02, 0.03] | -0.04 [-0.05, -0.03] | 0.02 [0.01, 0.02] | -0.00 [-0.01, 0.00] | 0.02 [0.01, 0.03] |

**Supplementary Table 10.** Neurosynth meta-analytic correlations with 95% bootstrap confidence intervals for two contrasts: All *>* Rest and Check vs Non-Check. Columns report Pearson correlations between group-level second-level GLM *z*-maps (positive direction = Experts *>* Novices; negative direction = Novices *>* Experts) and seven Neurosynth term maps, computed across grey-matter voxels, plus their difference Δ = *r*_pos_ ™ *r*_neg_. Values are formatted as *r* [CI_low_, CI_high_]. 95% confidence intervals from a 10,000-sample percentile bootstrap of voxels (`pingouin.corr`, two-sided). *p*-values FDR-corrected (Benjamini–Hochberg) across 7 terms × 2 directions per contrast at *α* = 0.05. The underlying *z*-maps were derived from second-level GLM contrasts on *n* = 20 Experts and *n* = 20 Novices (independent participants).

| Term | All > Rest |  |  | Check vs Non-Check |  |  |
| --- | --- | --- | --- | --- | --- | --- |
| | Pos+ | Neg- | $\Delta$ | Pos+ | Neg- | $\Delta$ |
| Working Memory | 0.16 [0.15, 0.16] | -0.13 [-0.14, -0.13] | 0.29 [0.28, 0.30] | 0.04 [0.03, 0.04] | -0.01 [-0.02, -0.01] | 0.05 [0.04, 0.06] |
| Navigation | 0.17 [0.17, 0.18] | -0.10 [-0.10, -0.09] | 0.27 [0.26, 0.28] | 0.03 [0.03, 0.04] | -0.02 [-0.03, -0.01] | 0.05 [0.04, 0.06] |
| Memory Retrieval | -0.01 [-0.01, -0.00] | -0.02 [-0.03, -0.02] | 0.02 [0.01, 0.03] | 0.09 [0.08, 0.10] | -0.07 [-0.08, -0.06] | 0.16 [0.15, 0.17] |
| Language Network | -0.04 [-0.05, -0.04] | 0.02 [0.02, 0.03] | -0.06 [-0.07, -0.05] | -0.06 [-0.07, -0.06] | 0.03 [0.02, 0.04] | -0.09 [-0.10, -0.08] |
| Object Recognition | -0.04 [-0.04, -0.03] | 0.03 [0.02, 0.04] | -0.07 [-0.08, -0.06] | -0.05 [-0.06, -0.05] | 0.06 [0.05, 0.06] | -0.11 [-0.12, -0.10] |
| Face Recognition | -0.08 [-0.08, -0.07] | 0.04 [0.03, 0.04] | -0.12 [-0.12, -0.11] | -0.08 [-0.08, -0.07] | 0.04 [0.03, 0.04] | -0.12 [-0.12, -0.11] |
| Early Visual | -0.06 [-0.07, -0.06] | 0.06 [0.05, 0.06] | -0.12 [-0.13, -0.11] | -0.03 [-0.03, -0.02] | 0.03 [0.03, 0.04] | -0.06 [-0.07, -0.05] |

**Supplementary Table 11.** Checkmate model (RSA): Experts > Novices. Glasser ROIs showing stronger model–brain correspondence in Experts than Novices. Columns: Glasser ROI label; Harvard–Oxford label; *M*_*diff*_ (Experts −Novices); *t*; 95% CI; raw *p*; FDR-corrected *p*. Per-parcel summaries from the searchlight RSA group-difference map (Experts −Novices) aggregated to the 360 fine-grained Glasser cortical parcels. Welch’s two-sample *t*-test (`scipy.stats.ttest_ind, equal_var=False`), two-tailed; *p*-values FDR-corrected (Benjamini–Hochberg) across 360 parcels at *α* = 0.05. *n* = 20 Experts and *n* = 20 Novices (independent participants).

| ROI | Harvard–Oxford Label | $M_{diff}$ | $t$ | 95% CI | $p$ | $p_{FDR}$ |
| --- | --- | --- | --- | --- | --- | --- |
| Area 55b | Juxtapositional Lobule Cortex (Suppl. Motor) | 0.027 | 3.351 | 0.011 to 0.043 | .002 | .048 |
| Seventh Visual Area | Cuneal Cortex | 0.027 | 3.067 | 0.009 to 0.045 | .004 | .048 |
| Area Lateral Occipital 1 | Intracalcarine Cortex | 0.020 | 3.163 | 0.007 to 0.032 | .003 | .048 |
| Area Lateral Occipital 2 | Intracalcarine Cortex | 0.014 | 3.797 | 0.007 to 0.022 | <.001 | .048 |
| Medial Area 7P | Precuneous Cortex | 0.030 | 3.172 | 0.011 to 0.049 | .003 | .048 |
| Lateral Area 7P | Precuneous Cortex | 0.038 | 3.424 | 0.016 to 0.061 | .001 | .048 |
| Area Lateral IntraParietal ventral | Precuneous Cortex | 0.037 | 3.274 | 0.014 to 0.060 | .002 | .048 |
| Area 8Av | ParaCing. Gyrus | 0.036 | 3.289 | 0.014 to 0.058 | .002 | .048 |
| Area 8C | Cing. Gyrus, anterior division | 0.032 | 3.064 | 0.011 to 0.054 | .004 | .048 |
| Rostral Area 6 | Unlabeled | 0.025 | 3.200 | 0.009 to 0.040 | .003 | .048 |
| Area IFJa | Cing. Gyrus, anterior division | 0.026 | 3.079 | 0.009 to 0.043 | .004 | .048 |
| Area 6 anterior | Juxtapositional Lobule Cortex (Suppl. Motor) | 0.027 | 3.137 | 0.010 to 0.045 | .003 | .048 |
| Area PGp | Cuneal Cortex | 0.029 | 3.134 | 0.010 to 0.048 | .003 | .048 |
| Area PGs | Precuneous Cortex | 0.030 | 3.295 | 0.012 to 0.049 | .002 | .048 |
| Area V4t | Lingual Gyrus | 0.018 | 3.162 | 0.006 to 0.030 | .003 | .048 |

**Supplementary Table 12.**
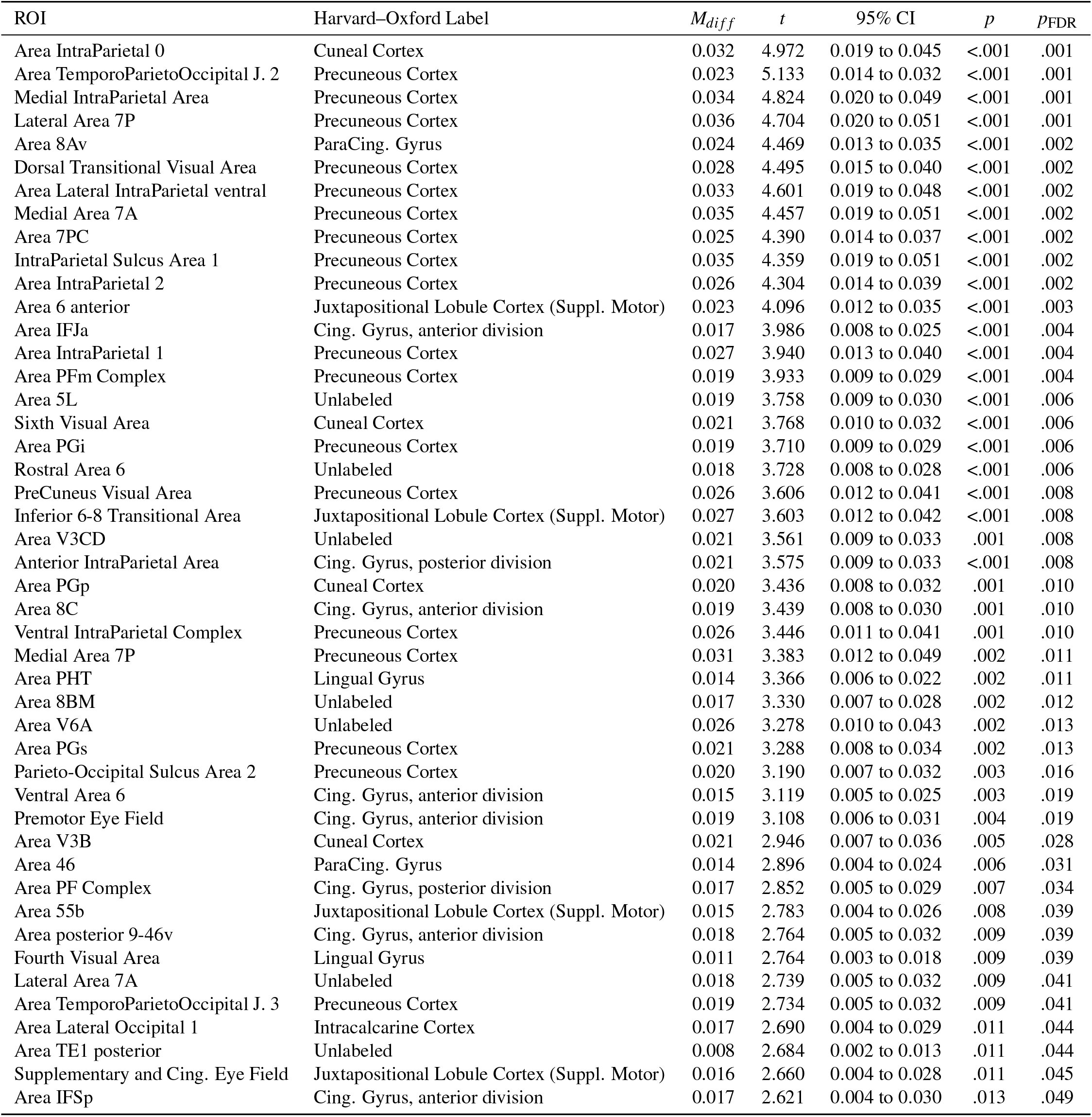
Strategy model (RSA): Experts > Novices. Glasser ROIs showing stronger model–brain correspondence in Experts than Novices. Columns: Glasser ROI label; Harvard–Oxford label; *M*_*diff*_ (Experts −Novices); *t*; 95% CI; raw *p*; FDR-corrected *p*. Per-parcel summaries from the searchlight RSA group-difference map (Experts −Novices) aggregated to the 360 fine-grained Glasser cortical parcels. Welch’s two-sample *t*-test (`scipy.stats.ttest_ind, equal_var=False`), two-tailed; *p*-values FDR-corrected (Benjamini–Hochberg) across 360 parcels at *α* = 0.05. *n* = 20 Experts and *n* = 20 Novices (independent participants).

**Supplementary Table 13.** Univariate contrast *All* > *Rest*: Experts > Novices. Glasser ROIs with higher second-level GLM contrast values in Experts than Novices. Columns show Glasser ROI label, Harvard–Oxford label, mean group difference *M*_*diff*_ (contrast units; Experts −Novices), *t* statistic, 95% CI, raw *p*, and FDR-corrected *p* value. Per-parcel summaries from the second-level GLM contrast aggregated to the 180-region Glasser cortical parcellation. Independent-samples Welch’s *t*-test (`scipy.stats.ttest_ind, equal_var=False`), two-tailed; *p*-values FDR-corrected (Benjamini–Hochberg) across 180 parcels at *α* = 0.05. *n* = 20 Experts and *n* = 20 Novices (independent participants).

| ROI | Harvard–Oxford Label | $M_{diff}$ | $t$ | 95% CI | $p$ | $p_{FDR}$ |
| --- | --- | --- | --- | --- | --- | --- |
| Medial Area 7A | Precuneous Cortex | 377.913 | 5.026 | 225.704 to 530.122 | <.001 | .002 |
| Area PGp | Cuneal Cortex | 306.595 | 4.626 | 172.423 to 440.766 | <.001 | .004 |
| Lateral Area 7P | Precuneous Cortex | 431.997 | 3.935 | 209.755 to 654.239 | <.001 | .021 |
| Area TemporoParietoOccipital J. 3 | Precuneous Cortex | 208.615 | 3.709 | 94.737 to 322.492 | <.001 | .029 |
| Area V6A | Unlabeled | 270.470 | 3.636 | 119.898 to 421.042 | <.001 | .029 |
| Area PHT | Lingual Gyrus | 146.853 | 3.543 | 62.935 to 230.770 | .001 | .032 |
| Ventral IntraParietal Complex | Precuneous Cortex | 304.701 | 3.381 | 122.234 to 487.169 | .002 | .043 |

**Supplementary Table 14.** Eyetracking decoding summary metrics for displacement and XY position features. Subject-level out-of-fold decoding accuracies from a linear SVM classifier (*n* = 40 independent participants combining Experts and Novices). Significance was assessed with a one-sample *t*-test against chance (0.5), two-tailed (`scipy.stats.ttest_lsamp`, default `alternative=‘two-sided’`). No multiple-comparison correction was applied (two pre-registered feature sets, reported separately). Wilson 95% CI on pooled run-level accuracy reported descriptively.

| Feature | Metric | Estimate | CI | $p$-value |
| --- | --- | --- | --- | --- |
| Displacement | Accuracy (subject mean, t-test) | 0.462 | 0.415 to 0.509 | .113 |
| Displacement | Accuracy (pooled runs, descriptive) | 0.468 | 0.417 to 0.520 |  |
| Displacement | Balanced Accuracy | 0.468 |  |  |
| Displacement | F1 | 0.468 |  |  |
| Displacement | ROC AUC | 0.099 |  |  |
| XY Position | Accuracy (subject mean, t-test) | 0.534 | 0.475 to 0.594 | .252 |
| XY Position | Accuracy (pooled runs, descriptive) | 0.532 | 0.480 to 0.583 |  |
| XY Position | Balanced Accuracy | 0.530 |  |  |
| XY Position | F1 | 0.532 |  |  |
| XY Position | ROC AUC | 0.457 |  |  |

**Supplementary Table 15.** Run-matched control: ROI-level RSA. Expert–novice difference in correlation (Δ*r*) for Visual Similarity, Strategy, and Checkmate. Statistical conventions identical to Supplementary Table 4: within-group test vs zero is a one-sample one-tailed *t*-test (`alternative=‘greater’`); between-group test is a two-sided Welch’s *t*-test (`equal_var=False`); 95% CI on the mean difference from scipy `result.confidence_interval(0.95)`. *p*-values FDR-corrected (Benjamini–Hochberg) across 22 ROIs separately within each model and each test family at *α* = 0.05. Both groups have identical run distributions (8 × 6 + 12 × 10 = 168 runs per group). *n* = 20 Experts and *n* = 20 Novices (independent participants).

| multirow2*ROI | Visual Similarity |  |  |  | Strategy |  |  |  | Check vs Non-Check |  |  |  |
| --- | --- | --- | --- | --- | --- | --- | --- | --- | --- | --- | --- | --- |
| | $\Delta r$ | 95% CI | $p$ | $p_{FDR}$ | $\Delta r$ | 95% CI | $p$ | $p_{FDR}$ | $\Delta r$ | 95% CI | $p$ | $p_{FDR}$ |
| Primary Visual | -0.005 | −0.047 to 0.037 | .825 | .957 | 0.011 | −0.009 to 0.031 | .294 | .308 | 0.014 | −0.004 to 0.031 | .122 | .168 |
| Early Visual | -0.013 | −0.063 to 0.038 | .609 | .957 | 0.028 | 0.008 to 0.048 | .007 | .016 | 0.020 | −0.001 to 0.041 | .064 | .118 |
| Dorsal Stream Visual | 0.020 | −0.017 to 0.057 | .287 | .957 | 0.048 | 0.026 to 0.071 | <.001 | <.001 | 0.046 | 0.008 to 0.084 | .019 | .038 |
| Ventral Stream Visual | 0.005 | −0.027 to 0.038 | .746 | .957 | 0.016 | −0.004 to 0.037 | .116 | .128 | 0.016 | −0.011 to 0.043 | .245 | .270 |
| MT+ Complex and Neighboring Visual Areas | 0.003 | −0.029 to 0.035 | .842 | .957 | 0.041 | 0.017 to 0.065 | .001 | .005 | 0.051 | 0.021 to 0.080 | .002 | .009 |
| Somatosensory and Motor | -0.016 | −0.036 to 0.005 | .127 | .935 | 0.024 | 0.001 to 0.047 | .039 | .057 | 0.049 | 0.012 to 0.086 | .012 | .029 |
| Paracentral Lobular and Mid Cingulate | 0.001 | −0.022 to 0.024 | .939 | .957 | 0.037 | 0.011 to 0.064 | .007 | .016 | 0.056 | 0.014 to 0.098 | .011 | .029 |
| Premotor | -0.007 | −0.029 to 0.014 | .482 | .957 | 0.048 | 0.023 to 0.073 | <.001 | .002 | 0.078 | 0.034 to 0.122 | .001 | .009 |
| Posterior Opercular | -0.004 | −0.026 to 0.017 | .684 | .957 | 0.022 | 0.001 to 0.044 | .041 | .057 | 0.021 | −0.003 to 0.045 | .087 | .127 |
| Early Auditory | 0.004 | −0.015 to 0.023 | .689 | .957 | 0.017 | −0.004 to 0.039 | .107 | .124 | 0.012 | −0.012 to 0.036 | .302 | .317 |
| Auditory Association | 0.000 | −0.016 to 0.017 | .957 | .957 | 0.026 | 0.002 to 0.050 | .032 | .051 | 0.018 | −0.009 to 0.044 | .184 | .224 |
| Insular and Frontal Opercular | -0.011 | −0.035 to 0.012 | .340 | .957 | 0.024 | 0.002 to 0.046 | .032 | .051 | 0.016 | −0.011 to 0.044 | .233 | .269 |
| Medial Temporal | -0.003 | −0.024 to 0.017 | .733 | .957 | 0.021 | −0.002 to 0.043 | .075 | .097 | 0.012 | −0.018 to 0.041 | .435 | .435 |
| Lateral Temporal | 0.001 | −0.016 to 0.018 | .939 | .957 | 0.026 | 0.004 to 0.049 | .025 | .045 | 0.027 | −0.004 to 0.057 | .082 | .127 |
| Temporo-Parieto Occipital Junction | 0.008 | −0.013 to 0.029 | .470 | .957 | 0.047 | 0.022 to 0.072 | <.001 | .002 | 0.061 | 0.022 to 0.099 | .003 | .014 |
| Superior Parietal | 0.005 | −0.024 to 0.035 | .709 | .957 | 0.075 | 0.049 to 0.101 | <.001 | <.001 | 0.080 | 0.034 to 0.126 | .002 | .009 |
| Inferior Parietal | 0.001 | −0.027 to 0.029 | .925 | .957 | 0.065 | 0.041 to 0.089 | <.001 | <.001 | 0.093 | 0.042 to 0.144 | .001 | .009 |
| Posterior Cingulate | 0.022 | −0.002 to 0.046 | .077 | .935 | 0.048 | 0.024 to 0.073 | <.001 | .002 | 0.075 | 0.023 to 0.128 | .007 | .022 |
| Anterior Cingulate and Medial Prefrontal | 0.004 | −0.018 to 0.026 | .729 | .957 | 0.020 | −0.003 to 0.042 | .085 | .104 | 0.025 | −0.011 to 0.062 | .169 | .219 |
| Orbital and Polar Frontal | 0.001 | −0.022 to 0.023 | .948 | .957 | 0.004 | −0.019 to 0.027 | .719 | .719 | 0.028 | −0.003 to 0.059 | .071 | .120 |
| Inferior Frontal | -0.019 | −0.044 to 0.006 | .124 | .935 | 0.029 | 0.010 to 0.049 | .004 | .012 | 0.056 | 0.011 to 0.100 | .016 | .035 |
| Dorsolateral Prefrontal | -0.013 | −0.037 to 0.010 | .259 | .957 | 0.029 | 0.006 to 0.053 | .016 | .031 | 0.074 | 0.025 to 0.123 | .005 | .017 |

**Supplementary Table 16.** Run-matched control: Participation Ratio (PR) by ROI. Group means with 95% confidence intervals, group differences (Experts − Novices) from Welch’s two-sample *t*-tests (`scipy.stats.ttest_ind, equal_var=False`; two-tailed). *p*-values FDR-corrected (Benjamini–Hochberg) across 22 ROIs at *α* = 0.05. Both groups have identical run distributions (8 × 6 + 12 × 10 = 168 runs per group). *n* = 20 Experts and *n* = 20 Novices (independent participants).

| multirow2*ROI | Experts |  | Novices |  | Experts – Novices |  | *p* | *p*<sub>FDR</sub> |
| --- | --- | --- | --- | --- | --- | --- | --- | --- |
|  | Expert | Expert | Novice | Novice | Delta | Delta |  |  |
| Primary Visual | 19.198 | 16.405 to 21.991 | 23.510 | 22.257 to 24.762 | -4.312 | -7.316 to -1.307 | .007 | .021 |
| Early Visual | 23.557 | 21.580 to 25.534 | 26.570 | 25.298 to 27.841 | -3.012 | -5.299 to -0.726 | .011 | .023 |
| Dorsal Stream Visual | 18.093 | 15.994 to 20.192 | 22.072 | 19.820 to 24.324 | -3.979 | -6.957 to -1.001 | .010 | .023 |
| Ventral Stream Visual | 25.794 | 23.542 to 28.047 | 29.307 | 27.516 to 31.098 | -3.513 | -6.301 to -0.725 | .015 | .025 |
| MT+ Complex Visual | 24.329 | 21.697 to 26.960 | 28.155 | 26.599 to 29.710 | -3.826 | -6.806 to -0.847 | .014 | .025 |
| Somatosensory and Motor | 27.705 | 24.966 to 30.443 | 28.885 | 26.957 to 30.813 | -1.180 | -4.431 to 2.071 | .466 | .514 |
| Paracentral Lob. and Mid Cing. | 27.796 | 25.412 to 30.179 | 30.073 | 28.713 to 31.433 | -2.277 | -4.954 to 0.400 | .093 | .120 |
| Premotor | 25.783 | 23.712 to 27.854 | 29.702 | 28.380 to 31.025 | -3.919 | -6.309 to -1.529 | .002 | .009 |
| Posterior Opercular | 23.016 | 21.241 to 24.790 | 22.785 | 21.182 to 24.388 | 0.230 | -2.083 to 2.544 | .841 | .881 |
| Early Auditory | 24.576 | 22.852 to 26.301 | 21.922 | 19.787 to 24.057 | 2.654 | -0.004 to 5.313 | .050 | .074 |
| Auditory Association | 27.396 | 25.432 to 29.361 | 26.436 | 24.530 to 28.343 | 0.960 | -1.688 to 3.607 | .467 | .514 |
| Insular and Frontal Operc. | 23.803 | 21.794 to 25.813 | 18.511 | 16.217 to 20.806 | 5.292 | 2.340 to 8.244 | <.001 | .005 |
| Medial Temporal | 29.432 | 27.579 to 31.284 | 27.241 | 25.660 to 28.821 | 2.191 | -0.166 to 4.548 | .068 | .093 |
| Lateral Temporal | 28.473 | 26.413 to 30.533 | 28.655 | 26.721 to 30.590 | -0.183 | -2.916 to 2.551 | .893 | .893 |
| Temporo-Parieto-Occipital J. | 24.520 | 22.707 to 26.333 | 27.092 | 25.560 to 28.623 | -2.571 | -4.869 to -0.274 | .029 | .046 |
| Superior Parietal | 18.459 | 15.937 to 20.980 | 24.280 | 22.529 to 26.032 | -5.822 | -8.803 to -2.840 | <.001 | .004 |
| Inferior Parietal | 22.477 | 20.460 to 24.494 | 25.528 | 24.403 to 26.653 | -3.050 | -5.305 to -0.796 | .010 | .023 |
| Posterior Cingulate | 24.881 | 22.948 to 26.813 | 28.158 | 27.036 to 29.281 | -3.278 | -5.457 to -1.099 | .004 | .016 |
| Anterior Cingulate and Medial Prefrontal | 22.972 | 20.682 to 25.262 | 24.895 | 23.605 to 26.186 | -1.923 | -4.488 to 0.642 | .136 | .166 |
| Orbital and Polar Frontal | 20.054 | 16.960 to 23.148 | 25.278 | 22.609 to 27.947 | -5.224 | -9.179 to -1.269 | .011 | .023 |
| Inferior Frontal | 22.257 | 20.081 to 24.433 | 27.199 | 25.389 to 29.010 | -4.942 | -7.683 to -2.202 | <.001 | .005 |
| Dorsolateral Prefrontal | 21.965 | 19.584 to 24.346 | 27.636 | 26.069 to 29.202 | -5.670 | -8.441 to -2.899 | <.001 | .004 |

## Notes

https://github.com/costantinoai/chess-expertise-2025

### Competing Interest Statement

The authors have declared no competing interest.

### Summary of Updates

Filled in some missing references to sections: in the main text, many "Sec. " were left empty (now, e.g., "Sec. 4.8").

